# Novel measures of Morris water maze performance that use vector field maps to assess accuracy, uncertainty, and intention of navigational searches

**DOI:** 10.1101/2021.07.04.451032

**Authors:** P. Meenakshi, D. Mehrotra, N. Nruthyathi, D. Almeida-Filho, Y.-S. Lee, A.J. Silva, J. Balaji

## Abstract

Most commonly used behavioural measures for testing learning and memory in the Morris water maze (MWM) involve comparisons of an animal’s residence time in different quadrants of the pool. Such measures are limited in their ability to test different aspects of the animal’s performance. Here, we describe novel measures of performance in the MWM that use vector fields to capture the motion of mice as well as their search pattern in the maze. Using these vector fields, we develop quantitative measures of performance that are intuitive and more sensitive than classical measures. First, we describe search patterns in terms of vector field properties and use these properties to define three metrics of spatial memory namely Spatial Accuracy, Uncertainty and, Intensity of Search. We demonstrate the usefulness of these measures using four different data sets including comparisons between different strains of mice, an analysis of two mouse models of Noonan syndrome (*Ptpn11* D61G and *Ptpn11* N308D/+), and a study of goal reversal training. Importantly, besides highlighting novel aspects of performance in this widely used spatial task, our measures were able to uncover previously undetected differences, including in an animal model of Noonan syndrome, which we rescued with the mitogen activated protein kinase kinase (MEK) inhibitor SL327. Thus, our results show that our approach breaks down performance in the Morris water maze into sensitive measurable independent components that highlight differences in spatial learning and memory in the MWM that were undetected by conventional measures.

## Introduction

The Morris water maze task (MWM) and its modifications have been extensively used in the field of neuroscience to study a variety of cognitive phenomena, especially related to hippocampal-dependent learning and memory (Brandeis et al., 1989; D’Hooge & De Deyn, 2001; Redish & Touretzky, 1998). In the original or reference version of the MWM task, rodents, over multiple trials, learn the spatial location of a hidden platform submerged in a pool of water using distal cues (R. Morris, 1984; R. G. M. Morris, 1981). Learning and memory is assessed with multiple performance measures, such as time to the platform during training, and proximity during probe trials given at different times during training. During a probe trial, the platform is removed, and an animal is allowed to search for the platform usually for 60s. Search behaviour during a probe trial is typically the most sensitive measure of an animal’s spatial learning and memory. Traditionally, learning and memory is usually assessed by comparing the time the animal spent in the quadrant of the pool where the platform was located during training (target quadrant) with the time spent in the other three quadrants (Petrosini et al., 1998). Time spent in zones or areas of multiple sizes centered in the platform’s location is also used as a parameter for measuring spatial learning and memory as this provides a more detailed readout for the spatial distribution of the animal’s searches. Similarly, platform crossings, which describe the number of times the animal crossed the platform location during a probe trial, is also used as a metric for spatial memory. Additionally, path length is a metric that measures the total distance covered in searching for the platform during a probe trial (Dalm et al., 2000). Other conventional measures include escape latency, the time taken to reach the platform location, thigmotaxis duration (Wolfer & Lipp, 1992), i.e., the time the animal spends near the wall, and average speed, an indicator of swimming or motor skills. While these measures provide information on the animal’s efforts to locate the platform, they do not capture important aspects of the animal’s navigational performance.

Additional methods to measure the animal’s performance have been proposed over the years (Pereira & Burwell, 2015), including a platform proximity measure (Gallagher et al., 1993). This approach calculates the average distance of the rodent from the center of the platform location, and it is effective in differentiating between search strategies of equal path length (i.e., a focused search near the platform and a diffused search around the platform). Incidentally, this proximity measure has been shown to be more sensitive than classical measures described above, including quadrant search measures, zonal and platform crossings (Maei, Zaslavsky, Teixeira, et al., 2009).

Another approach to quantify performance in the MWM uses more complex methods that classify the animal’s swim trajectories into different path strategies. These approaches capture qualitative measures of performance in the water maze task (Garthe et al., 2009; Gehring et al., 2015; Graziano et al., 2003). Path strategy-based methods use semi-automated or machine learning algorithms to classify swim trajectories. Interestingly, such methods can detect multiple search strategies within a given MWM session (Cooke et al., 2019; Vouros et al., 2018). A parameter free machine learning based algorithm was also developed to classify MWM search strategies (Overall et al., 2020). This approach is automated and uses trajectories to assign performances in individual trials to different classes. These methods are dependent on a defined set of possible strategies that have been previously identified in available datasets used to train the classifiers. Thus, they are useful to develop descriptive accounts of an animal’s performance with defined search categories.

A measure was also proposed that uses the concept of entropy from information theory (Maei, Zaslavsky, Wang, et al., 2009). This entropy-based measure (measure H) describes the extent of search at the platform location as well as the focus of this search (Maei, Zaslavsky, Wang, et al., 2009). The rational for this approach is that rodents improve the focus or “organize” their search efforts to locate the platform as a function of learning. Thus, with training swim trajectories go from a highly disorganized to a more organized state. Computing the entropy of the swim trajectories (summation of entropy in path variance (H_path_) and entropy in error variance (H_error_) provides a more sensitive measure than the simpler proximity measure. However, the improved sensitivity of the H measure is dependent on the relative weighting of its components (H_path_ and H_error_) in a given experimental setting, since one individual component maybe be more sensitive to specific search strategies. Such non-linear estimates, despite being highly sensitive, do not provide a uniform scale of measurement that is intuitive and easy to use. Additionally, the H measure primarily estimates the disorderliness of performances in the MWM and does not reflect specific spatial or navigational properties of these performances.

Here, we propose new measures based on vector fields constructed from velocity components that are oriented towards a specific point in the water maze. Our measures stem from the observation that animals tend to slow down at and around the platform location during probe trials with the expectation of landing on the platform. This behaviour gains prominence as animals learn the location of the platform. Such slowing down also occurs when they are approaching the wall of the pool. In addition, animals show similar slowing down when they overshoot the platform position and turn around. Thus, the measures described here to understand the properties of these vectors and their spatial distribution as a vector field could provide valuable information regarding how the animal perceives the location of the platform. We proposed that the framework and associated measures introduced here provide a much-improved strategy to access the navigation performance of animals in the MWM.

The measures introduced here enabled us to identify subtle memory deficits in the MWM. We propose that vector maps of the velocity vector component along the occupancy center(*V*_‖*t*_) and its orthogonal component (*V*_⊥*t*_), contain information that reflects the intention of the animal’s movement. Specifically, we propose that the velocity vector component along the occupancy center (*V*_‖*t*_) measures the animal’s movement that contributes to its approach to the platform location. In addition, the velocity vector component orthogonal to the occupancy center (*V*_⊥*t*_) measures the animal’s movement that contributes to motion around the occupancy center. We also calculate the vector field properties, namely divergence on *V*_‖*t*_, to describe the rate of change of these measures in 2D space. Specifically, we propose that spatially localized negative divergence peaks of *V*_‖*t*_ reveal convergence hotspots or putative platform search centers (P_cs_). Using the search center measure, we quantify spatial memory in terms of three independent metrics, namely accuracy, uncertainty, and intensity of search.

We argue that unlike the previously proposed measures, the three metrics introduced here, based on divergence field properties, capture the nature of spatial memory independently of factors unrelated to learning and memory-based phenomena, such as motor performance.

### Theory

We have used velocity-based vector fields to develop an approach that captures key components of an animals spatial learning and memory performance in the Morris water maze (MWM). This approach depends on three independent measures, namely accuracy, uncertainty and intensity of search. Briefly, the approach starts by identifying a place in the pool where the animal is most likely to be found given its trajectory and term it “occupancy center”(P_oc_). After this, we construct a vector projection that measures the contribution of the animal’s movement towards the occupancy center as follows,

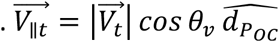

where 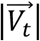 is the magnitude of the velocity vector and *θ*_*v*_ is the angle between the velocity vector 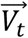 and a unit vector pointing towards the occupancy center 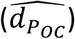 from the current position.

Then, we measure the divergence in such vector fields, and identify the peaks (extrema’s) in the divergence. We call these peaks putative search centers. From the putative search centers, we calculate the accuracy, uncertainity and intensity of search as follows

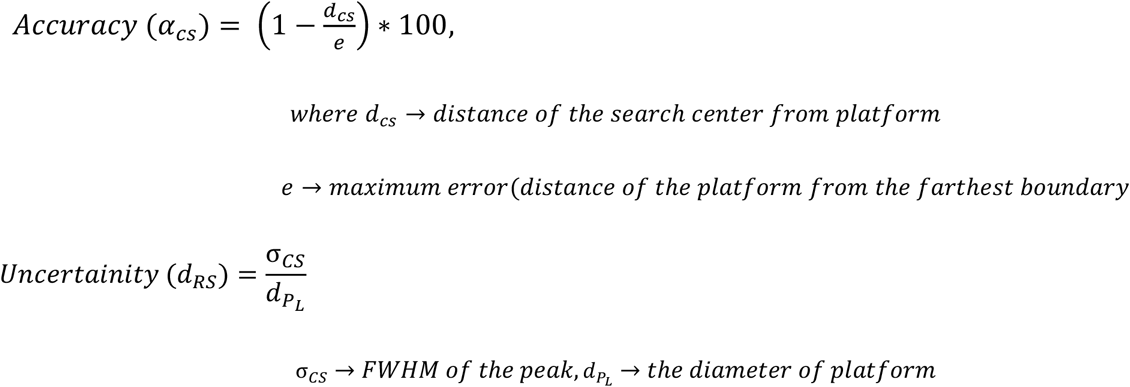

Additionally, we define absolute and relative intensity of search as follows:

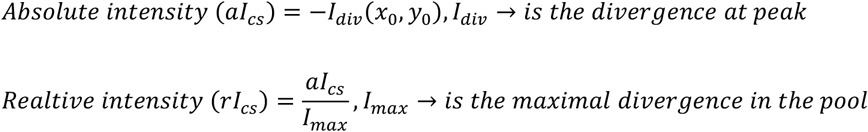

For a more complete explanation of our methods, please see Appendix: Theory.

## Results

### Swim trajectories described as velocity vector fields reveal that the speed of intentional movement varies over the pool space as a function of training

In an effort to assess spatial learning and memory, we first sought to describe the movement of the mice using the most basic kinematic measure, i.e., velocity. For this we used a previously published water maze data set from a mouse model of Noonan syndrome (Romano et al., 2010; Tartaglia & Gelb, 2005), the *Ptpn11^D61G/+^* mice and their littermate controls (Lee et al., 2014) .Using a time series of position data obtained from swim trajectories, we constructed velocity vectors (Eq. iii, Fig. 1C). These vectors describe the distance covered per unit time as the mice searched for the platform (Fig.2(A-D)). When we measured the magnitude of the velocity vector (speed), as a function of the radial distance from the platform, we noted that it was invariant across space when the mice had no knowledge of the platform’s location (Fig.3(A)). However, we observed that after training, there was a reduction in the speed of the mice closer to the platform location during the probe trials (Red Arrows in Fig.3(B), (C), (D)). There was also a reduction in speed of the mice when they were closer to the wall of the pool. We reasoned that studying the variation of speed across the spatial dimension would reveal the regions where the mice slowed down in their effort to locate the platform. Specifically, we propose that the divergence on the velocity vector field reveals the “sources” and “sinks” in that field, i.e., the regions where the mice swim away from with high speed (divergence points), and regions where the mice swim into with low speed (convergence points). In general, we recognise that at any given time, only a fraction of an animal’s movement, and not necessarily the entire movement, may be oriented towards its intended direction. Given this, we generated a vector field around the point in the pool where the mice are most likely to be found, i.e., the occupancy center. We propose that the regions of low residence typically result from incidental occupancy of these locations merely by being on the navigational path, rather than being the locations where the mice intent or decide to go. We identified the high occupancy regions as the regions where the mice intended to stay. We used maximum entropy(ME) to demarcate these regions as the regions where the mice are more likely to sample. We calculated the center of mass of these most likely occupied regions to get the occupancy center as shown in Fig2 (E-H). We carried out this analysis for wild type (WT) littermates of *Ptpn11^D61G/+^* mice on training day 1, as well as during probe trials given at the end of days 3, 5 and 7 of training. With this occupancy center, we described the swim trajectories of the mice in terms of a velocity vector component along the occupancy center (Eq. (vi)).

**Figure 1:**
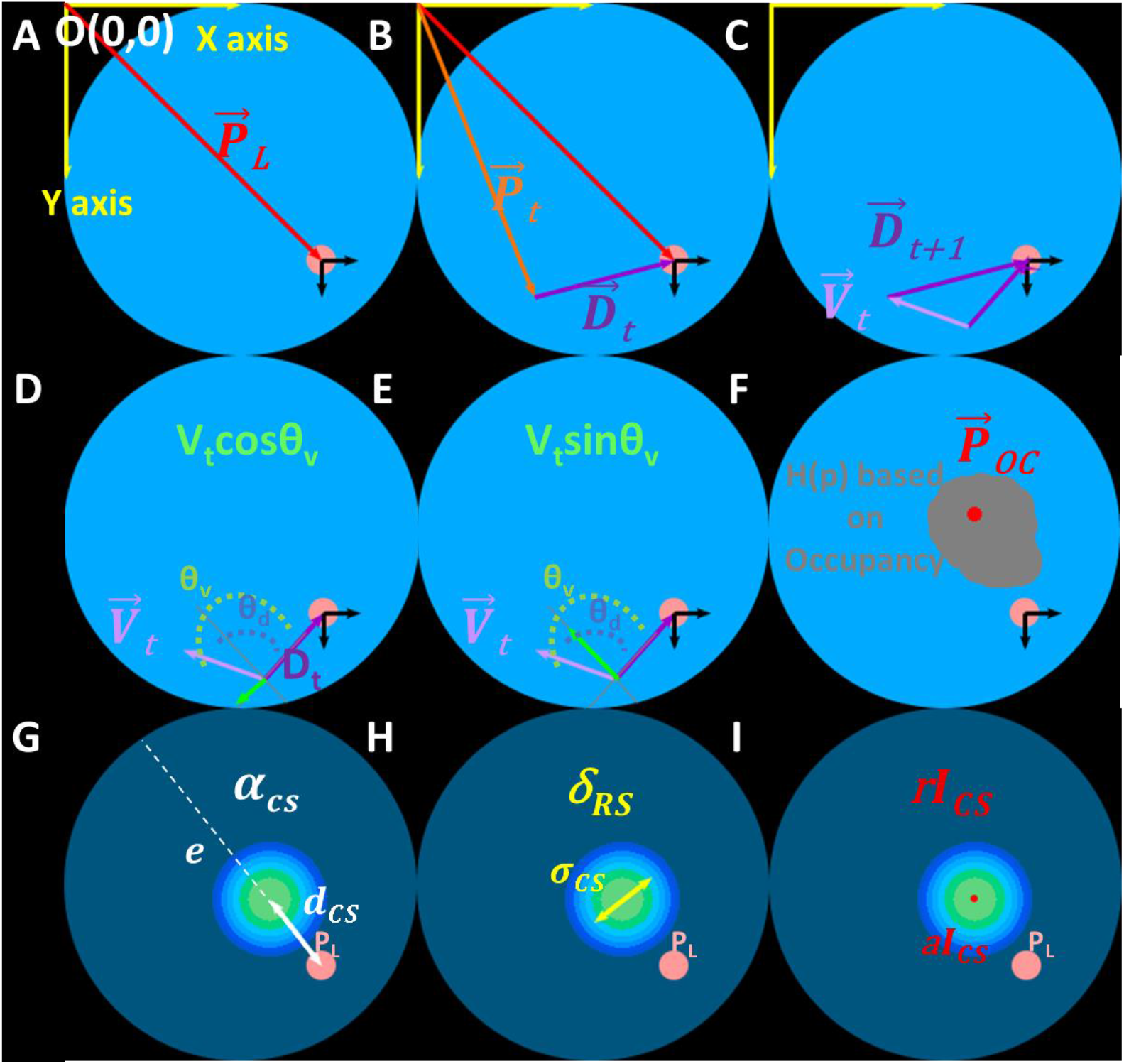
**Schematic representation of the coordinate system used to describe the water maze pool and mouse trajectory as a velocity vector field. Three metrics derived from the velocity vector field that we define, and use, measure the quality of the spatial memory, namely accuracy, uncertainty, and intensity of search (See Theory section for details).** (A) The co-ordinate system and the reference frame that is used to represent the video frame data point acquired from a trial/session of a navigational behaviour. Origin of the coordinate system (O (0,0)) is located at the top left corner of the image. PL (red arrow) is position vector of the platform (pink circle) centre. (B) We define a position vector Pt (orange vector) as the point at which the mouse is located at a given time/video frame (*t*). Thus, the current position of the mouse to the platform location is given by the displacement vector Dt (purple vector). C) Velocity vector Vt (light purple vector) is calculated as the difference in displacement vectors obtained from consecutive video frames. The velocity vector describes the movement of the mouse. It is used as the base measure for developing three metrics for assessing spatial memory and its retention. (D) Resolving velocity vector into its components, a component along a vector pointing towards the platform from current position (*V*_‖*P*_*L*__ = *V*_*t*_ *cos θv*, green vector) measures the mouse’s movement that contributes toward or away from the platform. In this schematic the component is pointing away from the platform. (E) Similarly, the velocity vector component orthogonal to the above component could represent the movement contributing to circular motion of the mice centred about the platform (*V*_⊥PL_ = V_t_ sin θ*v*, green vector). (F) However, to assess the quality of spatial memory, we resolve the velocity vector into its components with respect to the occupancy centre 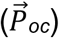 given by the centre of mass (COM) of occupancy. COM calculated on a maximum entropy thresholded occupancy image and represents the most likely region occupied by the mouse during probe trial. We create a vector field from these components to describe the mouse’s intentional movement towards the occupancy centre as well as the extent of circling about that point. Analysing the field properties, namely divergence and curl, allows us to assess the spatial memory. The divergence heat map reveals convergence hotspots, the peak of which represents the putative search centres 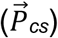. (G) We define accuracy in spatial memory (α_cs_) as a measure that reflects the accuracy with which the mouse remembers the platform location. It is expressed as percentage of 1 minus the fractional error, where the error (dcs) is the displacement between the search centre 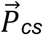 and the platform location PL, and the maximum possible error (e, white dashed line) is the displacement of the farthest boundary/periphery of the pool from the platform. (H) We define our second metric, uncertainty in search, as the spatial spread of the search (σcs), defined as the full width half maxima (FWHM) in x and y axis of the convergence peak. We describe the uncertainty in terms of relative search diameter (δ_RS_), where the search diameter is normalised to platform diameter (dPL). (I) Lastly, intensity of search (Ics) reveals the intensity or intent with which the mouse moves to the search centre. The absolute intensity of search (*aI_cs_*) is represented by the convergence value at the search centre, whereas the relative intensity of search (*rI_cs_*) is the convergence value normalised to the maximum convergence value in the pool space. Since the convergence value looks at the rate of change of the velocity vectors in a small area, the measure is not confounded by differences in swim speed among mice.

**Figure 2:**
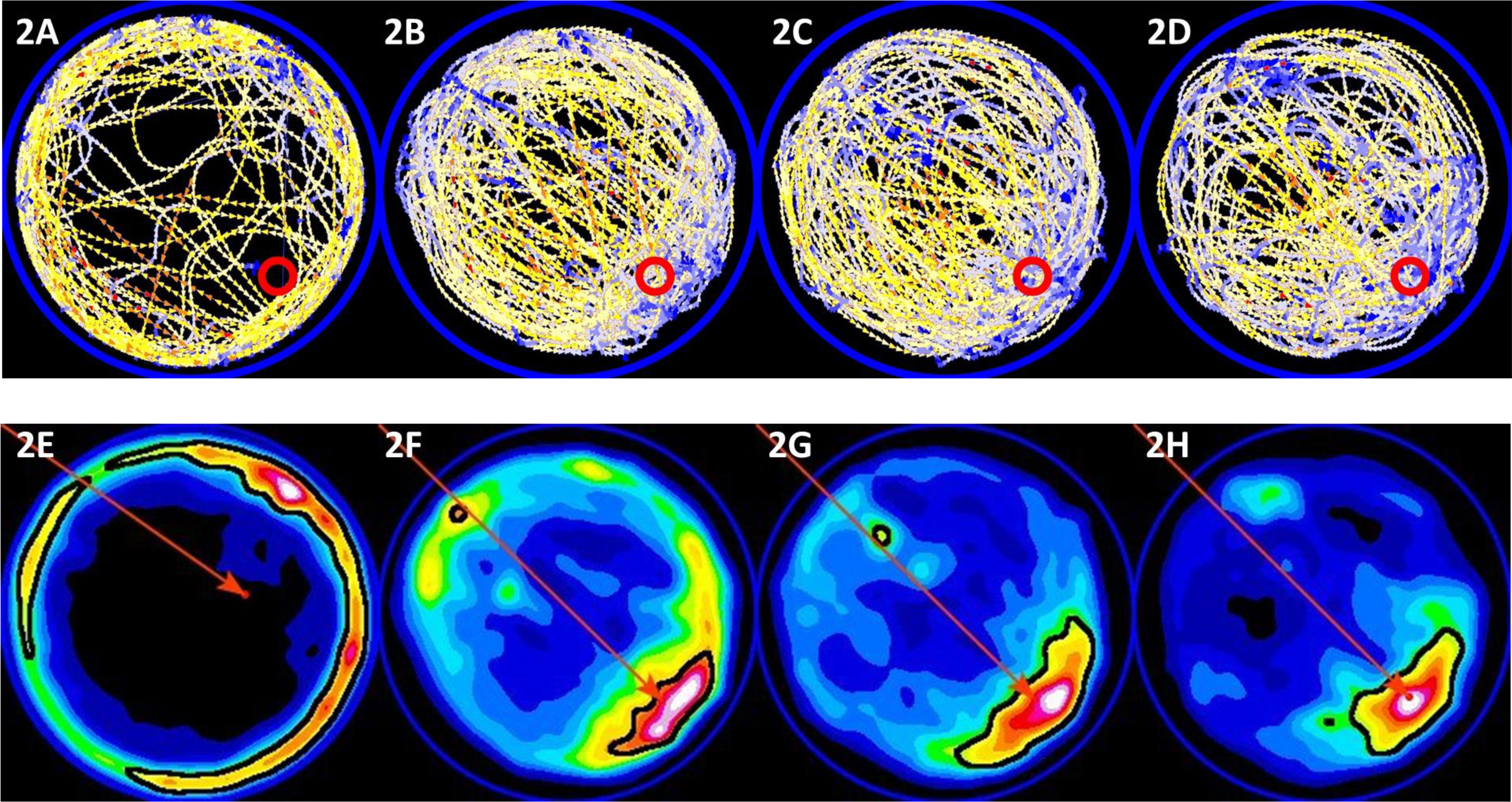
**Velocity vector field, residence time heat map defines the occupancy centre.** The occupancy centre 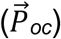 is shown for the population data of *Ptpn11*+/+ wild type mice. (A)-(D) show the velocity vector field on training day 1 (D1), probe day 3 (PD3), probe day 5 (PD5) and probe day 7 (PD7) respectively. LUT scale: 0 – 8 in pixels/frame. (E)-(H) show the occupancy heat map on D1, PD3, PD5 and PD7 respectively. The regions enclosed in black outline represent the most likely regions of occupancy obtained using maximum entropy threshold (see Appendix: Theory for details.) Based on the most likely regions, the occupancy centre is calculated (shown as a red point). The position vector of this point in the image reference frame is shown as red arrows. See Supplementary Figures S1 and S2 for velocity vector field map, residence time heat map showing most likely occupied region, and occupancy centres for all strains/datasets used for analysis in this study. The images are Gaussian smoothed (radius = 6) to approximate the point object to real world dimensions.

**Figure 3:**
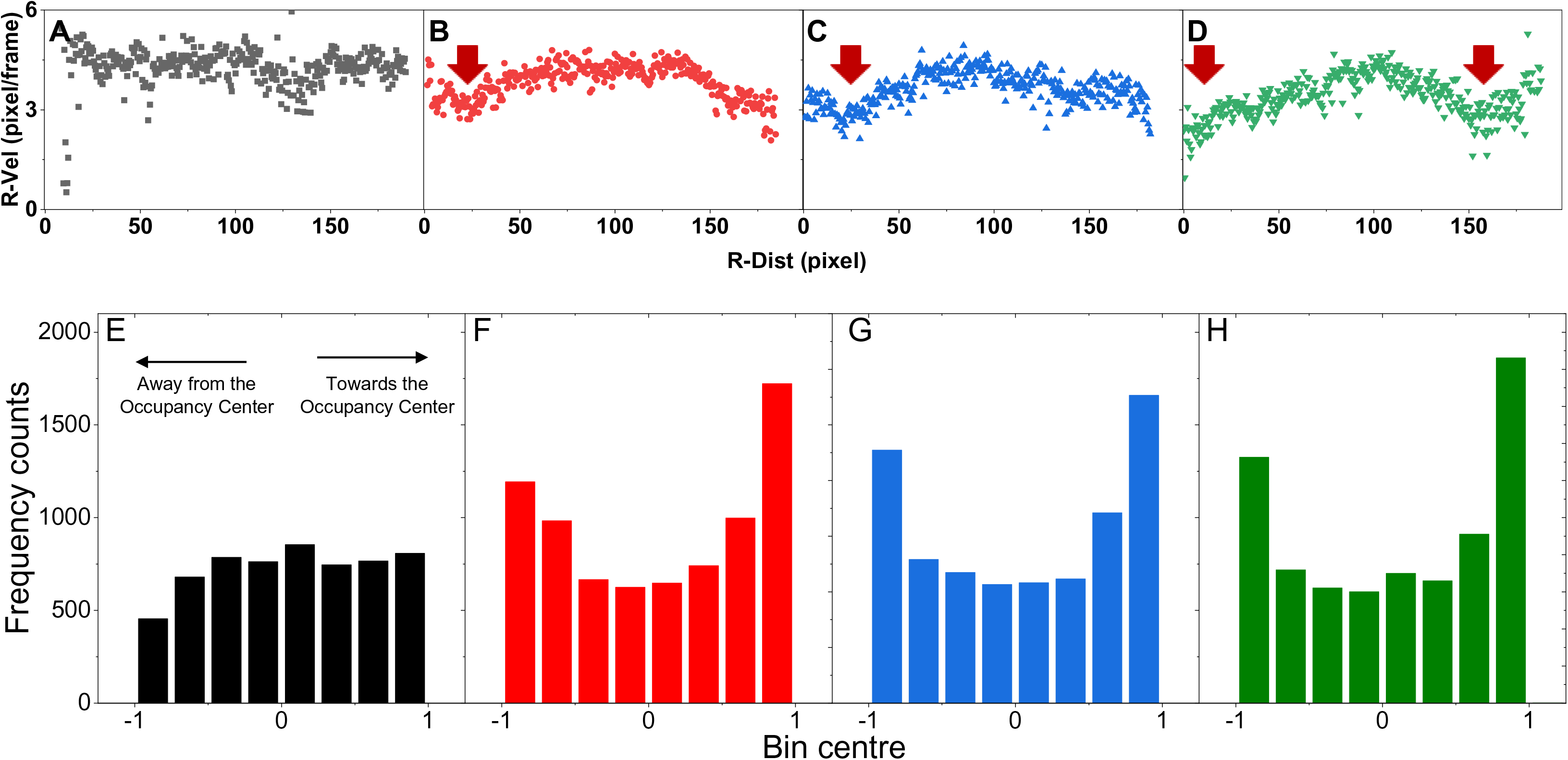
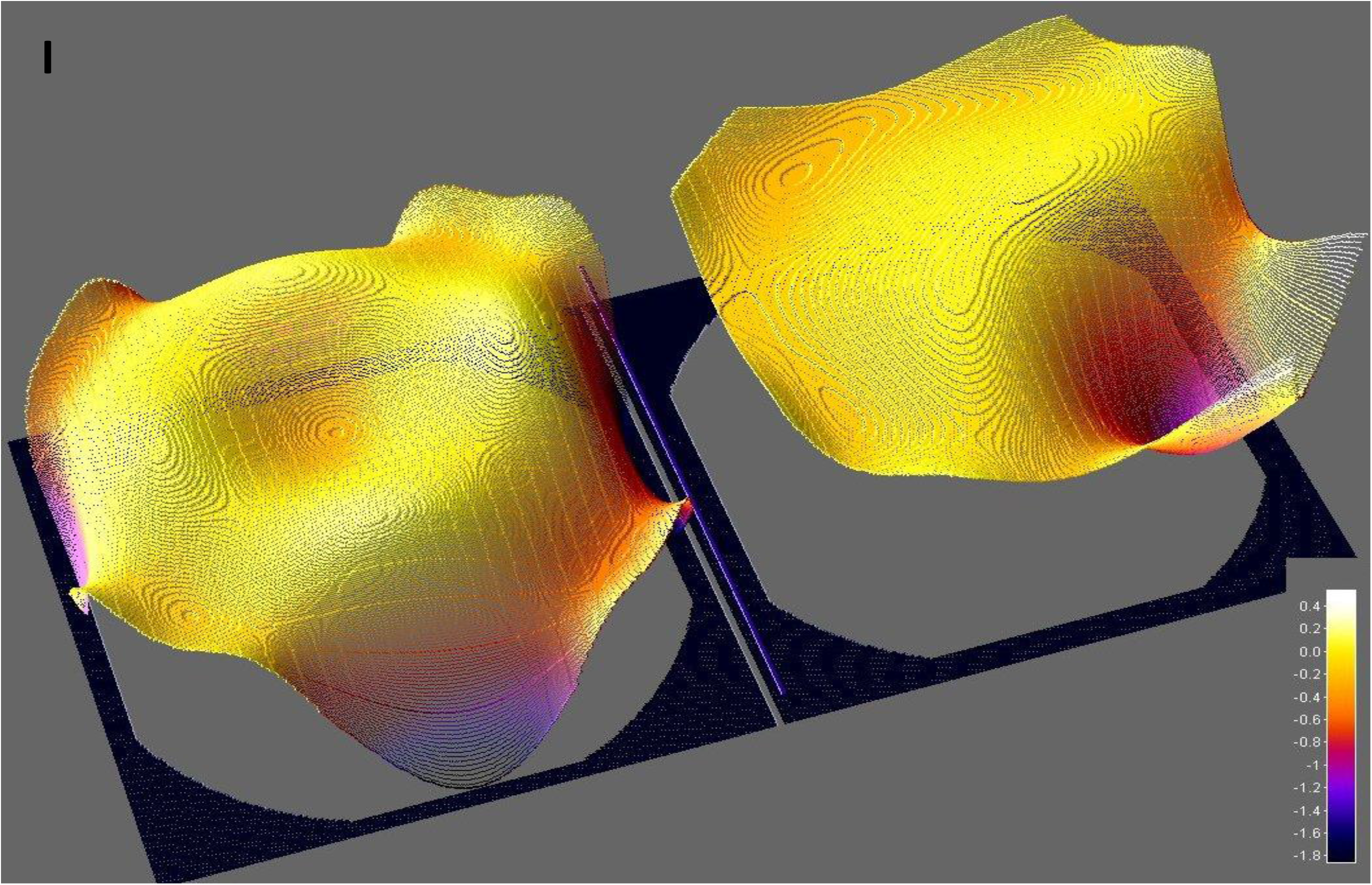
**Divergence maps identify putative search centres or “sinks” as a result of differential distribution of velocity:** (A)-(D) The solid shapes (circles, squares, and triangles) are the magnitude of velocity (R-Vel) measured as a function of radial distance (R-Dist) on training and probe days (D1, PD3, PD5 and PD7 respectively). On D1 the speed of the mice is invariant across space as the scatter plot is almost parallel to the x axis representing the distance from the platform. With training, the speed of the mice is reduced near the platform and boundary (r → 0 or r → 200 pixels). R-Dist is in bins of 0.5 pixel and R-Vel represented as an average of all the times the animal traversed that bin. (E)-(H) Histogram of efficiency of movement toward occupancy centre, given by V||/Vt, in Ptpn11 +/+ mice on D1, PD3, PD5 and PD7 indicated as a function of training. The fraction of the movements oriented along the occupancy centre increases as shown by the counts in bins -1 (aligned completely but away from occupancy centre) and 1 (aligned completely toward the occupancy centre). (I) Surface plot of divergence measure that is developed in theory generated using dataset Ptpn11 +/+ mice on Day 1(left) and on PD7 (right) shows that spatially uniform progressive reduction in speed leading to convergence peaks (negative divergence seen as valleys). Such convergence peaks are analogous to sinks in electro/fluid dynamics. On Day 1 the divergence surface is largely invariant across the pool surface except for a mild depression at the centre and at periphery of the pool. On probe day 7, as the mice acquires the spatial memory for the platform location the surface changes its shape and has a clear valley centred around the platform location.

We reasoned that if a part, and not necessarily the whole of the movement, reflects the mice’s intentions, then we might be able to see a change as a function of training. Accordingly, we observed that the mice show movements that are more aligned to the occupancy center during the probe trials (Fig. 3F-H) compared to that of training (Fig. 3E). Thus, we used the velocity vector component, defined in Eq. (vi), to construct vector fields and obtain the divergence measure (Eq. (viii)). Fig. 3I depicts the surface of the divergence measure estimated using the performance of WT mice during training day 1 (D1) (Left panel) and probe day 7 (PD7) (right panel). This analysis reveals a valley near the platform location as the mice’s performance improves.

### The negative divergence in the vector field of the velocity component along the occupancy center, rather than velocity itself, describes the putative search centers as convergence hotspots for a given swim pattern

To describe swim patterns during probe trials, we constructed a velocity vector field based on the movement of the mice. Together with the velocity vector field, we also constructed a vector field of the velocity component along the occupancy center (Eq.vi). We rationalised that the velocity component along the occupancy center quantifies the effect of an intentional movement towards the occupancy center. We also determined convergence peaks on these vector fields. Given our assertion that for a given swimming pattern, a convergence peak represents the point the mice swim towards, the convergence peak is a proxy for the perceived platform location, as it represents a mouse’s putative search center.

We can generate such a convergence heat map for an individual mouse as well as for a group of mice. For example, we show the velocity vector field maps for an individual *Ptpn11*^+/+^ mouse in Fig. 4(A), as well as the velocity vector field for a group of *Ptpn11*^+/+^ mice (n =15; Fig. 4(C)) for their trajectories on probe day 7. The divergence map calculated on the vector fields shows the presence of convergence peaks near the platform (Fig. 4(F) and 4(H)). In comparison, the divergence map calculated on the velocity component shows a sharp, well-defined convergence peak near the platform site for both the individual and group’s trajectories (Fig. 3I, 4(J) and 4(L)). While the spatial sampling of the pool during the probe trial by an individual mouse is sufficient to detect convergence peaks (thus allowing us to identify putative search centers for each individual mouse), for subsequent analyses, we used the group trajectories.

**Figure 4:**
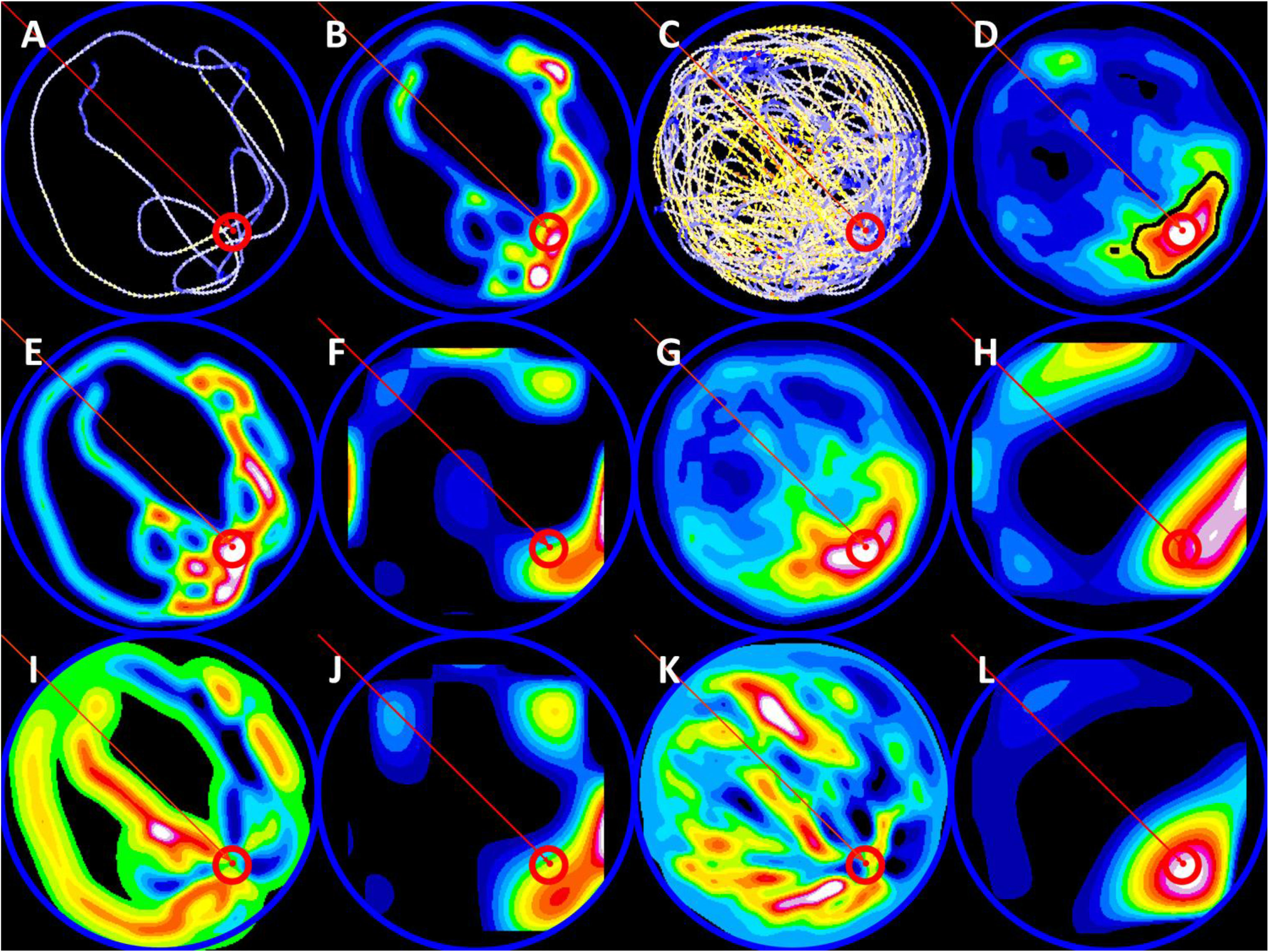
**Component of the velocity along the occupancy centre is better than the velocity at identifying putative search centre.** (A) and (C) show the velocity vector field maps for an individual *Ptpn11*+/+ mouse and a population of *Ptpn11*+/+ mice (n =15) on PD7 respectively LUT scale: 0 – 8 pixel/frame. (B) and (D) show the occupancy map for individual and population of mice. The most likely region is demarcated as a black outline in (D). This region is identified based on maximum entropy segmentation for the population representation. Subsequently, the occupancy centre is estimated and shown as a red arrow in both (B) and (D). (E) and (G) are the velocity magnitude heat maps for the individual mouse and population respectively. Divergence calculated in Cartesian coordinates is used to generate a map and is placed inside the pool image. Bounding rectangle is used to orient and place the divergence map inside the pool. The generated velocity vector fields reveal several convergence hotspots as seen in (F) and (H). In comparison, (I) and (K) represent the heat map representing the magnitude of velocity vector component along the occupancy centre for individual mouse and population, respectively. The divergence calculated as above, using these vector fields, are placed in the pool image and resultant images show relatively fewer but prominent and localised convergence hotspots than (F) and (H). We term these convergence hotspots in velocity component maps as the putative search centres (P_cs_). We use the vector field representing the population of mice during a session in subsequent analysis for identifying the search centre. Blue ROI marks the pool perimeter and red ROI marks the platform.

### Accuracy, uncertainty and intensity of search centers are informative in distinguishing differences between spatial memory of two strains of mice: BALB/cJ and SWR/J

We analysed the WM swim trajectories of the BALB/cJ and SWR/J mouse strains. The divergence heat maps show a distinct convergence hotspot for both, BALB/cJ (n = 10) and SWR/J (n=9/10), groups of mice on all three probe days (i.e., PD3, PD7 and PD10 (Fig.5(A-D)).

**Figure 5:**
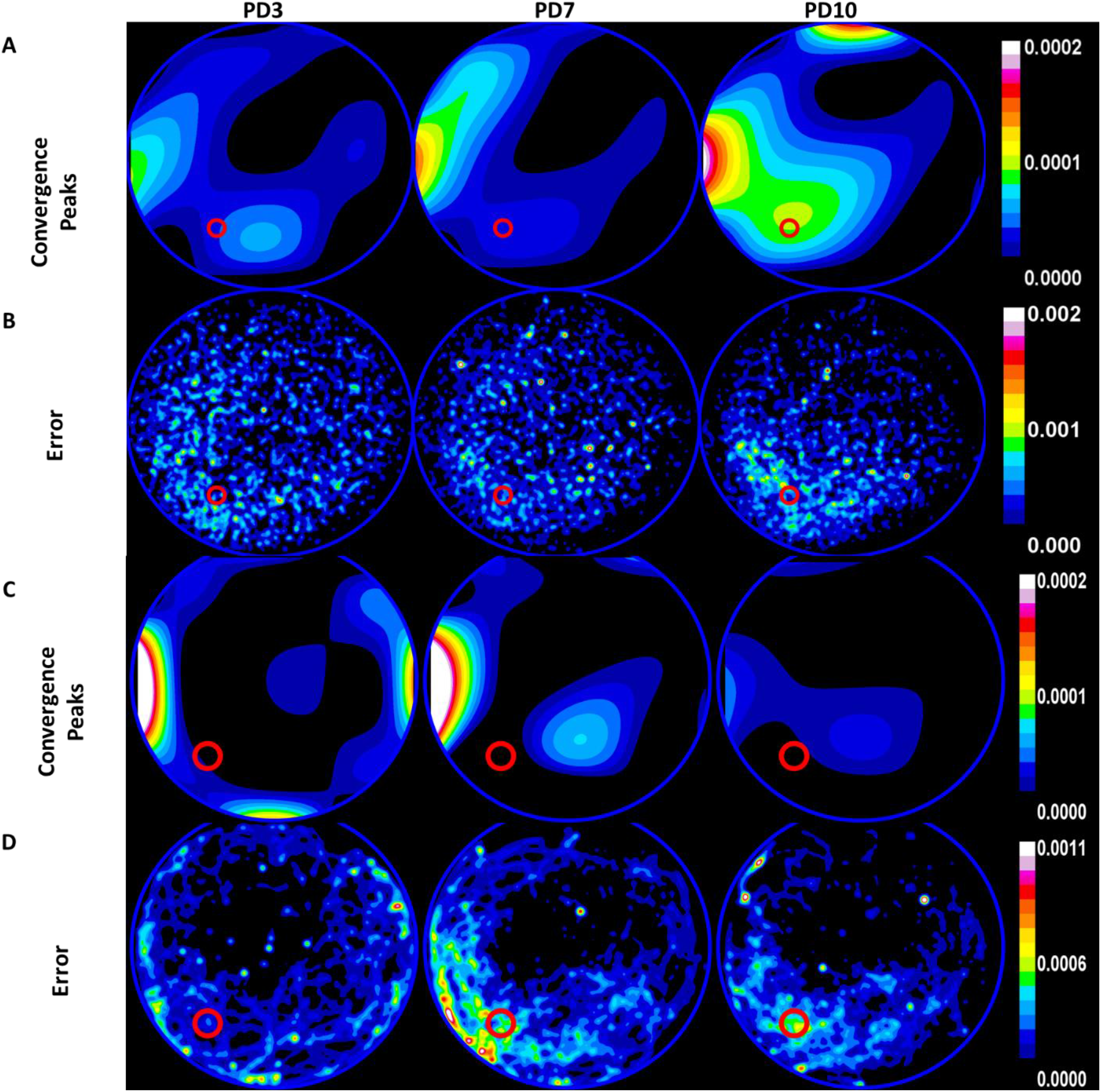
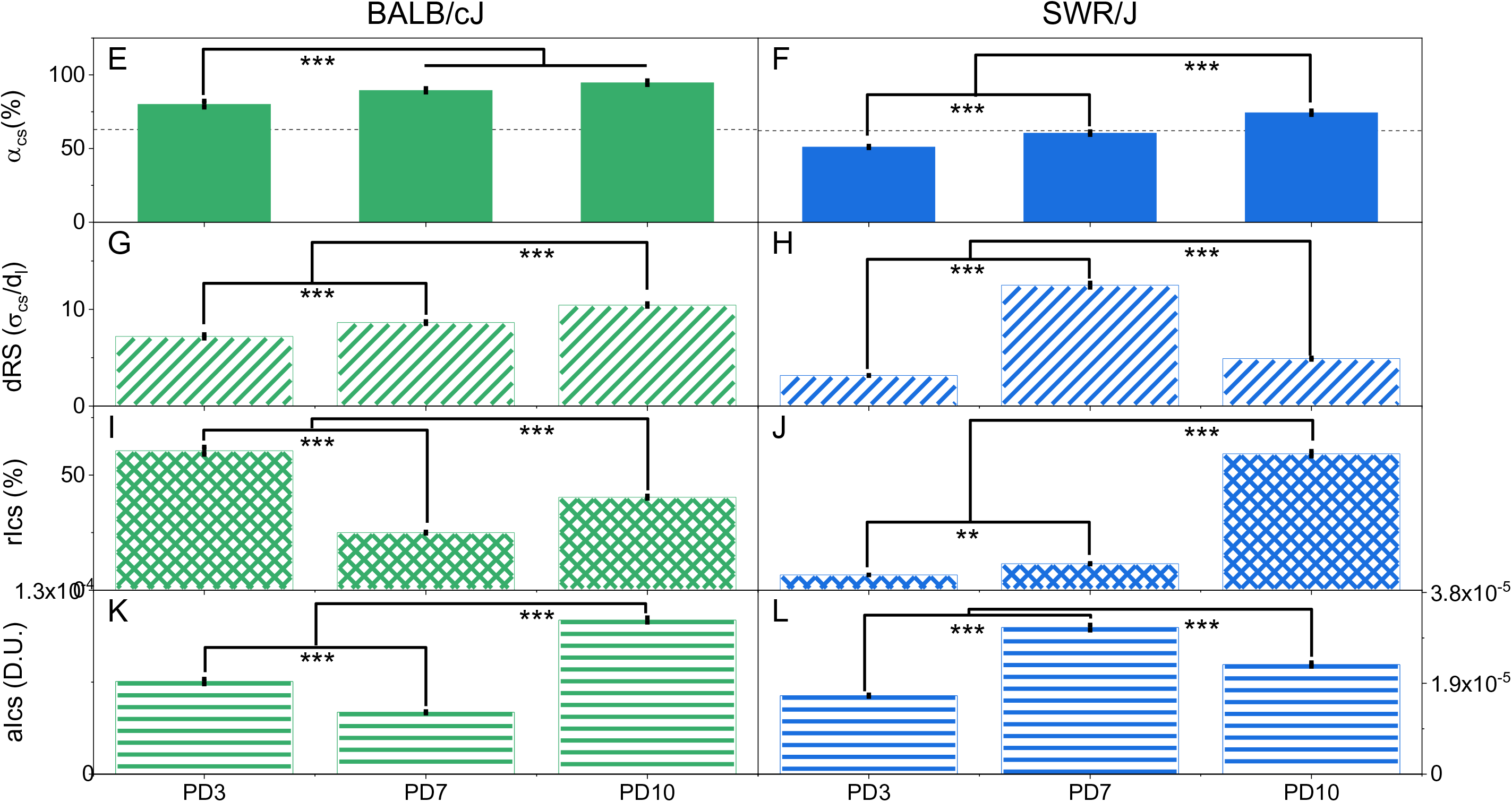
Description of spatial memory in BALB/cJ and SWR/J strain of mice using the three parameters: accuracy (***α***), uncertainty (**δ_RS_**), and intensity of search (***I*_*cs*_**). (A) and (C) are the divergence heat maps showing the convergence hotspots for the population of BALB/cJ (n=10, A) and SWR/J (n = 9-10, C) mice, respectively, on probe day 3 (PD3), probe day 7 (PD7) and probe day 10 (PD10). (B) and (D) are the error heat maps of the calculated divergence within the population of mice of BALB/cJ (B) and SWR/J (D) mice. A 2D Gaussian smoothing of radius = 5 (BALB/cJ) or 10(SWR/J) is used for visual representation. Pool perimeter is shown as a blue circle and the platform location is shown as red circle. (E) and (F) describe the accuracy of search centre for BALB/cJ (E) and SWR/J (F). The accuracy in search increases to maximum asymptotically across probe trials for BALB/cJ mice (Green solid bars, α = 80± 2.2 % (PD3), α = 91 ± 1.3 % (PD7), α = 95 ± 1.5 %(PD10)) as expected. Comparatively, SWR/J mice show poor accuracy for the platform below chance level (defined in theory) on PD3 and at chance on PD7 (One-sample t-test, PD3: p < 0.001, PD5: p > 0.05) whereas it shows an improvement in accuracy (p < 0.001) on PD10 (Blue solid bars, α = 51 ± 0.66 (PD3), α = 61 ± 1.15 (PD7), α = 74 ± 1.41 (PD10)). Dashed line shows the chance accuracy value (BALB/cJ: 63%, SWR/J: 62%). Two-way ANOVA with mice strain and training (measured across probe days) as factors revealed that accuracy is significantly different among the mice strain (F1,52 = 467, p < 0.001), and across training (F2,52 = 82, p < 0.001), however with a significant interaction between the mice strain and their training (F2,52 = 5.66, p< 0.01). Subsequent post hoc analysis indicated that BALB/cJ and SWR/J mice improved its accuracy over training (See table for pair-wise comparisons S. Table 2). Based on the difference in accuracy, we conclude that BALB/cJ are better learners than SWR/J mice. (G) and (H) describes the uncertainty in search centre as relative search diameter (δ_RS_) for BALB/cJ and SWR/J mice respectively. The search strategy around the search centre appears focused in nature on PD3 for BALB/cJ (Green diagonal bar, δ_RS_ = 7 ± 0.20). In case of the SWR/J mice, although the search centre is inaccurate, its search is focused (Blue diagonal bar, δ_RS_ = 3.17 ± 0.04). The relative search diameter increases on subsequent probe trials, indicating that both strains of mice use a more diffused search strategy in later probe trials (BALB/cJ: Green diagonal bars, δ_RS_ = 9 ± 0.13 (PD7), δ_RS_ = 10 ± 0.17 (PD10) and SWR/J: Blue diagonal bars, δ_RS_ = 12.50 ± 0.24 (PD7), δ_RS_ = 5 ± 0.28 (PD10)). Two-way ANOVA for uncertainty with mice strain and training showed that the means of the strains are significantly different (F1,52 = 217, p < 0.001), training had an effect (F2,52 = 586, p < 0.001) however the effect of training is different among the strains as there is an interaction (: F2,52 = 504, p < 0.001) between these factors. Post hoc analysis revealed that the search diameter shows a differential change as function of training (see S. Table 3 for pair-wise comparisons) (I) and (J) represents the relative search intensity (rI_cs_) for BALB/cJ and SWR/J mice. Since the relative intensity of search is obtained from normalising the absolute intensity at search centre with the maximum intensity observed during the session, we can compare the relative intensity across datasets that have been sampled at different frame rates. In BALB/cJ mice, the relative search intensity on the PD3 is the highest (Green hatched bars, rI_cs_ = 61 ± 2%). However, on PD7, the intensity of search reduces (rI_cs_ = 25 ± 0.4%) possibly due to effect of extinction but recovers and increases at PD10 (rI_cs_ = 40 ± 0.6%). In SWR/J mice, the intensity of search improves with probe trials (Blue hatched bars, rI_cs_ = 6.6 ± 0.086% (PD3), rI_cs_ = 11.5 ± 0.22% (PD7), rI_cs_ = 59.3 ± 3.38% (PD10)) though the overall extent of this search intention is poor (<50%). Two-way ANOVA for relative search intensity with strain and training as factors showed that the search intensity significantly differed both as function of strain (F1,52 = 488, p < 0.001) as well as training (F2,52 = 608, p < 0.001) with a strong interaction (F2,52 = 835, p< 0.001). Post hoc analysis (See S. Table 4 for pair-wise comparisons) substantiated our interpretation stated above. (K) represents the absolute search intensity (aI_cs_) for BALB/cJ on PD3, PD7, and P10. In BALB/cJ mice, the intensity of search is given in green horizontal bars, aI_cs_ = 6.5E-5 ± 1.8E-6 (PD3), aI_cs_ = 4.4E-5 ± 6.6E-7 (PD7), aI_cs_ = 1.1E-4 ± 1.7E-6 (PD10)). One-way ANOVA for absolute search intensity showed the means of the absolute search intensity are significantly different (F2,27 = 488, p < 0.001). Post hoc analysis established that the differences between the probe trail days are significant (See S. Table 5 for pair-wise comparisons). (L) represents the absolute search intensity (aI_cs_) for SWR/J mice on PD3, PD7, and P10. In SWR/J mice, the intensity of search improves with probe trials (Blue horizontal bars, aI_cs_ = 1.64E-5 ± 2.13E-7 (PD3), aI_cs_ = 3.07E-5 ± 1.749E-6 (PD7), aI_cs_ = 2.29E-5 ± 4.36E-7 (PD10)). One-way ANOVA for absolute search intensity showed the means of the absolute search intensity are significantly different (F2,25 = 286.7677, p < 0.001). Post hoc analysis established that the progressive increase seen across the probe trial days is significant (See S. Table 5 for pair-wise comparisons).

On calculating the accuracy in search for BALB/cJ mice, we observed as expected that the accuracy increases progressively and reaches an asymptote with additional training (Fig.5E, Green solid bars, α_cs_ = 80 ± 2.2% (PD3), α_cs_ = 91 ± 1.3% (PD7), α_cs_ = 95 ± 1.5% (PD10), S.Table 2)), reflecting accurate knowledge of the platform’s location. Comparatively, SWR/J mice show an accuracy worse than chance levels (as defined in the theory section Eq. (xii)) on a probe trial at day 3 (PD3) and PD7, indicating that the mice are searching away from both the pool center and the platform’s location. Further training of SWR/J mice marginally improves the accuracy on PD10 (Fig.5F, Blue solid bars, α_cs_ = 51 ± 0.66% (PD3), α_cs_ = 61 ± 1.15% (PD7), α_cs_ = 74 ± 1.41% (PD10)). The occupancy-based quadrant and platform measures show that SWR/J mice do not preferentially reside in the target quadrant (P4/Q4) on PD3, but with training, these mice preferentially reside in the training platform zone on PD10 (S.Fig. 4 (H), (J)). However, our analysis of swim trajectories shows that these mice focus their search away from both the platform location and the pool center, indicating that they lack a specific, localised spatial memory of the platform’s location. However, on PD10 these mice do show signs of having acquired a specific spatial memory for the platform’s location.

Additionally, the relative search diameter increases as a function of probe trials for both BALB/cJ and SWR/J mice (Fig.5(G), BALB/cJ: Green diagonal bars, δ_RS_ = 7 ± 0.20(PD3), δ_RS_ = 9 ± 0.13 (PD7), δ_RS_ = 10 ± 0.17 (PD10) and Fig.5(H), SWR/J: Blue diagonal bars, δ_RS_ = 3.17 ± 0.04 (PD3), δ_RS_ = 12.50 ± 0.24 (PD7), δ_RS_ = 5 ± 0.28 (PD10)). With training, their search strategies around the search center appear to shift from focussed and localised searches to more diffused and generalised searches.

For BALB/cJ mice, the search intensity is highest on the first probe trial (PD3; Fig.5(I), Green hatched bars, rI_cs_ = 61 ± 2%). Combined with the accuracy value, these two measures suggest that the mice have learned the platform location and concentrate their searches in the correct location. However, on PD7, there is a reduction in the intensity of the searches (rI_cs_ = 25 ± 0.4%). Since the accuracy of the search center is high (>90 %), the reduction in intensity could possibly be due to a memory of searching for the absent platform in PD3. The results suggest that the memory of the platform location is intact, but that the intention to locate it is reduced because there may be some acquired knowledge during PD3 that it may be absent from the pool. On PD10, the mice show an improvement in search intensity (rI_cs_ = 40 ± 0.6%), potentially reflecting the effect of continued extended training between PD3 and PD10.

In SWR/J mice, the intensity of search improves from PD3 to PD10 (Fig.5(J), Blue hatched bars, rI_cs_ = 6.6 ± 0.086% (PD3), rI_cs_ = 11.5 ± 0.22% (PD7), rI_cs_ = 59.3 ± 3.38% (PD10). The extent of this search intention is poor (<50%) for PD3 and PD7, reflecting poor spatial learning.

Thus, using convergence peaks as search centers, we can describe and compare the quality of spatial learning and memory across different probe trials for different strains of mice. From our comparisons, we find that the BALB/cJ mice perform better in the MWM than the SWR/J strain of mice.

### *Ptpn11*^D61G/+^ mice learn the platform location with additional training as indicated by accuracy, uncertainty, and intensity of search

Next, we analysed the water maze performance of *Ptpn11* ^D61G/+^ mice, a model of Noonan syndrome (Araki et al., 2004). These mice have a gain-of-function point mutation in non-receptor protein tyrosine phosphotase *Ptpn11* gene found in Noonan patients (Araki et al., 2009, 2004), and they show learning and memory impairments, including spatial learning and memory deficits in the MWM (Lee et al., 2014). Previous studies (Lee et al., 2014) showed that *Ptpn11*^D61G/+^ mice do not learn and remember the platform location after three training sessions with 4 trials each, as indicated by quadrant occupancy and proximity measure on the first probe trial (PD3). In contrast, with the same training WT littermates (*Ptpn11*^+/+^ mice) show clear evidence of having learned the platform’s location. Additional training failed to improve the performance of *Ptpn11*^D61G/+^ mice, as reported by quadrant occupancy and proximity measures on PD7. Our methods of analyses indicate that although *Ptpn11*^D61G/+^ mice did not show evidence that they had learned the platform location on PD3, additional training does improve their performance in the MWM: The divergence on velocity vector field along the occupancy center reveals a search center near the platform location on PD5 and PD7 (Fig.6(C)). Describing the convergence peak or search center in terms of accuracy of search, uncertainty about search, and intensity of search, reveals more clearly the quality of the spatial memory performance of *Ptpn11*^D61G/+^ mice during probe trials. The accuracy of the search center to the platform location in *Ptpn11*^+/+^ mice is not statistically different across probe days (Fig.6(E), Black solid bars, α = 59 ± 2.25% (D1), α = 93 ± 2.41% (PD3), α = 93 ± 2.60% (PD5), α = 96 ± 1.34% (PD7)). Interestingly, the accuracy of the search center in *Ptpn11*^D61G/+^ mice on PD5 is comparable to that of *Ptpn11*^+/+^ mice on PD3 (Fig.6(F), Red solid bars, α = 60 ± 2.10% (D1), α = 62 ±1.91% (PD3), α = 87 ± 1.57% (PD5), α = 92 ± 2.12% (PD7) ; Two-way ANOVA for accuracy with genotype as between-subjects factor and probe day within-subjects factor, genotype × probe day interaction: F_3,92_ = 21, p< 0.001; Bonferroni: *Ptpn11*^+/+^ PD3 – *Ptpn11*^D61G/+^ PD3: p < 0.001, *Ptpn11*^+/+^ PD3 – *Ptpn11* D61G/+ PD5: p > 0.5).

**Figure 6:**
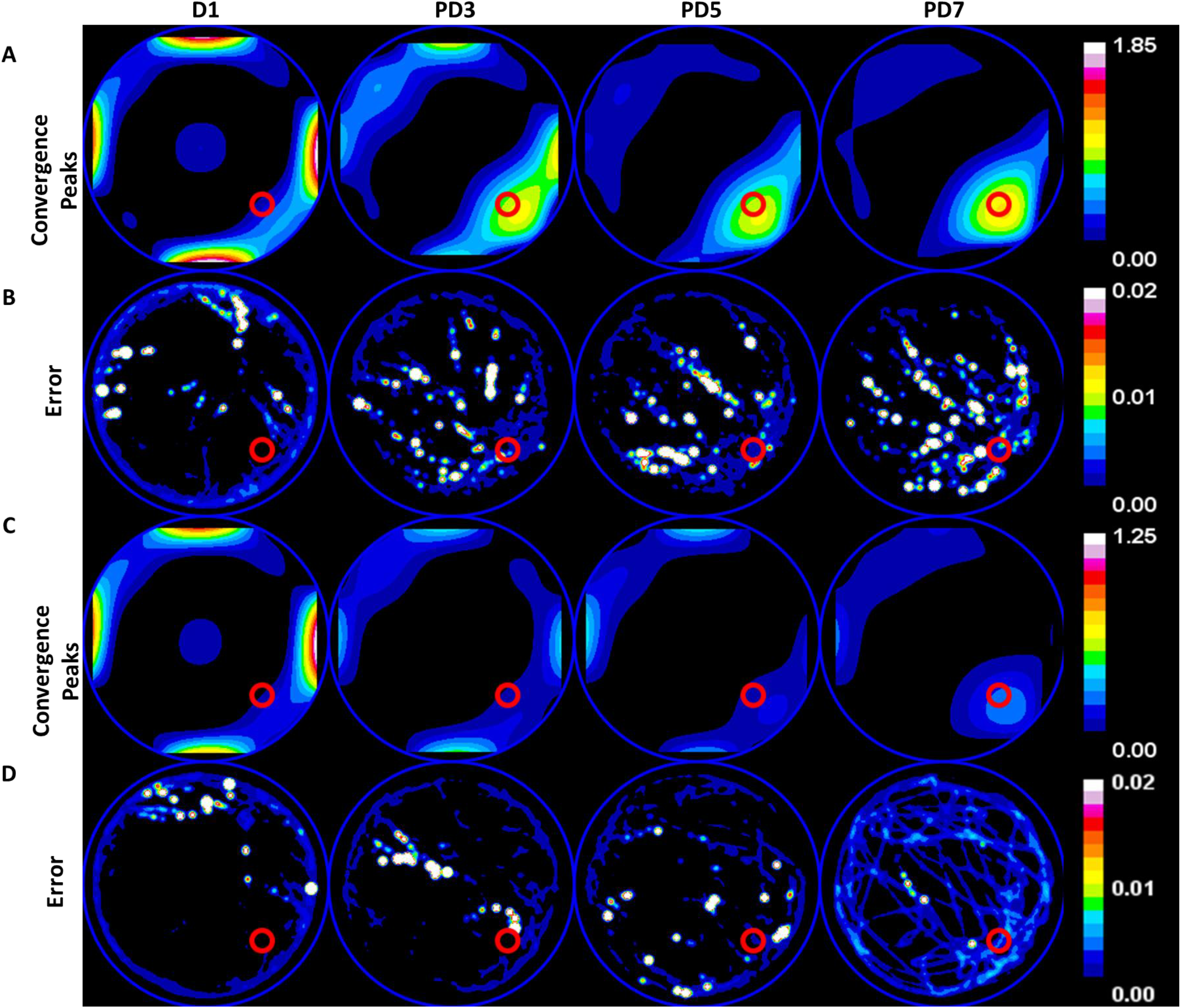
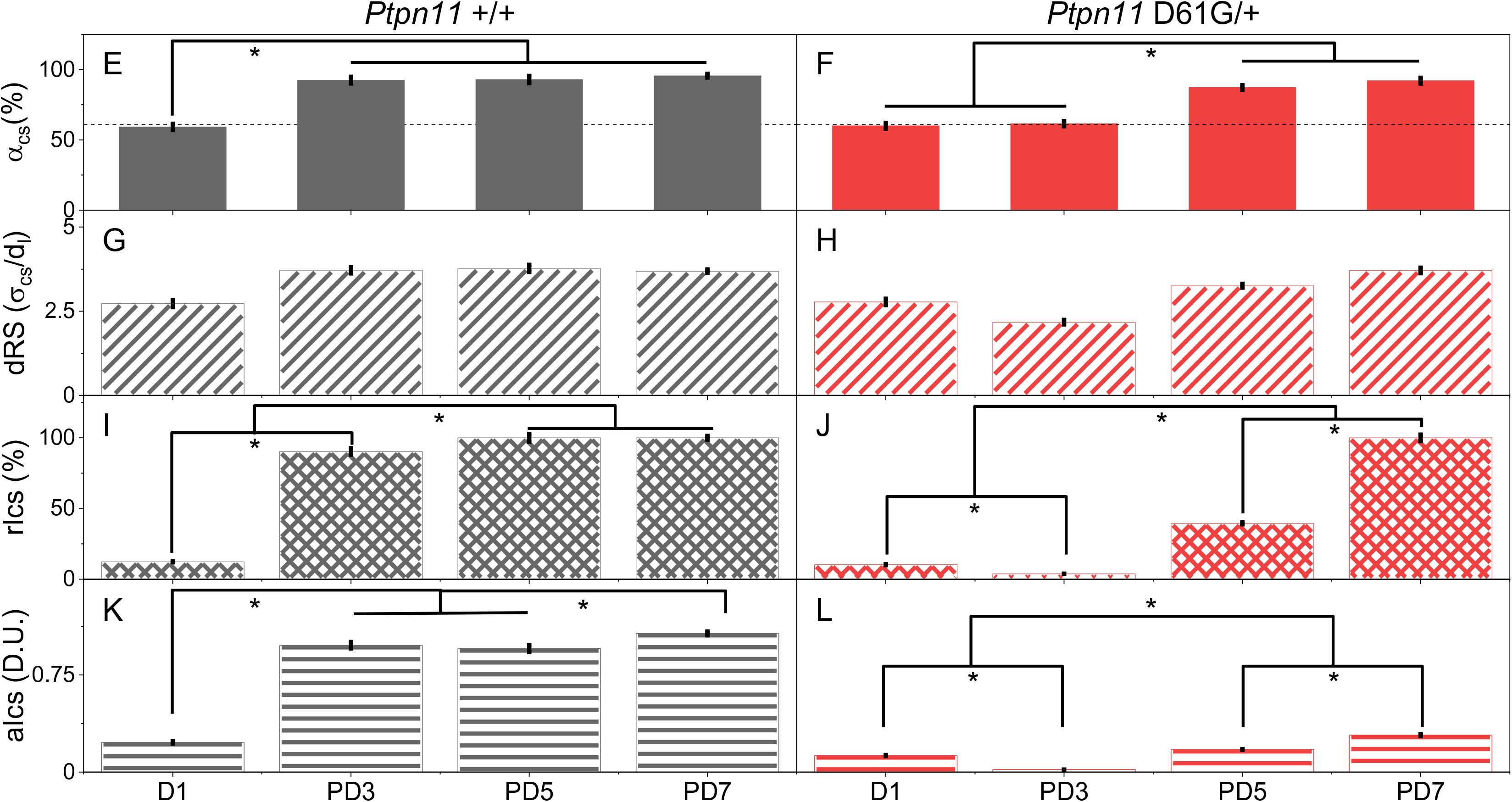
**Description of MWM spatial memory in *Ptpn11* D61G/+ and wild type litters mates (*Ptpn11* +/+) using the three parameters accuracy (*α*), uncertainty (*δ*_RS_), and intensity of search (*I*_*cs*_) show that with additional training D61G/+ mice recall the platform location with accuracy and relative search intensity comparable to wild type littermates (PD3).** (A) and (C) are the divergence heat maps showing the convergence hotspots for the population of *Ptpn11* +/+ wild type (WT, n = 15) and *Ptpn11* D61G/+ transgenic (D61G, n = 10) mice, respectively, on training day 1 (D1), probe day 3 (PD3), probe day 5 (PD5) and probe day 7 (PD7). D1 heat map serves as a representation of the surface when the mouse has no knowledge of the platform location. After 3 training sessions of 4 trials each, WT mice show a convergence peak close to the platform location in the first probe trial on PD3, whereas the D61G/+ mice do not show any peak near the platform. With additional training, D61G show a convergence peak adjacent to the platform location on PD5 and PD7 indicating the presence of spatial memory. (B) and (D) is a visual representation of the error estimates of calculated divergence within a population of mice for *Ptpn11*+/+ and *Ptpn11* D61G/+ mice respectively. A gaussian blur of radius 2 was applied for visual representation. Pool perimeter is shown as a blue circle and the platform location is shown as red circle. (E) and (F) quantify the accuracy of search centre. In both wildtype and mutant mice, the accuracy of search centre starts at a chance level (61%, dashed line) on D1 (WT: α = 59 ± 2.25 (D1, black solid bars); one-sample t-test, p > 0.05 and D61G/+: α = 60 ± 2.10 (D1, red solid bar); one-sample t-test p > 0.05). In WT mice the accuracy reaches a maximum on PD3 itself (α = 93 ± 2.41 (PD3), α = 93 ± 2.60 (PD5), α = 96 ± 1.34 (PD7)). We see that D61G/+ mice perform at chance level when tested on PD3 (α = 62 ±1.91 (PD3, red solid bar); one-sample t-test, p > 0.05). However, with additional training, the accuracy of search centre improves on PD5 and PD7 (Red solid bars α = 87 ± 1.57 (PD5), α = 92 ± 2.12 (PD7); Two-way ANOVA for accuracy with genotype and training as factors showed that the means across the genotype (F1,92 = 39, p < 0.001), as well as across training day (F3,92 = 97, p<0.001) are different with a significant interaction(F3,92 = 21, p< 0.001). Post hoc analysis (See S. Table 6 for pair-wise comparison) are consistent with our interpretation provided above. Interestingly, we see that the accuracy of TG mice on PD5 is comparable to that of WT mice on PD3 (Bonferroni: WT PD3 – D61G/+ PD3: p < 0.001, WT PD3 – D61G/+ PD5: p > 0.5) indicating that D61G/+ mice are able to learn the spatial memory task despite their cognitive deficits however with a slow rate of acquisition. (G) and (H) quantifies the uncertainty in search centre in terms of relative search diameter (δ_RS_) for WT and D61G/+ mice. In both WT (Black diagonal bars, δ_RS_ = 2.7 ± 0.103 (D1), δ_RS_ = 3.7 ± 0.097 (PD3), δ_RS_ = 3.7 ± 0.106 (PD5), δ_RS_ = 3.7 ± 0.052 (PD7)) and D61G/+ (Red diagonal bars, δ_RS_ = 2.7 ± 0.097 (D1), δ_RS_ = 2.18 ± 0.067 (PD3), δ_RS_ = 3.3 ± 0.059 (PD5), δ_RS_ = 3.7 ± 0.085 (PD7)) mice, the relative search diameter was found to be comparable across the three probe trials. Two-way ANOVA for uncertainty in search with genotype and training as factors showed that the difference in means across both, the genotype (F1,92 = 1.78, p > 0.05) and across training day (F3,92 = 1.47, p > 0.05), is not statistically significant. The interaction between the genotype and training day was also not statistically significant (F3,92 = 0.996, p > 0.05, See S. Table 7). This indicates that the search area or focus does not change across probe trials. (I) and (J) quantifies the relative intensity of search (rI_cs_) for WT and D61G/+ mice. In case of WT mice, the relative intensity of search at the search centre (P_cs_) is high on PD3 itself and reaches a maximum on PD5 and PD7 (Black hatched bars, rI_cs_ = 12 ± 1 (D1), rI_cs_ = 90 ± 9 (PD3), rI_cs_ = 100 ± 11 (PD5), rI_cs_ = 100 ± 5 (PD7)). In D61G/+ mice, the intensity of search at P_cs_ steadily improves from a low value on PD3 to a maximum on PD7 (Red hatched bars, rI_cs_ = 10 ± 1 (D1), rI_cs_ = 3.8 ± 0.26(PD3), rI_cs_ = 40 ± 2 (PD5), rI_cs_ = 100 ± 9 (PD7)). Two-way ANOVA for relative intensity with genotype and training as factors show that the means are significantly different both as a function of genotype (F_1,92_ = 1.3E7, p < 0.001) and training day (F_3,92_ = 31.3E7, p < 0.001 The interaction between the two factors was also statistically significant (F_3,92_ = 4.5E6, p< 0.001). Post hoc analysis shows that the WT mice learned to search intently on PD3 (D1 < PD3) even though PD5 and PD7 search intensities are higher (PD3 < PD5, PD7) while the mutant mice progressively increased its search intensity from only after PD3 (D1 ∼ PD3), that is intensity of search at PD3 < PD5 < PD7 as seen from pairwise comparisons in S. Table 8. The evidence further supports the notion that D61G/+ mutant mice learn the platform location and contribute its effort at the search centre after intense training. (K) and (L) quantifies the absolute intensity of search (aI_cs_) for WT and D61G/+ mice. The absolute intensity of search at P_cs_ is significantly different on probe days compared to training day in WT mice (Black horizontal bars, aI_cs_ = 0.229 ± 0.0087 (D1), aI_cs_ = 0.98 ± 0.02548 (PD3), aI_cs_ = 0.955 ± 0.02674 (PD5), aI_cs_ = 1.072 ± 0.01501 (PD7)). In D61G/+ mice, the absolute intensity of search at P_cs_ steadily improves with training (Red horizontal bars, aI_cs_ = 0.128 ± 0.00448 (D1), aI_cs_ = 0.019 ± 5.89E-4 (PD3), aI_cs_ = 0.175 ± 0.00315 (PD5), aI_cs_ = 0.285 ± 0.00656 (PD7). Both D61G as well as their littermates are trained and probed on the same experimental setup (with experimenter being blind to the genotype) as a result the absolute intensities can be compared across genotypes. Two-way ANOVA for comparing absolute intensity across genotypes and training as two factors indicate that absolute search intensities are significantly different between WT and D61G mice(F_1,92_ = 2677, p < 0.001) and as a function of training they increase (F_3,92_ = 285, p< 0.001) their intention of search though at different rates (genotype × probe day interaction: F_3,92_ = 224, p< 0.001). See S. Table 9 for pair-wise comparison. This we interpret as resulting from possible motor as well as cognitive deficits.

Additionally, the search intensity relative to the pool (rI_cs_) increases as a function of training. *Ptpn11*^D61G/+^ mice increased their focused searches and their intention at search center on PD7, and these are similar to their WT littermates in this probe trial (Fig.6(I-J)). However, there is a significant difference in the absolute intensity of search (aI_cs_) between *Ptpn11*^+/+^ and *Ptpn11*^D61G/+^ mice (Fig.6(K-L)). Absolute intensity at a location in a divergence image measures the extent by which vector decreases over a unit distance. Given that in our case the video is uniformly sampled, this measure directly reflects a decrease observed over time, and hence the velocity. On the other hand, relative intensity measures the fraction of slow down with respect to the steepest decline observed in the pool. Thus, the differences in search intensities (al_cs_) could reflect either cognitive or motor deficits, while rl_cs_ differences reflect differences in spatial learning and memory. The difference we observe in search intensities (al_cs_) in WT and mutants could be due to the differences in their swimming ability. In contrast, rl_cs_, accuracy and uncertainty are sensitive to deficits in spatial learning and memory and are not directly affected by differences in motor performance.

To further test the increased sensitivity of our methods and reduce the effect of motor impairment characteristic of Noonan syndrome (Lee et al., 2014; Romano et al., 2010), we a ran another set of analyses with a smaller number of animals in which the Noonan Syndrome mutation was restricted to the dorsal hippocampus (HPC), a region related to spatial memory processing (Andersen et al., 2009). Previous results showed that the adeno-associated virus induced expression of the PTPN11^D61G^ allele (AAV-PTPN11^D61G^) only in the HPC leads to spatial memory deficits similar to transgenic animals, including in the MWM (Lee et al, 2014). The SFig.6A and B show the convergence map generated for this set of mice at D1, PD5, PD7 and PD11. Subsequent measures of accuracy (SFig. 6C and D), uncertainty (SFig. 6E and F) and relative intensity of search (SFig. 6G and H) show clear evidence for learning induced behavioral changes, while the residence time measure (SFig. 6I and J) of this subset does not show any discernible change suggesting that statistical power of residence time measure is substantially lower and requires a greater number of mice to be used for detecting these subtle changes.

### The spatial learning and memory deficit of *Ptpn11* ^N308D/+^ mice in the MWM is rescued by SL327

Next, we analysed the MWM performance of another mouse model of Noonan syndrome (*Ptpn11* ^N308D/+^). *Ptpn11*^N308D/+^ mice have an heterozygous knock-in gain-of-function mutation found in some Noonan patients (Araki et al., 2009, 2004) and their spatial learning and memory deficits are known to be milder than those in *Ptpn11*^D61G/+^ mice (Lee et al., 2014). While *Ptpn11*^N308D/+^ mice show similar performance to WT littermates as measured by latency to reach the platform during training, during a probe trial after three days of training (PD3), these mice spend significantly less time in the target quadrant than their WT littermates. *Ptpn11*^N308D/+^ mice also search farther away from the platform, as shown by the proximity measure. However, with two additional days of training, these mice show comparable spatial learning and memory performance in PD5, as measured using quadrant (SFig. 4 (AA-AD)) and proximity measures (SFig. 5 (I)).

We analysed the swim trajectories of *Ptpn11*^N308D/+^ and *their WT* littermates on PD5. Additionally, we investigated whether SL327, a MEK inhibitor, previously shown to reverse the memory deficits of *Ptpn11*^D61G/+^ mice (Lee et al., 2014), could also improve the accuracy and uncertainty in search of the platform location in *Ptpn11*^N308D/+^ mice. First, we found that there are subtle but significant differences in spatial learning and memory performance in terms of accuracy, uncertainty, and intensity of search between *Ptpn11*^N308D/+^ and their WT littermates. *Ptpn11*^N308D/+^ showed an accuracy of 80 ± 2% with an uncertainty of 5.5 ± 0.16 relative search diameter, whereas the WT mice showed a superior accuracy of 94 ± 2 % with an uncertainty of 4.01 ± 0.092 in relative search diameters (refer to S. Tables 10 and 11 for statistical comparison). After treatment with SL327, *Ptpn11*^N308D/+^ improved their accuracy to 95 ± 2% with a relative search diameter of 4.33 ± 0.074 (S. Tables 10 and 11) Incidentally, treating WT mice with SL327 has the opposite effect in their accuracy and search area, i.e., α_cs_ = 88 ± 4% and δ_RS_ = ± 0.235 ((Fig.7(C-E)), Two-way ANOVA for accuracy with genotype as between-subjects factor and treatment within-subjects factor, genotype × treatment: F_1,38_ = 15, p< 0.001; Bonferroni: Ptpn11 N308D/+ Veh – Ptpn11 N308D/+ SL p < 0.01, Ptpn11 N308D/+ Veh – Ptpn11 +/+ Veh: p < 0.01). These results are consistent with our interpretation that that there is an impairment in spatial memory in *Ptpn11*^N308D/+^ mice and that the MEK inhibitor SL327 is capable of rescuing this deficit even though it causes a mild decline in performance in WT littermates, a compelling demonstration of the specificity of this drug treatment.

**Figure 7:**
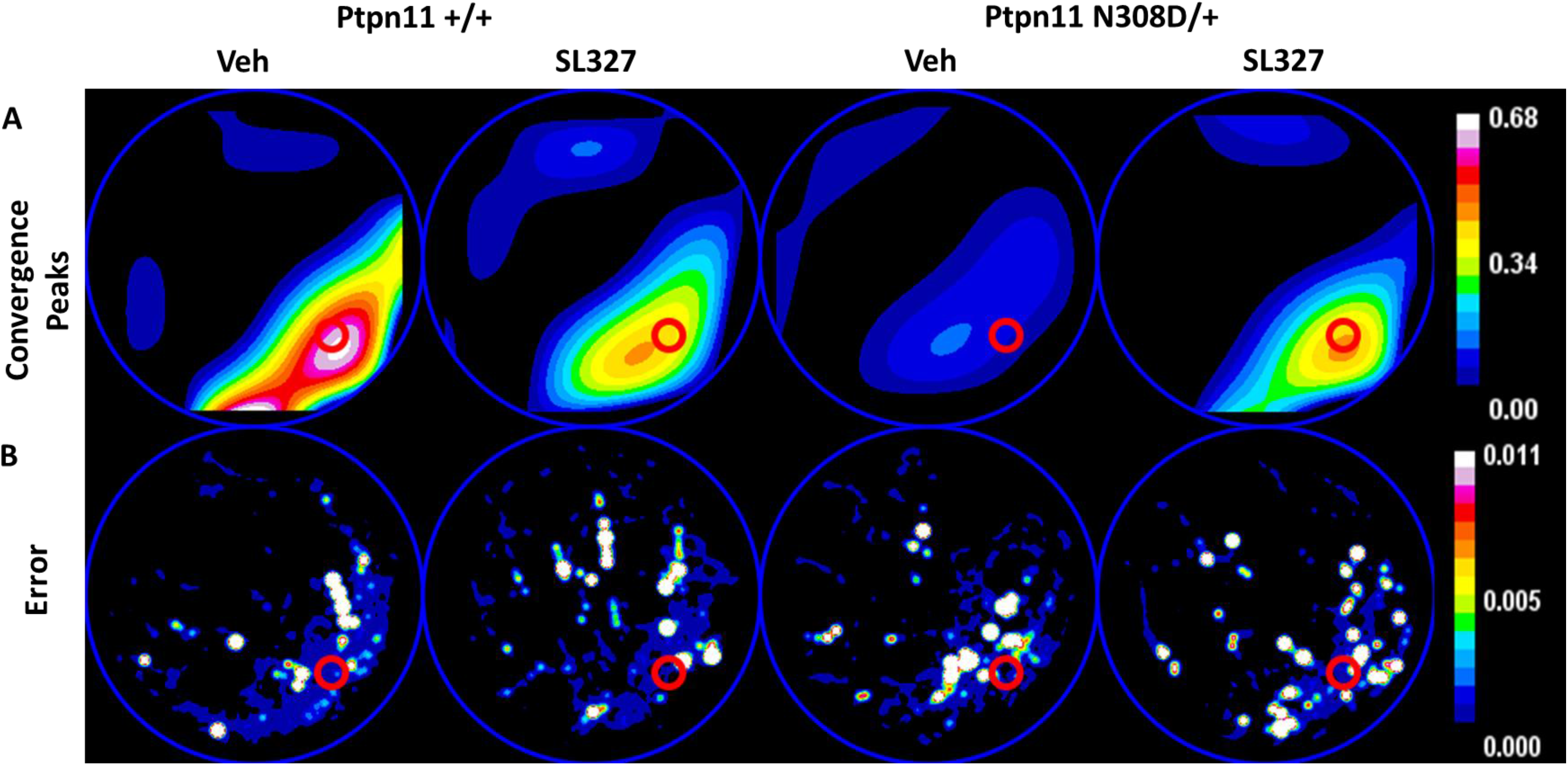
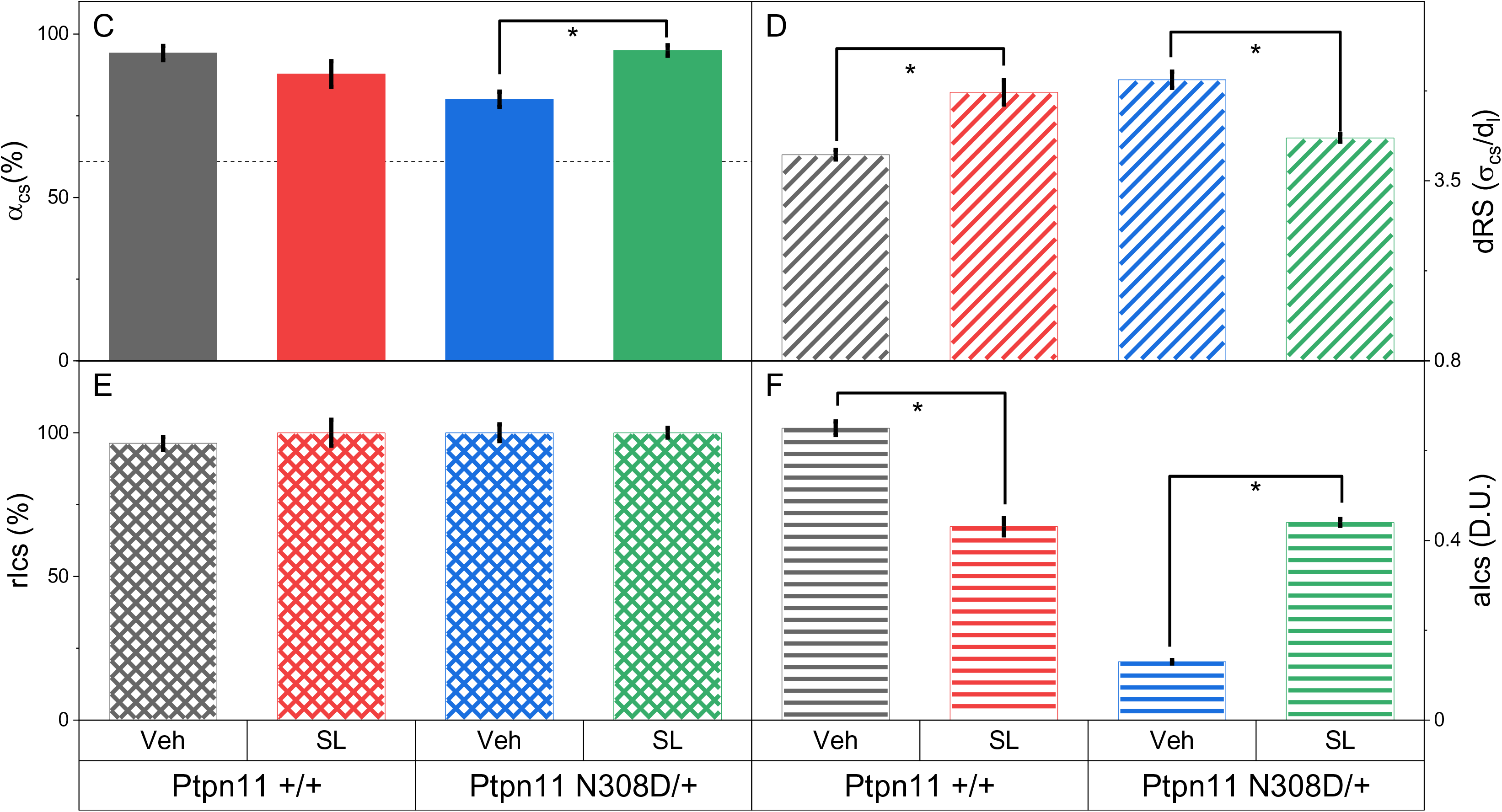
**Accuracy and uncertainty are able to detect the rescuing of spatial memory deficits seen in *Ptpn11* N308D/+ mice by the administration of SL327.** (A) Divergence heat maps representing the convergence peaks for *Ptpn11*+/+ WT mice (n=11) and *Ptpn11* N308D/+ mutant mice (n=10) for saline (veh) and drug (SL327) treated conditions. (B) Heat map of the error estimated on divergence values indicates the variation within the population of mice. A gaussian blur (radius = 6) was applied for visualising the error in the sampled pixels. (C) Accuracy of search centre reveals that *Ptpn11* +/+ and *Ptpn11* N30D/+ show subtle difference in spatial memory (Solid bars: *Ptpn11* +/+ Veh α_cs_ = 94 ± 2 % (black), *Ptpn11* N308D/+ Veh α_cs_ = 80 ± 2% (red)). Treatment with MEK inhibitor SL327 rescues the deficit and brings the performance of N308D/+ mutant to the level of WT (*Ptpn11* +/+ SL327 α_cs_ = 88 ± 4 (blue), *Ptpn11* N308D/+ SL327 α_cs_ = 95 ± 2% (green); Bonferroni: *Ptpn11* N308D/+ Veh – *Ptpn11* N308D/+ SL p < 0.01, *Ptpn11* N308D/+ Veh – *Ptpn11* +/+ Veh: p < 0.01.). Two-way ANOVA for accuracy with genotype and treatment as two factors established that although there is no difference in the means of genotype (F_1,38_ = 1.6, p > 0.05) and the treatment (F_1,38_ = 2.4, p > 0.05) however there was significant interaction (F_1,38_ = 15, p< 0.001) indicating that the means of these factors are changing in opposite directions as revealed by the post hoc analysis with the rescue due to SL327 is significantly higher than the decrement in accuracy seen in the wild type following the administration of SL327 (See S. Table 10 for pair-wise comparisons). Dashed line shows the chance accuracy value (61%). (D) Uncertainty in spatial memory is statistically different between N308D/+ mutant and WT littermates (Diagonal bars: *Ptpn11* +/+ Veh δ_RS_ = 4.01 ±0.092 (black), *Ptpn11* N308D/+ Veh δ_RS_ = 5.47 ± 0.159 (red); Bonferroni: *Ptpn11* N308D/+ Veh – *Ptpn11* +/+ Veh: p < 0.001). Administration of SL327 makes the search focussed for the mutant *(Ptpn11* N308D/+ SL327 δ_RS_ = 4.33 ± 0.074 (green)), however it also increases the search diameter in the wild type llittermates. (*Ptpn11* +/+ SL327 δ_RS_ = 5.22 ± 0.235 (blue),). N308D/+ mutant mice improve their performance (Bonferroni: *Ptpn11* N308D/+ Veh – *Ptpn11* N308D/+ SL p < 0.001), whereas the search area of their wildtype littermates becomes more diffused. Two-way ANOVA for uncertainty performed with genotype and treatment showed significant interaction (genotype × treatment interaction: F_1,38_ = 56, p< 0.001), without any main effect (genotype: F_1,38_ = 3.3, p > 0.05, treatment: F_1,38_ = 0.068, p < 0.05) (,. See S. Table 11 for pair-wise comparison. (E) The relative search intensities at the search centre are similar in both strains of mice (Hatched bars: *Ptpn11* +/+ Veh rI_cs_ = 96 ± 2 (black), *Ptpn11* N308D/+ Veh rI_cs_ = 100 ± 3 (red)). Treatment of SL327 did not alter the relative intensity contributed to the search centre in case of both strains of mice (*Ptpn11* +/+ SL327 rI_cs_ = 100 ± 5 (blue), *Ptpn11* N308D/+ SL327 rI_cs_ = 100 ± 2 (green)). Two-way ANOVA for relative search intensity with genotype and treatment as two factors found no significant difference among the means nor any significant interaction between the factors. (genotype: F_1,38_ = 0.36, p > 0.05, treatment: F_1,38_ = 0.36, p > 0.05, genotype × treatment interaction: F_1,38_ = 0.36, p > 0.05. See S. Table 12). (F) The absolute intensities measured from these two groups of mice share the same units as they are carried out on the same setup at the same frame rate. The intensity of search differs in case of *Ptpn11* N308D and its WT littermates (Horizontal bars: *Ptpn11* +/+ Veh aI_cs_ = 0.65 ± 0.015 (black), *Ptpn11* N308D/+ Veh aI_cs_ = 0.13 ± 0.004 (red)).. Treatment of SL327 rescues the deficit seen in N308D mice such that the treated mice search more intently than that of the vehicle treated mutant mice. Although SL327 affects the performance of the WT type mice but the direction of change is opposite to that of the N308D mutant. (*Ptpn11* +/+ SL327 aI_cs_ = 0.43 ± 0.019 (blue), *Ptpn11* N308D/+ SL327 aI_cs_ = 0.44 ± 0.007 (green); Bonferroni: *Ptpn11* N308D/+ SL – *Ptpn11* +/+ SL p > 0.5.) Two-way ANOVA done on the absolute search intensity with genotype and treatment as two factors showed that means of both the factors (genotype: F_1,38_ = 1749, p < 0.001, day: F_1,38_ = 55, p< 0.001), as well as interaction is significantly different (F_1,38_ = 1217, p< 0.001) Post hoc analysis (S. Table 13) confirms our earlier interpretation that SL327 rescues the spatial memory deficit seen in N308D mice although the treated group search intention is still significantly lesser than the wild type vehicle. We also note that administration of SL327 significantly lowers the search intention in the wild type animals.

### Tracking performance in a goal reversal task in the MWM

Finally, we analysed a publicly available dataset with two strains of mice (10-week old female mice C57Bl/6J and DBA/2J) tested in a goal reversal task (Overall et al., 2020). These analyses highlight the power of the current vector field property-based measures in detecting search centers. In the goal reversal task, the mice are thought to acquire a memory for the old as well as for the new platform locations, and due to a recency effect, they bias their search towards the new platform location.

Previous results showed that DBA/2J mice perform poorly in hippocampal-dependent tasks, including in the MWM (Paylor et al., 1994; Upchurch & Wehner, 1988). Indeed, our analyses revealed that DBA/2J mice do not show the presence of a specific search center on PD3 (Fig.8(C)). In comparison, analyses of PD3 revealed that C57Bl/6J mice learn the platform location with an accuracy of 86 ± 3 %, relative search diameter of 4.4 ± 0.15 and relative search intensity of 100 ± 3% (Fig.8(A), (E), (G),(I)).

**Figure 8:**
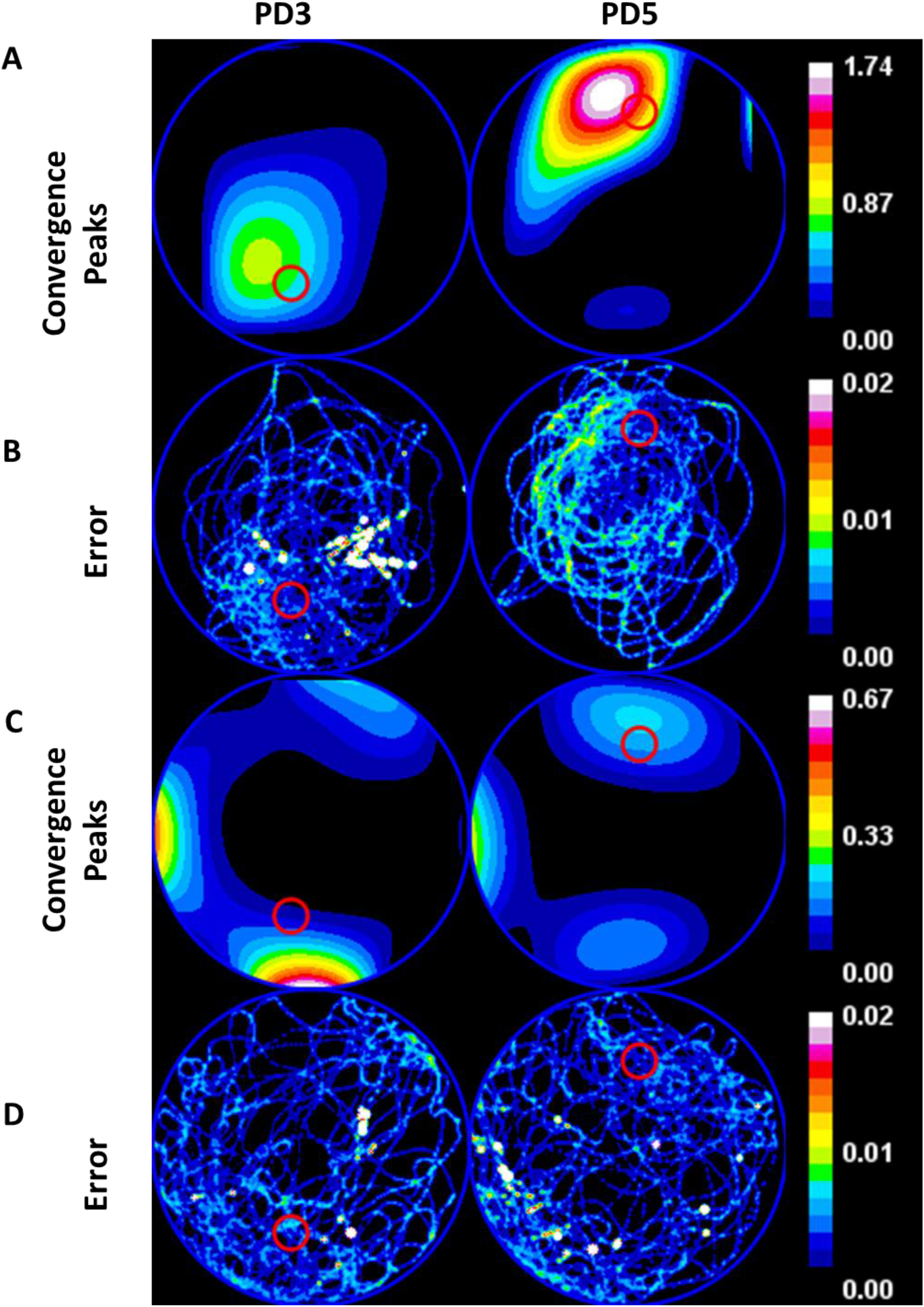
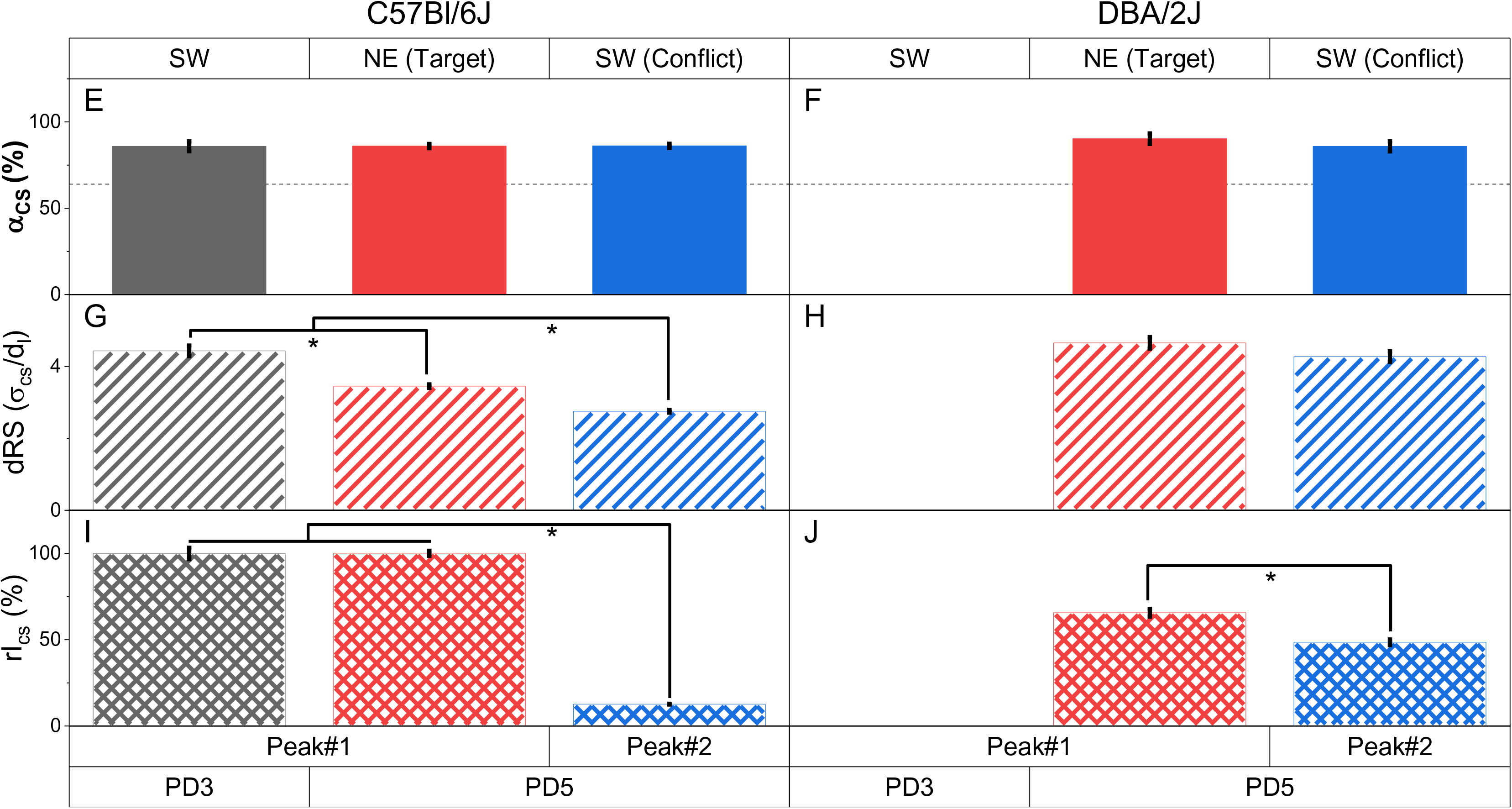
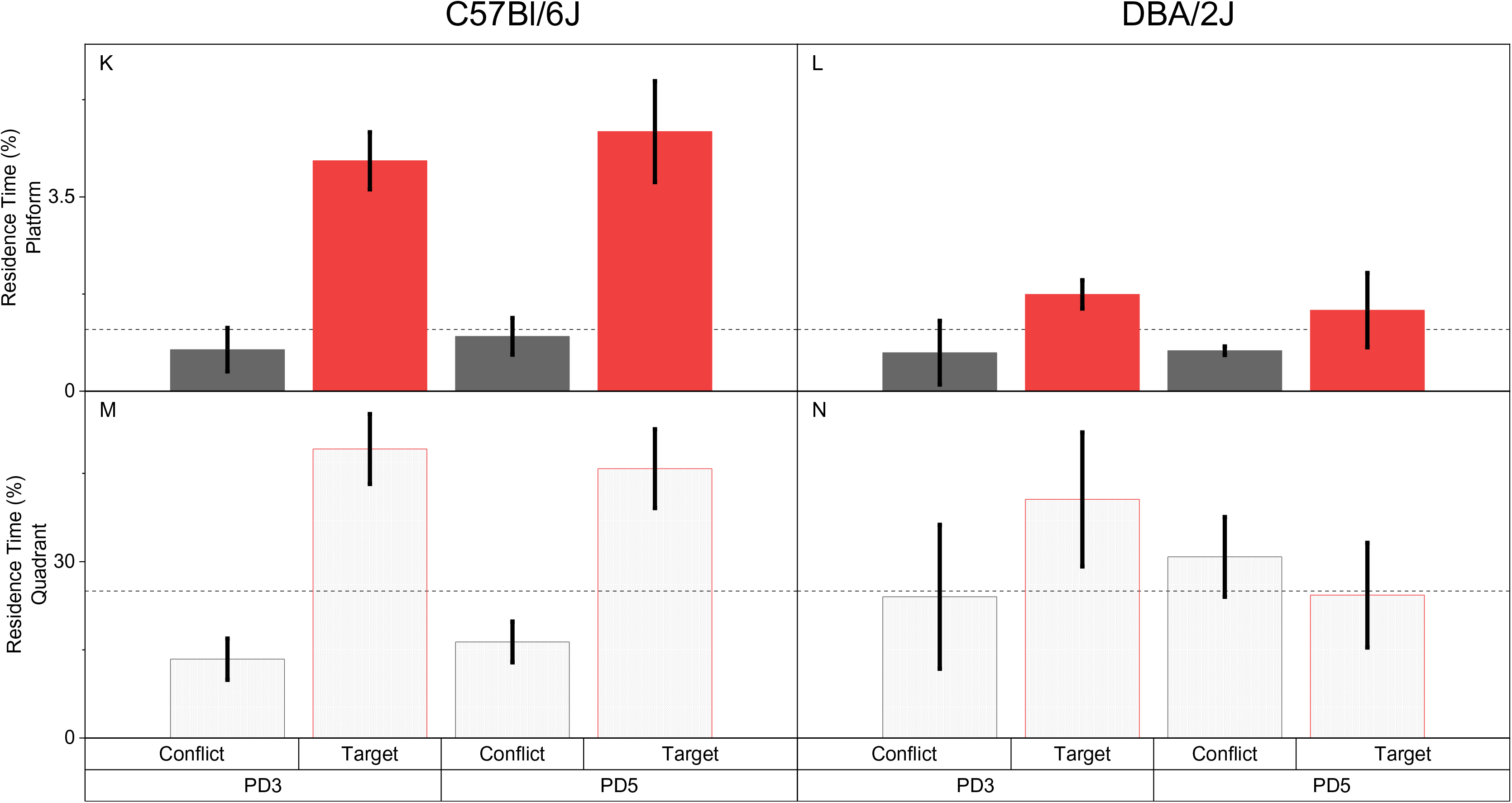
Convergence hotspots detects the memory for old as well as new platform location in goal reversal task. C57Bl/6J and DBA/2J strains of mice were trained with the platform in the south-west (SW) quadrant on probe day 3 (PD3), whereas the platform was shifted to the north-east (NE) quadrant on probe day 5 (PD5). Using accuracy (α), uncertainty (δ_RS_), and intensity of search (*I*_*cs*_), we show that C57Bl/6J and DBA/2J mice have memory for both platform locations, whereas the residence time measure shows that on PD5, C57Bl/6J mice do not search in old platform location above chance and that DBA/2J mice did not learn either of the platform location. (A) and (C) shows the divergence heat map for the population of C57Bl/6J and DBA/2J mice (n = 5). C57Bl/6J mice show a distinct peak on PD3 and two peaks on PD5. In contrast, DBA/2J mice do not show a convergence peak on PD3 but show two peaks on PD5. We consider both peaks on PD5 for further analysis and label the peak in NE quadrant/new platform location as Peak#1 and peak in SW quadrant/ old platform location as Peak#2. (B) and (D) are the error in divergence contributed by variation within the population. A gaussian blur (radius = 4.75) was applied for visualising the sampled pixels. (E) describes the accuracy of search centre for C57Bl/6J on PD3 and PD5. C57Bl/6J show a high degree of accuracy (Solid black bar, α = 86 ± 2.8) on PD3. On PD5, the accuracy of Peak#1 measured with respect to the target platform (NE quadrant) (Solid red bar: α_cs_ = 86 ± 1%), as well as the accuracy of Peak#2 measured with respect to the conflict platform location (SW quadrant) (Solid blue bar: α_cs_ = 86 ± 1%) is close to maximum. These results indicate that Bl6 mice have a highly accurate and precise spatial memory for both the platform locations, i.e. target location for PD5 and the conflict location on PD3. One-way ANOVA F_2,12_ = 0.0034, p > 0.05. (F) describes the accuracy of search centre for DBA/2J mice on PD3 and PD5. The absence of a convergence peak on PD3 indicates that DBA/2J do not possess spatial memory for the PD3 platform location. However, on PD5, DBA/2J mice have two search centres as shown in the convergence heat map. The accuracy of Peak#1 measured with respect to the target platform (NE quadrant) (Solid red bar: α = 90 ± 3%), as well as the accuracy of the Peak#2 measured with respect to the conflict platform location (SW quadrant) (Solid blue bar: α = 86 ± 3%) is maximum. One-way ANOVA F_1,8_ = 1.09, p > 0.05. Thus, illustrating the ability of our method to detect and describe the aspect of spatial memory corresponding to two distinct platform locations. Dashed line shows the chance accuracy value (64%) in plots (E)-(F). Two-way ANOVA done for comparing the accuracy on PD5 with strain and peaks (platform locations) as factors showed that the means are not significantly different and there is no significant interaction (Strain F_1,16_ = 0.78, p > 0.05, peak: F_1,16_ = 0.897, p > 0.05, strain × peak interaction: F_1,16_ = 0.973, p > 0.05. (See S. Table 14(b)). (G) describes the uncertainty in search for C57Bl/6J. On PD3, the relative search diameter is (Diagonal black bar) δ_RS_ = 4.43 ± 0.15. The relative search diameters of Peak#1 new PL (Diagonal red bar, δ_RS_ = 3.45 ± 0.048) and Peak#2 old PL (Diagonal blue bar, δ_RS_ = 2.75 ± 0.039) are not statistically different (One-way ANOVA F_2,12_ = 85, p < 0.001), thus, the uncertainty in search is similar for both the platform locations (SW and NE). (H) describes the uncertainty in search for DBA/2J mice. The relative search diameters of Peak#1 new PL (Diagonal red bar, δ_RS_ = 4.66 ± 0.16) and Peak#2 old PL (Diagonal blue bar, δ_RS_ = 4.27 ± 0.15) are not statistically different (One-way ANOVA F_1,8_ = 3.18, p > 0.05), thus, the uncertainty in search is similar for both the platform locations (SW and NE). Two-way ANOVA for mean uncertainty on PD5 with strain and peak as two factors showed that the uncertainty is different across the strain (F_1,16_ = 148, p <0.001), and for location of the peaks (F_1,16_ = 23, p < 0.001) with no significant interaction(F_1,16_ = 1.993, p > 0.05). From Post hoc analysis we see the search diameter is lesser for the older location then the new one (See S. Table 15(b)) reduces the uncertainty in search area. (I) Relative intensity of search shows C57Bl/6J mice maximally search at platform location on PD3 (Hatched black bar, 100 ± 3%) while on PD5 they search at both the new platform location (Hatched red bar, 100 ± 1%) as well as the old location (Hatched blue bar, 12.7 ± 0.2%). However, the intensity of search is higher for new location (One-way ANOVA F_2,12_ = 592, p < 0.001). (J) Relative intensity of search of DBA/2J mice on PD5. The relative intensity of search for the platform at the target location (NE) (Hatched red bar, 66 ± 2 %) is significantly more than at the conflict location (SW) (Hatched blue bar, 49 ± 2%) (One-way ANOVA F_1,8_ = 38, p < 0.001). Thus, the mice search more intently in the target location, compared to the conflict location as a result of goal reversal training. We note that intensity of search for the old platform location in DBA/2J is considerably more compared to the C57Bl/6J. Two-way ANOVA for relative search intensity with strain and peak as two factors show that difference in means across the strains is not significant : (F_1,16_ = 0.17, p > 0.05) but mean intensity of search for the peaks are significantly different (F_1,16_ = 1127, p < 0.001)with a significant interaction(F_1,16_ = 510, p < 0.001) between the factors. Post hoc analysis (See S. Table 16(b)) shows that goal reversal is complete for C57Bl/6J as indicated by lower search intensity for the old platform location while DBA/2J mice shows significantly more freezing at the old platform location. We estimate the efficiency of goal reversal training as (Inew – Iold)/(Inew + Iold). Thus, a value of 1 would indicate complete goal reversal while -1 would indicate no reversal with zero indicating that there is a conflict. Here we find this efficiency for C57Bl/6J to be ∼0.78 and DBA/2J to ∼0.15. (K) and (M) The zone and quadrant residence time measure on PD5 for C57BL/6J mice. Both measures show that the mice spend significantly more time in the target location (See SFig. 4 and the corresponding tables), whereas it resides in the conflict location at chance level. Thus, the residence time-based measures are unable to detect the presence of spatial memory for the older platform location. (L) and (N) The zone (solid bar) and quadrant (dotted bar) residence time measure PD5 for DBA/2J mice. DBA/2J mice occupy the target zone or quadrant at chance level. Thus, the measures are unable to detect the presence of spatial memory for both platform training locations. Dashed line represents the chance value (1.11% and 25% respectively) in plots (K)-(N).

As predicted, after being trained on two platform locations, analyses of the mice’s performance with our methods reveals the presence of two convergence peaks or search centers on probe day 5 (PD5). The two search centers correspond to the presumed memories for the target location (i.e., the platform location in north-east quadrant on training days 3 and 4), as well as memory for the conflict location (i.e., the platform in the south-west quadrant on training on days 1 and 2 (Fig.8(A-D)). Quantifying the search centers in terms of accuracy, uncertainty and intensity of search reveals differences in the nature of the two memories.

Interestingly, we did not detect a difference in accuracy of the search centers to their respective platform positions (i.e., search center near the target location (Peak#1 new) and search center near the conflict location (Peak#2 old)). We conclude that since the mice recall the old as well as the new platform locations with similar accuracy, the strength of the memory for both locations is comparatively high (Fig.8(E-F)). Thus, learning the second location does not seem to significantly weaken the memory for the first location.

In C57Bl/6J mice, the relative intensity of the search (r_Ics_) at the search center corresponding to the new platform location (Peak#1 new), is significantly higher than the relative intensity of search corresponding to the old platform location (Peak#2 old; Fig.8(I)), indicating that a possible recency effect for the new platform location biases the mice to search more intensely in the target location. However, when the mice fail to find the platform at the target location, they also investigate the other (i.e., conflict) location. Nevertheless, the mice spend considerably less time searching in the conflict location. Similarly, the relative intensity of search (rI_cs_) at the search center corresponding to the new platform location (Peak#1 new), is slightly higher than the relative intensity of search corresponding to the old platform location in DBA/2J mice (Peak#1 old; Fig.8(J)). We infer that the DBA/2J mice search at both platform locations, though they prefer to search at the new platform location slightly more intensely. Similar to C57Bl/6J, the DBA/2J mice search at the most recent training location, although they never truly learnt the first platform location.

Since the purpose of this study was to determine whether training resulted in goal reversal, we defined a goal reversal efficiency measure (GRE) as rI_cs_ at the new location subtracted by the rI_cs_ at the old platform location divided by the sum of rI_cs_ at both the old and the new platform locations 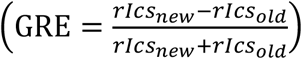 A ratio of 1 in this measure would indicate goal reversal, while -1 would indicate no reversal, and 0 would mean no learning. We find the goal reversal efficiency for C57Bl/6J to be ∼0.78 and DBA/2J to be∼0.15, indicating that C57Bl/6J are more adapt at goal reversal than DBA/2J mice.

## Discussion

Here, we introduce a new set of informative measures to describe spatial learning and memory in a navigational task based on time-stamped xy trajectories. The use of velocity-based vector fields and their properties allows the pin-pointing of putative search centers in the MWM. With these search centers, it is possible to describe spatial memory using three novel metrics: accuracy of search center to the target location, uncertainty of the search or spread in search area, and the intensity or extent of effort in search. We note that these three are independent of each other and are sufficient to quantify spatial learning and memory. To illustrate this, we consider a real-life navigational task of visiting a favourite café located block “A” between intersections “I” and “II”. In this context if the person remembers and goes to block “B” then *accuracy* captures the difference between the true location of the café and the actual location person went to. On reaching the place and not finding the café the person back tracks and searches around the place. The extent of area within which the person searches reflect their own estimate of how erroneous their memory for the location is. In this example the person knows the café is between two intersections but not remembering the block. *Uncertainty* captures this effect in our measure. Finally, the intensity of the search at block “B” captures the probability and hence the intention of that the person to target the search for the café at that point.

To illustrate the sensitivity and advantage of metrics introduced here, we used multiple MWM data sets. We show that our methods can detect subtle memory differences previously undetected using occupancy-based quadrant, platform, proximity, or entropy measures.

Moreover, we argue that the metrics introduced here reflect more closely spatial learning and memory than previous measures. For example, our methods reveal whether rodents use single or multiple search strategies in a trial, as well as the combined effect of the search results in swimming toward a particular region or location. This effect of swimming toward a given location is detected as a convergence peak or hotspot (i.e., the divergence field property is not preferentially sensitive or dependent on a particular search strategy). Additionally, since divergence measures the change in velocity in space, differences in absolute speed between strains of mice do not confound our spatial learning and memory measures, as they confound the interpretation of previous measures (i.e. the divergence field property can dissociate a memory deficit from a motor deficit). Differences in swim speed affect many of the conventional measures since speed affects quadrant occupancy and area covered during searches. We note that moderate differences in swim speed do not necessarily affect training since the mice are usually left in the pool until they reach the platform. Both fast as well as slow swimmers can reach the platform during training. On learning the platform location, the mice normally reach the platform in 1/10^th^ of the maximum training duration (usually 60 seconds). Thus, with our measures, only mice that are considerably slower (>10 times slower) fail to reach the platform during this time, and thus only in this rare case would slow motor performance affect our spatial learning and memory measures. On the other hand, conventional measures such as quadrant occupancy, platform crossings and proximity, are dependent on total occupancy, and therefore are affected by changes in swim speed. However, the divergence of vector field measure introduced here reflects how the vectors are changing in space rather than the magnitude of the vectors per se. This minimizes, if not eliminates, the effects of difference in swim speeds. Thus, the proposed metrics, based on divergence of vector fields, improve the sensitivity of measuring memory performance by uncoupling the confounding effects of non-memory related processes, such as swimming speed, from measures of spatial learning and memory.

While we have presented the description, derivation, and potential utility of both the divergence and curl vector field properties, here we focused our analyses on developing and highlighting the power of the metrics derived from divergence vector field properties.

In summary, analysing swim trajectories using the proposed metrics enabled us to detect subtle memory deficits in four very different MWM datasets. We show a difference in the strength of spatial learning and memory for two different mice strains subjected to the same training paradigm (BALB/cJ and SWR/J). We also analysed two different mouse models of Noonan syndrome, including *Ptpn11*^D61G/+^ mice that were previously thought to lack spatial learning and memory even after extended training. Our analyses revealed that *Ptpn11*^D61G/+^ mice do indeed show evidence of learning the platform location after extended training, and that their accuracy and uncertainty in the search area on PD5 is comparable to that of their WT littermates on PD3. We also show that the *Ptpn11*^N308D/+^ mouse model of Noonan syndrome have a more subtle spatial learning and memory deficit in the MWM that was not easily detected by conventional measures. Additionally, treatment with a MEK inhibitor rescued the learning and memory deficits in *Ptpn11*^N308D/+^ mice, adding to the evidence that suggest that inhibitors of this kinase could be used to treat cognitive deficits in patients. Lastly, we investigated whether the divergence field property on the velocity-based measure could detect the presence of multiple search centers, reflecting multiple spatial memories, in a MWM goal reversal data set. The two strains of mice (C57Bl/6J and DBA/2J) trained in this goal reverse task revealed the presence of two search centers, corresponding to the old as well as the new target platform locations. Quantification of the difference in strength or nature of these two spatial memories, in terms of the extent of intention of search at the two locations or search centers, showed that both strains of mice prefer to search at the most recently trained platform location, and that acquisition of the new memory for the platform location did not disrupt memory for the first location. All together our studies demonstrate that the measures introduced here are capable of extracting new information from readily accessible analyses of performance in the MWM, and consequently that they are capable of yielding novel insights that would be otherwise overlook by older measures of this widely used task for spatial learning and memory.

## Appendix: Theory

### Velocity-based vector fields for describing the intentional movement of rodent swim trajectories

i) Description of velocity vector fields

We first developed an analytical framework for describing the swim trajectories in terms of a velocity vector field. The position vector 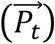 represents the point at which the rodent is located at a given time (‘t’-usually measured in video frames). The origin of the reference frame *O*(0,0) is set to be the top left corner (Fig.1 (A)). Thus, from an image taken from the top, the pixel co-ordinates (P_x_, P_y_) can be used to write (P_t_) as follows:

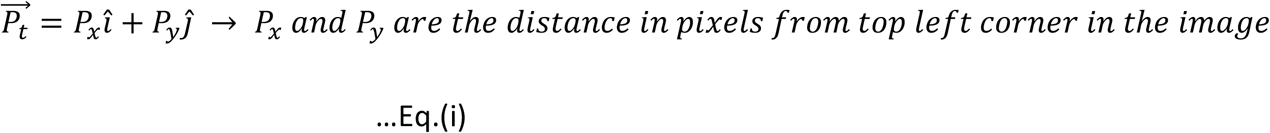

For convenience, we define a displacement vector 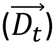 that provides a relation of the rodent’s position with respect to a point of interest, e.g., the platform. This is calculated as the difference between the platform vector 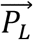 and the instantaneous position vector 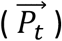 defined above. (Fig. 1 (B)).

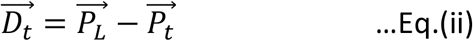

The velocity vector 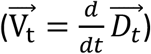 is a metric that describes the movement of the rodent. Since we make these measurements from time lapse videos that are recorded at uniform frame rates, we can calculate the velocity vector as the difference between two consecutive displacement vectors. (Fig. 1 (C)).

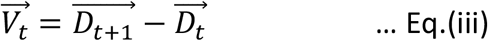

ii) Definition of occupancy center

Since we measure the component of velocity of the animal that is directed towards reaching the “goal”, it is convenient to define the occupancy center P_oc_ as the point where the likelihood of finding the animal is high given its trajectory. We calculate P_oc_ as the center of mass obtained from the residence time map that is segmented using a maximum entropy threshold. Maximum entropy (ME) segmentation determines a threshold (T) that has the maximum entropy of the background (H_B_) and foreground (H_F_) in an image (Sahoo et al., 1988).

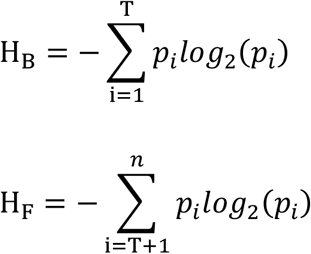

where *p_i_* is the number of pixels with value *i* divided by the total number of pixels, and *n* is the maximal possible intensity in the image. This enables us to segment out the regions with zero or low occupancy from regions of high occupancy without bias. The use of ME segmentation increases the fidelity and hence the confidence in our estimate of the occupancy center as the most likely resided region based on a rodent’s swim trajectory. This is given by (Fig. 1(F))

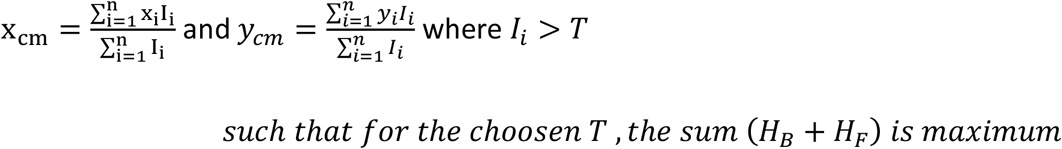

iii) Description of vector fields with respect to the occupancy center

Using the occupancy center, we computed two different measures to describe the rodent’s movement. First, the component of velocity vector along the occupancy center, a spatial property that measures the contribution of the movement at that location towards the occupancy center. This component is the vector projection or the velocity vector *V(t)* along the direction of the occupancy center 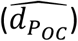, and is given by (Fig. 1(D))

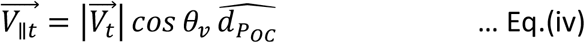

Where 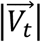 is the magnitude of the velocity vector and *θ*_*v*_ is the angle between the velocity vector 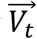 and a unit vector pointing towards the occupancy center 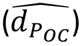 from the current position.

Second, the velocity vector component orthogonal to the occupancy center is a spatial property that measures at that position, the contribution of the rodent’s movement to the circular motion about the occupancy center. It is given by (Fig. 1(E))

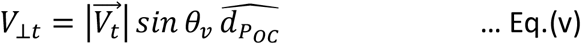

where 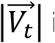 is the magnitude of the velocity vector and *θ*_*v*_ is the angle between velocity vector 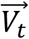 and the unit vector pointing towards the occupancy center 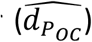 We note if ME segmentation yields multiple disconnected regions of comparable sizes then we need to consider multiple occupancy centers. In such cases we evaluate the velocity vector components towards each of them and take the weighted sum of their magnitudes. The weights are determined by the fractional area of the patch. In the current study we did not use such as approach.

Next, we resolved these vectors Eq.(iv) and Eq.(v) in a frame of reference defined with respect to the image of the pool. The origin is set at the top left corner of the image with right and down being the positive directions of its axis. Thus, the above vectors could be written as

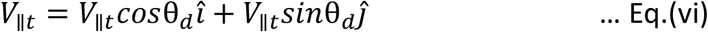

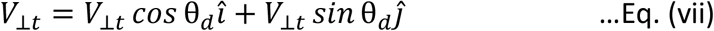

where θ_d_ is the angle, *î* and *ĵ* are the unit vectors along the height and width of the image frame.

### Analytical expression of the putative search centers obtained from divergence in velocity-based vector fields

i) Vector field properties

Divergence of a vector field indicates the difference or change in flux around an infinitesimal volume. We reason that given a vector field *V*_‖*t*_, the point of convergence (“sink” in a vector field) could represent the point where the animal intends to reach. We hypothesize that if a rodent intents to reach a specific location 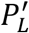 in the pool, then the velocity field would point inward from the region with progressively decreasing magnitude (as described in the introduction) around the point 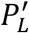 leading to a negative divergence value (i.e., indicating convergences) at 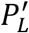 However, in general 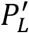 need not necessarily coincide with the platform location or occupancy center, for they represent the center of a region where the rodent is most likely found, and not the actual platform location. We construct the vector field from vector projections (Eq. 1) to measure and represent the intention of the animal to move towards the occupancy center as a function of its position in the pool. The heat map of the global minima of such a divergence would be indicative of the platform location as perceived by the animal. We calculate the divergence on the vector field describing the velocity along the occupancy center in the image reference frame as

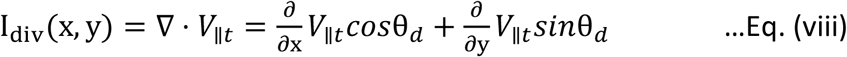

Similarly, we reason that the curl of a vector field could indicate the tendency of rotation of an object placed at that location, and hence we propose that it could be used to extract the tendency of the animal to circle around any point in the pool. For completion, we describe the curl, but in subsequent analyses, we restricted our analyses to divergence.

Curl of a field is formally defined as the circulation density at each point of the field (i.e., the extent of rotation about a point). Thus, curl on *V*_⊥*t*_ vector field would be informative of the extent of circular movement that the animal makes about the perceived platform location while searching for it. Using our notation, the curl can be written as:

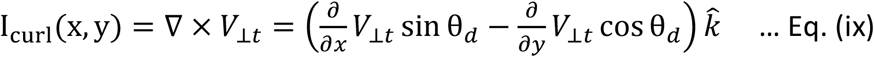

ii) Description of the putative search center (*P_cs_*)

Search centers can be obtained from the divergence calculated on the vector field of the velocity component pointing towards the occupancy center. The convergence peaks in such a map are indicative of the search centers 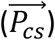 or the platform locations as perceived by the rodent.

We obtain the search center or the convergence point by estimating the minima of the divergence on the velocity-based measure. For a two-dimensional surface, we obtain the critical point ((*x*_*0*_, *y*_*0*_)) by equating the first derivative to be zero and forcing the second derivative to be positive as follows:

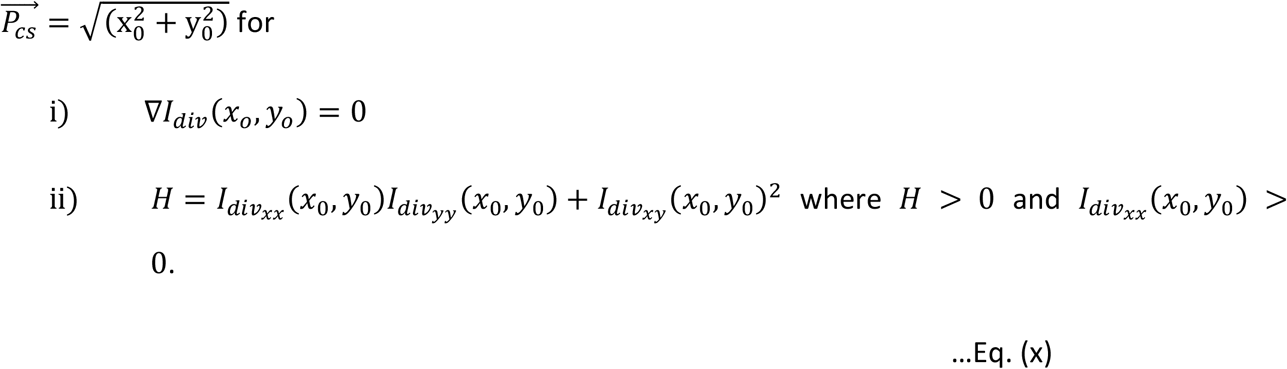

#### Metrics to assess spatial memory

We define and use three measures that are independent of each other to assess spatial memory in terms of various attributes of the search center 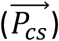. The three independent measures namely, including accuracy of search, uncertainty about the search center, and intensity at the search center, are described next.

i) Accuracy of the search center (α_cs_) (Fig. 1(G))

Since the search center is indicative of the animal’s perceived platform location, the distance between the search center and the platform location measures inaccuracy or error in the animal’s ability to learn and remember the platform location. The difference in location measured in units of length can be difficult to compare across different laboratories, since the effective pool and platform sizes can vary between laboratories. To address this problem, we developed a percentage measure that can be used for comparisons of animals’ performances across laboratories. First, we realized that the maximum error in accuracy that the animal can make can be captured by the distance between the platform center and the farthest pool boundary (*e*). Next, we defined the accuracy of search (α_cs_) as 1 minus the fractional error, with fractional error being the ratio between the distance of the search center from the platform and the maximum error. Thus, the accuracy of the search center is given by

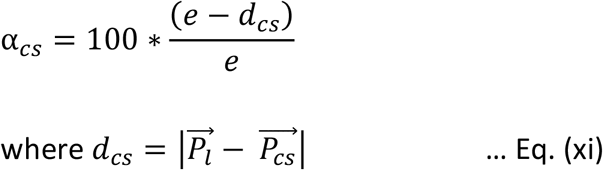

Furthermore, to estimate the chance factor we consider the case of a naïve animal that does not have any knowledge of the platform location (e.g., an animal on its first day of training). At this point of training, animals usually exhibit wall-hugging behaviour (i.e., they swim in circles close to the wall). In such a scenario, the animal is moving about the pool center as its navigational center, and by virtue of symmetry the pool center would reflect the search center. We make use of this fact to define the chance accuracy as the accuracy when the search center coincides with the pool center and is given by

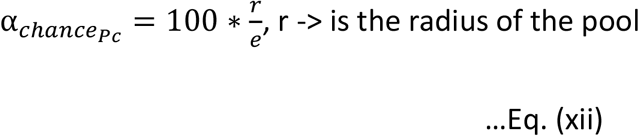

Thus, an accuracy value that is comparable to the chance factor (Eq. x) would reflect whether the animal is still exhibiting symmetric navigational behaviour. As the rodent learns the rules of the water maze, this symmetry is broken. Comparisons of accuracy with the chance factor above yield a categorical measure of learning and memory in the maze.

ii) Uncertainty about the search center (σ_*cs*_) (Fig. 1(H))

Apart from the location of the search center, the spread of the divergence values around the peak provide additional insights into the level of uncertainty in the internal representation of the platform position in the animal’s performance. We calculated this by measuring Uncertainty about the search center and defined it as the full width half maximum (FWHM) of the convergence peak representing the search center. We assume that the convergence peak is a 2D Gaussian, and compute the FWHM by linearising the 2D Gaussian equation as previously described (*2D Gaussian fitting macro (Fiji/ImageJ) for multiple signals. | BIII*, n.d.). We express the uncertainty about the search center in terms of the relative search diameter, defined as a ratio of the major axis of the Gaussian ellipse to the platform diameter.

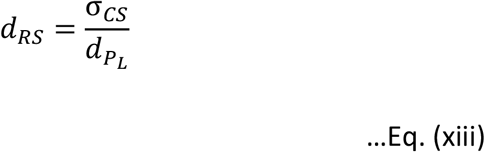

iii) Intensity of search (*I*_*cs*_) (Fig. 1(I))

The absolute search intensity (aI_cs_), defined as the magnitude of convergence at the search center, is a measure of the animal’s searches in the search center, i.e.,

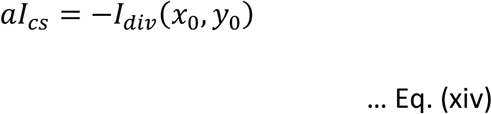

Additionally, we define the Relative search intensity (rI_cs_) as the magnitude of convergence at the search center normalised to the maximum convergence seen anywhere in the pool. We propose that this measure reflects the fractional effort of the animal’s search at the search center.

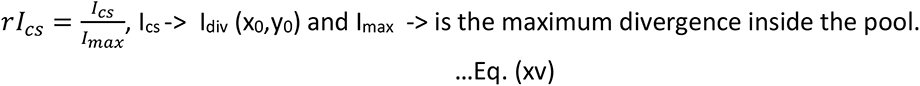

### Materials and Methods

#### Animal information

BALB/cJ (Stock no: 000651) and SWR/J (SWR/J Stock no: 000689) inbred mice were obtained from Jackson Laboratory, USA and maintained at the Central Animal Facility, IISc. All protocols were approved by the Institute’s Animal Ethics Committee.

For the experiments using Noonan Syndrome (NS) mice, we used both transgenic and viral-induced mice models. NS virus-induced mice models were created by the induction of adeno-associated virus (AAV)–mediated expression of the NS-associated mutation (*PTPN11^D61G^*) through bilateral injection into the dorsal CA regions of the hippocampus of 3-month old C57Bl/6J male mice. To construct pAAV-hSyn-*PTPN11^D61G^*-HA-T2A-tdTomato plasmid, *PTPN11^D61G^*-HA-T2A-tdTomato was synthesized by using recombinant PCR and inserted into the AAV vector pAAV-hSyn-eGFP.WPRE.bGH vector (Addgene #105539), replacing eGFP between NheI and HindIII site.

The NS mice were trained as reported previously (Lee et al., 2014). Briefly, training sessions comprised 4 trials (2 blocks with 2 trials each) per day, with 1-min intertrial interval and 45-min interblock interval. Animals were allowed to search the platform for 60 sec or until reaching the platform, as shown in the original paper. The probe tests were done immediately after the completion of training on days 3, 5, 7, or 11, depending on the experiment. For the experiments with MEK inhibitor, SL327 (32mg/kg, DMSO) was injected intraperitonially everyday 30 min before MWM training.

#### Generation of velocity-based vector field via surface fit

We describe the swim trajectories in terms of the velocity component along the occupancy center for all pixels sampled by the mice. The value at each pixel is normalised to the number of times it has been sampled (i.e., the mean value at the pixel).

We perform a 5^th^ order polynomial surface fit (ImageJ Polynomial Surface Fit plugin) using an ROI of sampled pixels. The resultant surface is a continuous surface representing the velocity-based vector field. We require that the fit interpolates values for unsampled pixels within the sampled pixels ROI to compute the partial differential along the horizontal and vertical axis of the pool reference frame. Since the regions outside the ROI are not sampled, we do not consider it to be a part of the vector field. We modified and used the Differentials Plugin (Unser, 1999) in ImageJ to obtain the components of divergence.

#### Estimation of error in generated velocity-based vector field

The error in 5^th^ order polynomial surface fit is described by estimating the goodness of surface fit. The reduced chi-sq 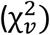 describes the variance in the velocity-based measure among the population of mice as well as the variance in the estimate of fit values at each pixel.

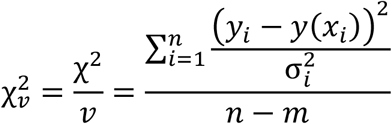

where *y_i_* is the actual value at the pixel *i*, *y(x_i_)* is the fit value at the pixel *i*, 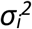 is the variance in velocity-based measure value at the pixel *i*. *v* is the degree of freedom defined as the difference between the number of points sampled on the surface (*n*) minus the number of fitted parameters (*m*). In case of a 5^th^ order polynomial equation, there are 14 coefficients or fitted parameters, hence the degree of freedom *v* = *n – 14*.

#### Estimation of error in field properties

We represent the estimated error in the field properties as a percent error in residual of the surface fit.

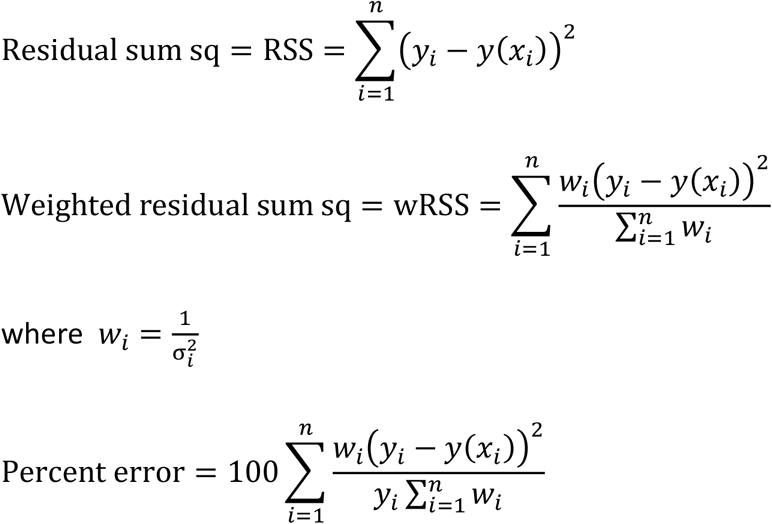

where *y_i_* is the actual value at the pixel *i*, *y(x_i_)* is the fit value at the pixel *i*, 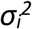 is the variance in velocity-based measure value at the pixel *i*.

Hence, we calculate the total error at each pixel of the field property as the sum of the fractional errors contributed by the x- and y-components with respect to the pool reference frame, given by,

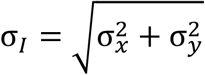

We utilize the mode of the distribution of error values to best represent the most likely error estimate. We propagate this mode value of error when representing the derived parameters/metrics, i.e., accuracy of search, uncertainty of search and intensity of search

#### Identification of putative search centers

We identify the putative search center as the global minima of the divergence heat map. The obtained divergence heat map is separated into two images, each containing either only positive values (divergence values) or negative values (convergence values). The image with negative values is inverted to convert the minima to maxima for visual representation. Next, we normalize the intensity value to the maximum intensity value. We locate the maxima using the ImageJ’s plugin Maximum Finder class. The algorithm is set such that a maximum is identified if and only if the point has a peak value difference that is more than 0.1% and that the point is not an edge.

#### Estimation of full width half maximum (FWHM) of convergence peak

We estimate the FWHM of a convergence peak as previously described (*2D Gaussian fitting macro (Fiji/ImageJ) for multiple signals. | BIII*, n.d.). Briefly, the identified putative search center is fit to a linearised 2D-gaussian function to obtain the FWHM in x- and y-axis. The size of the peak is initialised to 20 pixels before performing custom fit using ImageJ’s Curve Fitter class implementing Simplex algorithm.

#### Statistical analysis

All statistical analysis was carried out in Origin (v2020b or 2021b). To perform two-way ANOVA on summarised statistics (i.e., ANOVA using mean and SEM), the raw data was recreated for ‘n’ individual mice by randomly sampling from a normal distribution using the population mean and SD of the metric being analysed. Based on ANOVA results, post hoc analysis was performed to carry out pair-wise comparisons. The data distribution was assumed to be normal. All data are represented as mean ± SEM.

## List of abbreviations

MWM: Morris water maze
ROI: Region of interest
ME: Maximum entropy

## Declarations

### Ethics approval and consent to participate

All protocols were approved by the Institute Animal Ethics Committee.

### Consent for publication

Not applicable.

### Availability of data and materials

The datasets used and/or analysed in the current study are available after publication from the corresponding author on written request. The software, workflow along with the codes developed will be made available in the github (https://github.com/neurodynamicslab).

### Author Contributions

M.P, D.M., N.N. and B.J. designed the water maze experiment for BALB/cJ and SWR/J mice. D.M. and N.N. collected the water maze data for BALB/cJ and SWR/J mice. Y.S.L. and A.J.S designed the SL327 rescue experiments with *Ptpn11* N308D/+ mice and provided the Noonan mouse model data. Y.S.L and D.A.F. performed the Noonan mouse model experiments in A.J.S.’s lab. M.P. and B.J. developed the theory, wrote the program for data extraction and analysis, analysed the data, and authored the manuscript. Y.S.L, D.A.F and A.J.S read and provided corrections for the manuscript.

## Acknowledgments

The authors thank the funding source for their support and Prof. Kempermann and Prof. Rupert Overall, CRTD, Dresden for sharing the C57Bl/6J and DBA/2J data, Dr. Hyun-Hee Ryu for AAV vector construction. We also thank Prof. Chandra Murthy, IISc for reading through the manuscript and providing his valuable comments on the manuscript. We especially acknowledge his suggestion for estimating the field when there are multiple isolated areas of residence.

## Supplementary Materials

**Supplementary figure 1:**
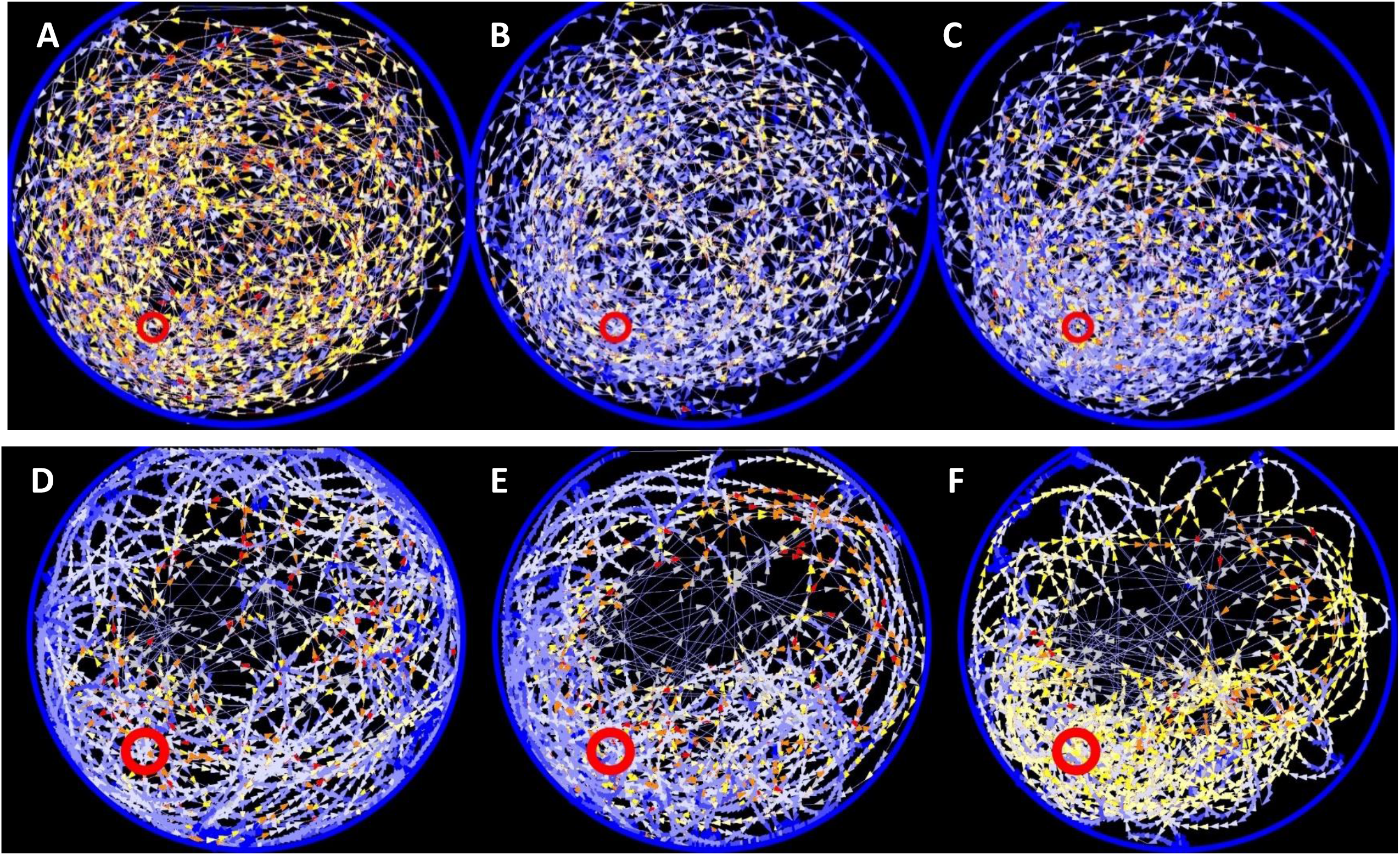
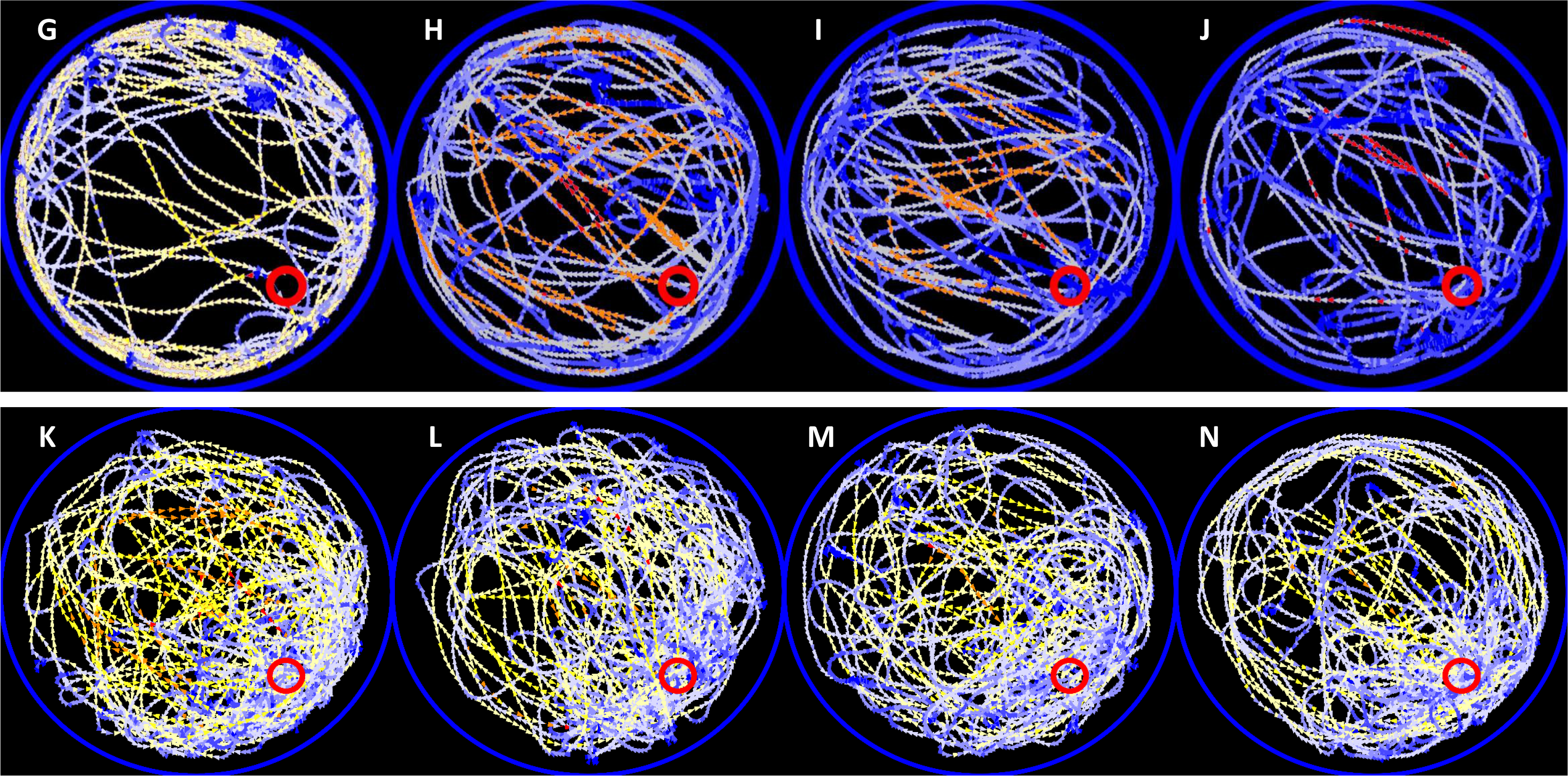
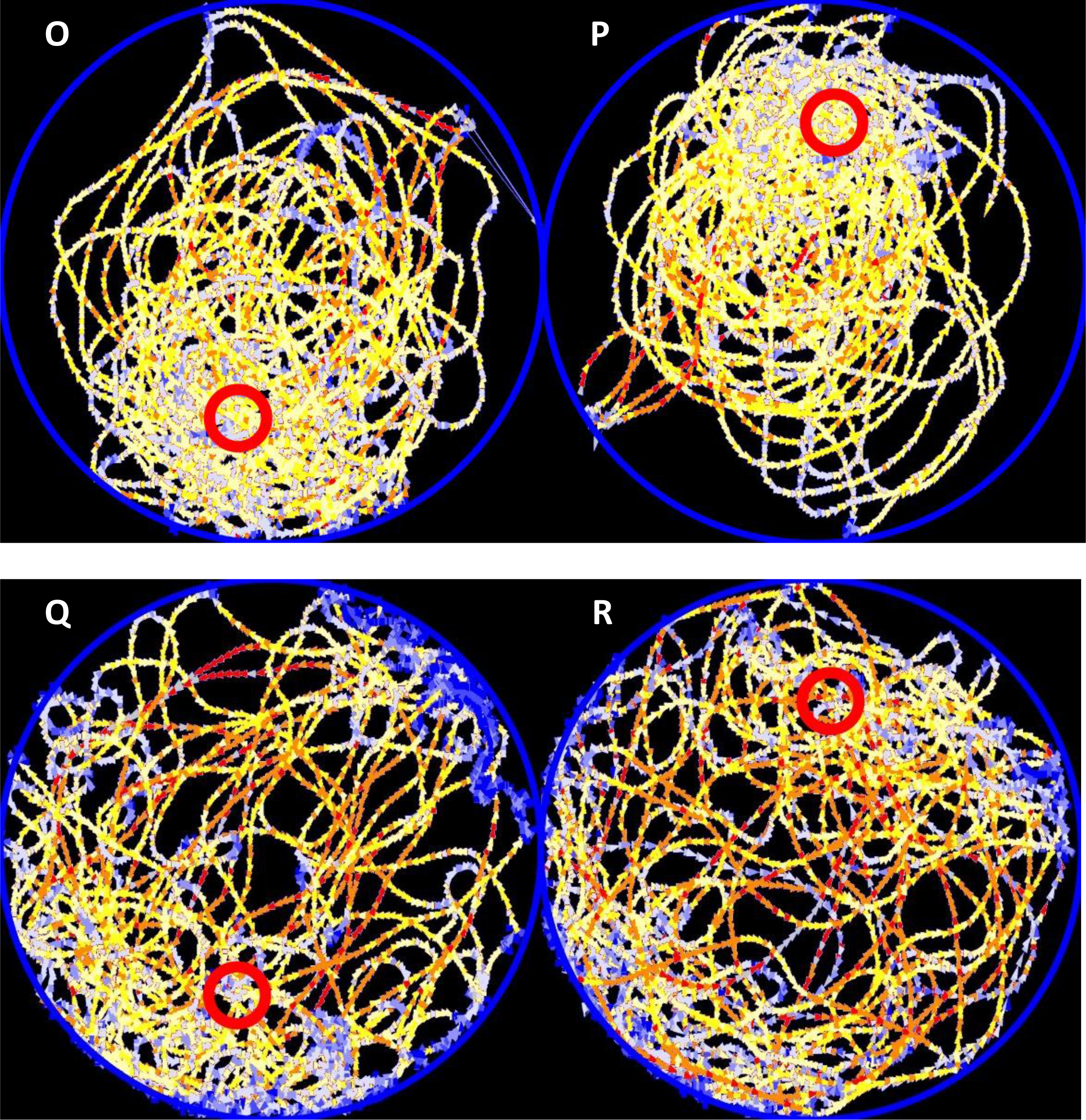
Velocity vector field map obtained from the population of mice used in each dataset. The pool perimeter is indicated as a blue ROI and the platform location is marked as a red ROI. A-C: BALB/cJ, D-F: SWR/J, G-J: *Ptpn11* D61G/+, K: *Ptpn11* +/+ Veh, L: *Ptpn11* +/+ SL327, M: *Ptpn11* N308D/+ Veh, N: *Ptpn11* N308D/+ SL327, O-P: C57Bl/6J, Q-R: DBA/2J. LUT scale: D61G 0 – 8, BALB/cJ 0 - 100, SWR/J 0 - 50, C57Bl/6J and DBA/2J 0 – 5 all in pixels/frame.

**Supplementary figure 2:**
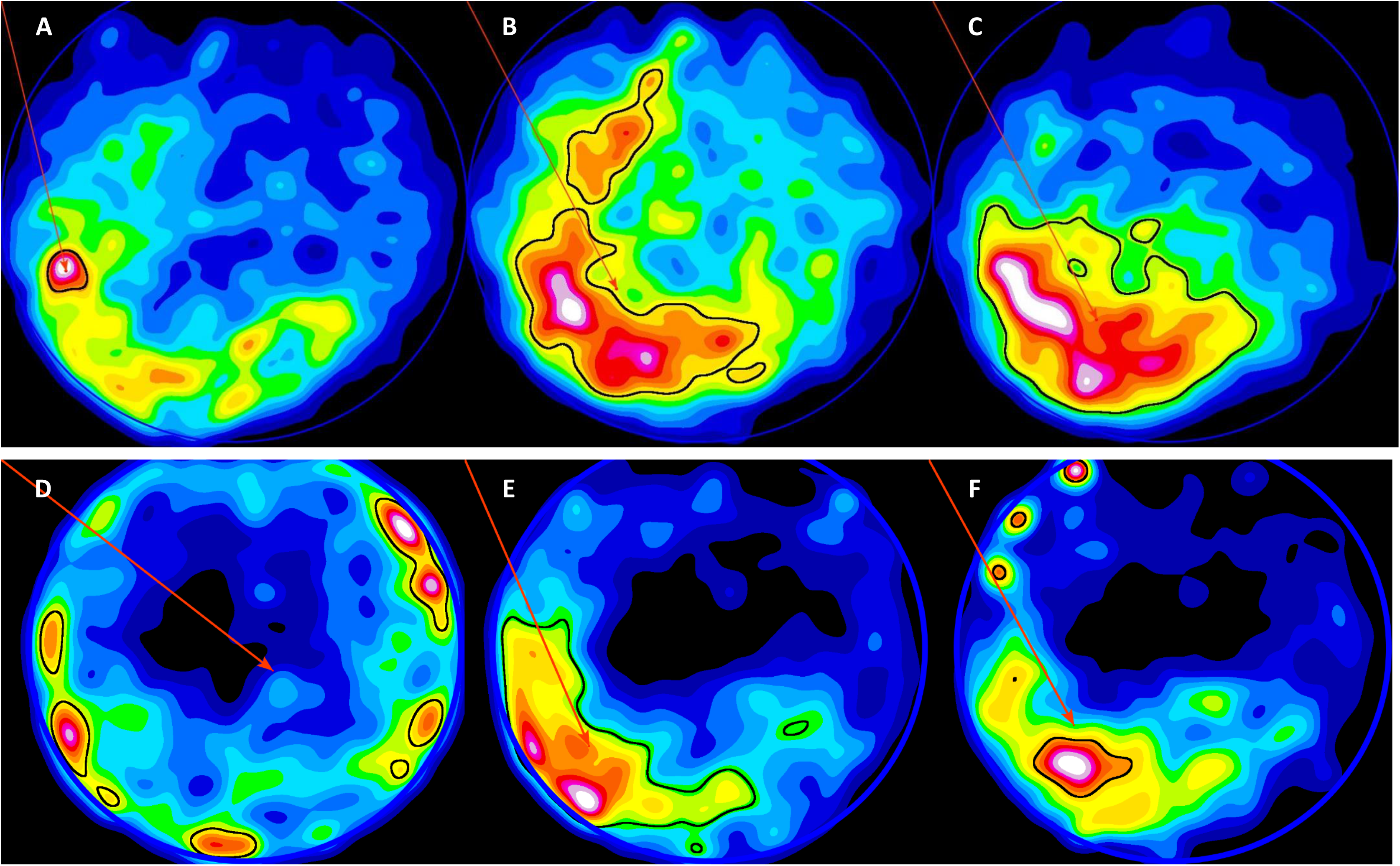
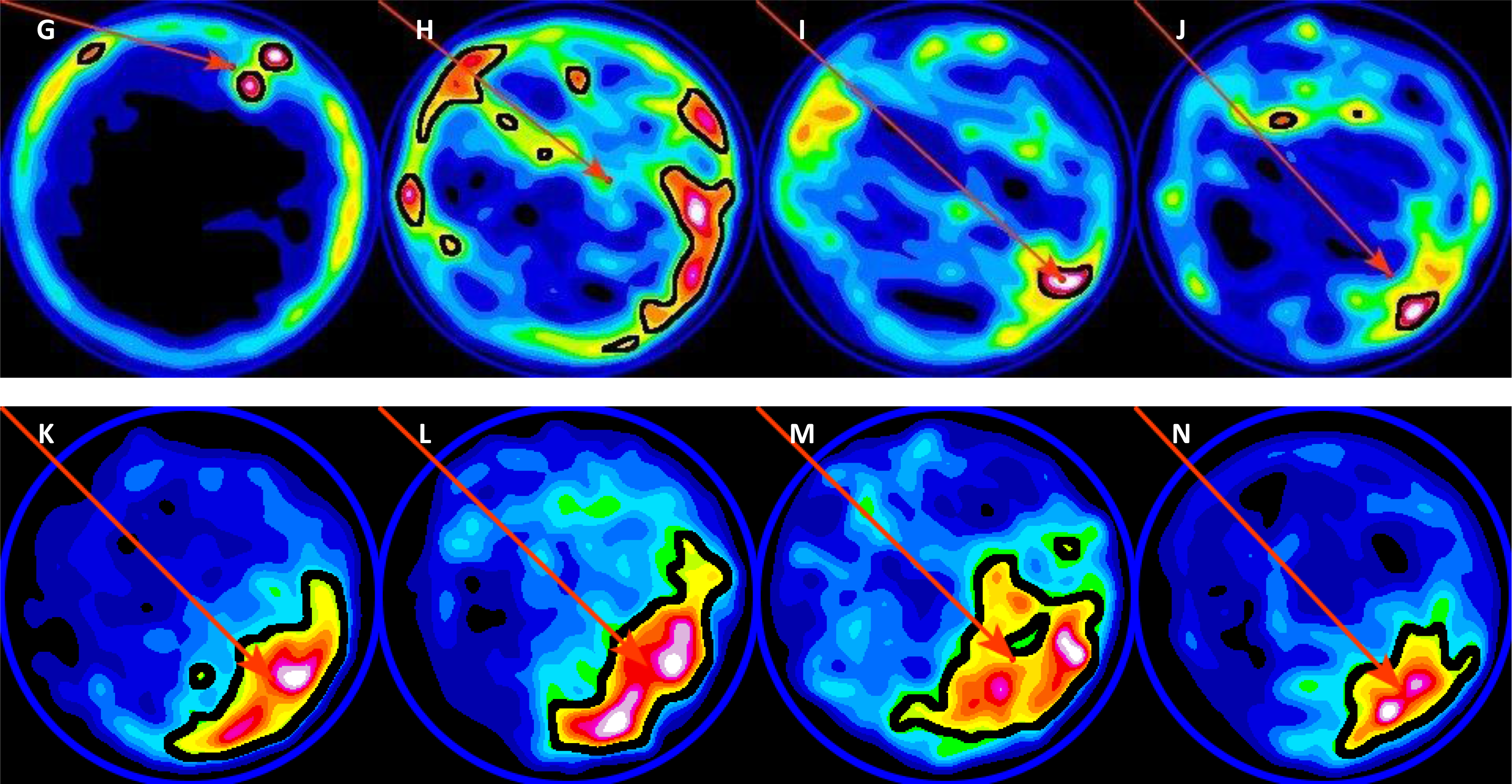
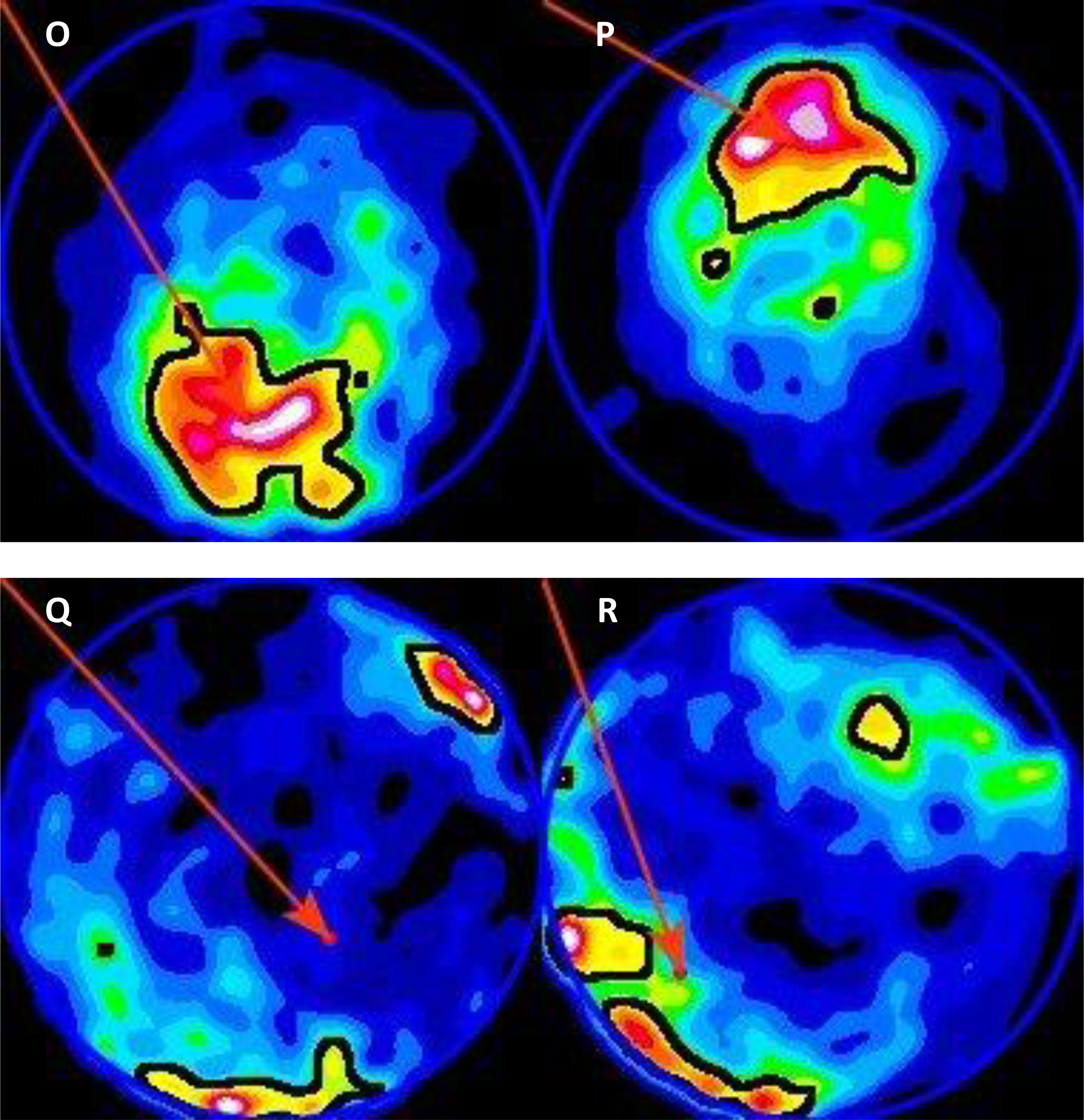
**Estimating occupancy centre using residence time heat map and centre of mass.** Occupancy heat map for population of mice used for each dataset. Gaussian blur was applied to the image to approximate the point object to real world dimensions (Gaussian blur value details provided in S. Table 1). Occupancy was thresholded using maximum entropy algorithm in ImageJ to obtain region of most likely occupancy (marked as a black outlined ROI), on which the centre of mass was estimated to obtain the occupancy centre (marked as a red dot with a position vector indicating the location). The pool perimeter is indicated as a blue ROI and the platform location is marked as a red ROI. Higher residence times are represented by warmer colours. A-C: BALB/cJ, D-F: SWR/J, G-J: *Ptpn11* D61G/+, K: *Ptpn11* +/+ Veh, L: *Ptpn11* +/+ SL327, M: *Ptpn11* N308D/+ Veh, N: *Ptpn11* N308D/+ SL327, O-P: C57Bl/6J, Q-R: DBA/2J.

**Supplementary figure 3:**
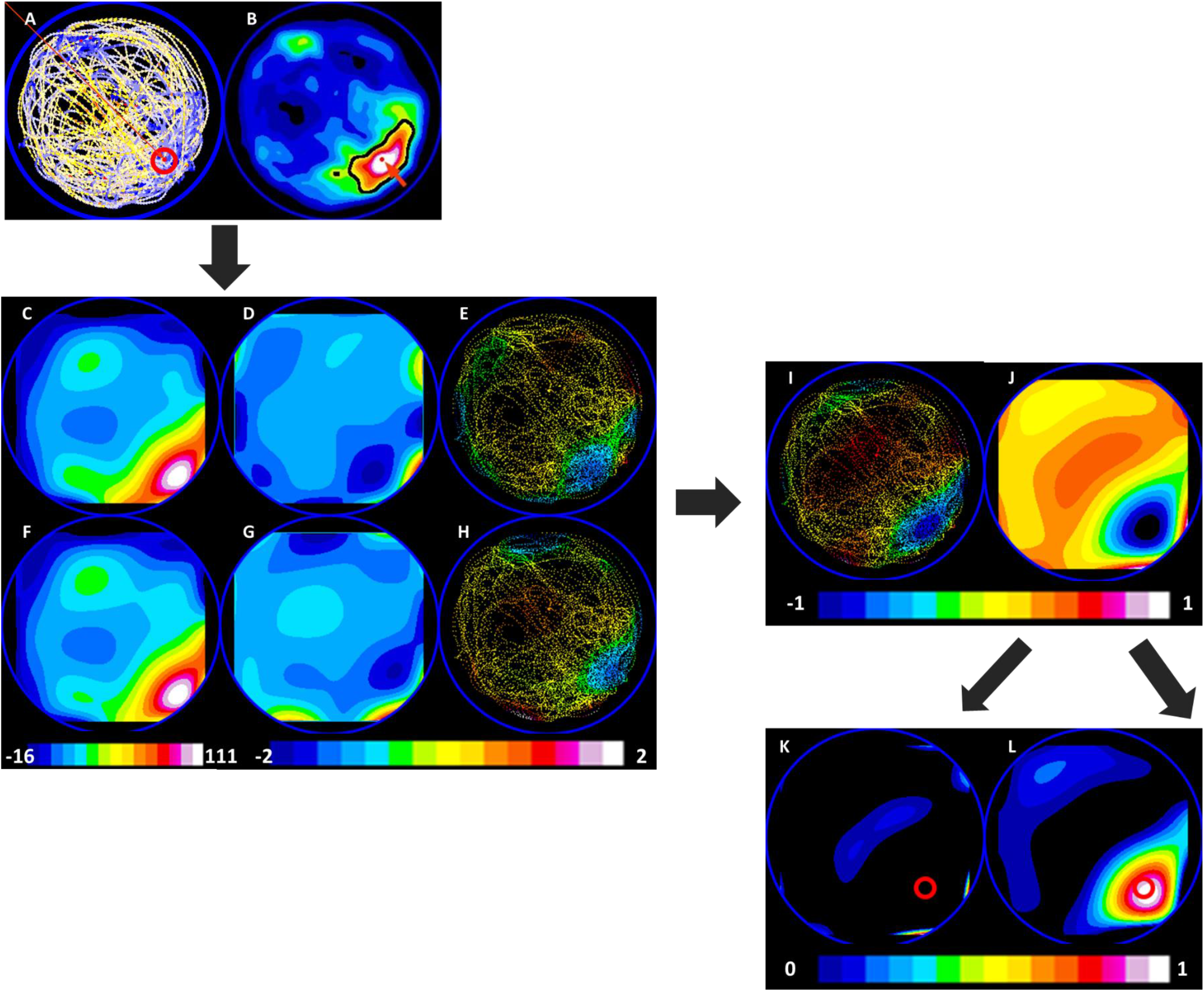
**Workflow to generate divergence heat map revealing convergence hotspots.** (A) shows the population velocity vector field (n = 15 mice, data points = 8997 frames). (B) shows the occupancy heat map. Using maximum entropy auto thresholding, the area for estimating COM of occupancy is determined (demarcated in black ROI). Occupancy centre is marked as a red dot. The red arrow indicates the position vector of the occupancy centre. (C) and (F)Using the velocity vector field and the occupancy centre, velocity vector component along the Poc (*V*_‖_) was generated for spatially resolved components of the velocity vector, i.e. x-Image and y-Image. The x-Image (C) is the component along the horizontal axis, whereas the y-Image (F) is the component along the vertical axis. An ROI encompassing the pixels that have been sampled is used to obtain an image that is a 5^th^ order polynomial surface fit (D and G), thus obtaining a surface describing the velocity-based movement towards the occupancy centre. The change in measure along the horizontal or vertical axis is obtained by taking the differential with respect to x-axis or y-axis. We maintain the calculated values in the pixels that were sampled. (E)shows the differential with respect to x-axis for the x-Image, and (H)shows the differential with respect to y-axis for the y-Image, for the pixels sampled by the mice. Adding the differential images (E) and (H) gives the divergence map as shown in (I). A smooth surface (J) representing the divergence values continuously in space is obtained after a 4^th^ Order surface fit on (I). The divergence map (J) is split into an image with only positive values to represent the points of divergence (K), and only negative values to represent the convergence hotspots (L).

**Supplementary Figure S4:**
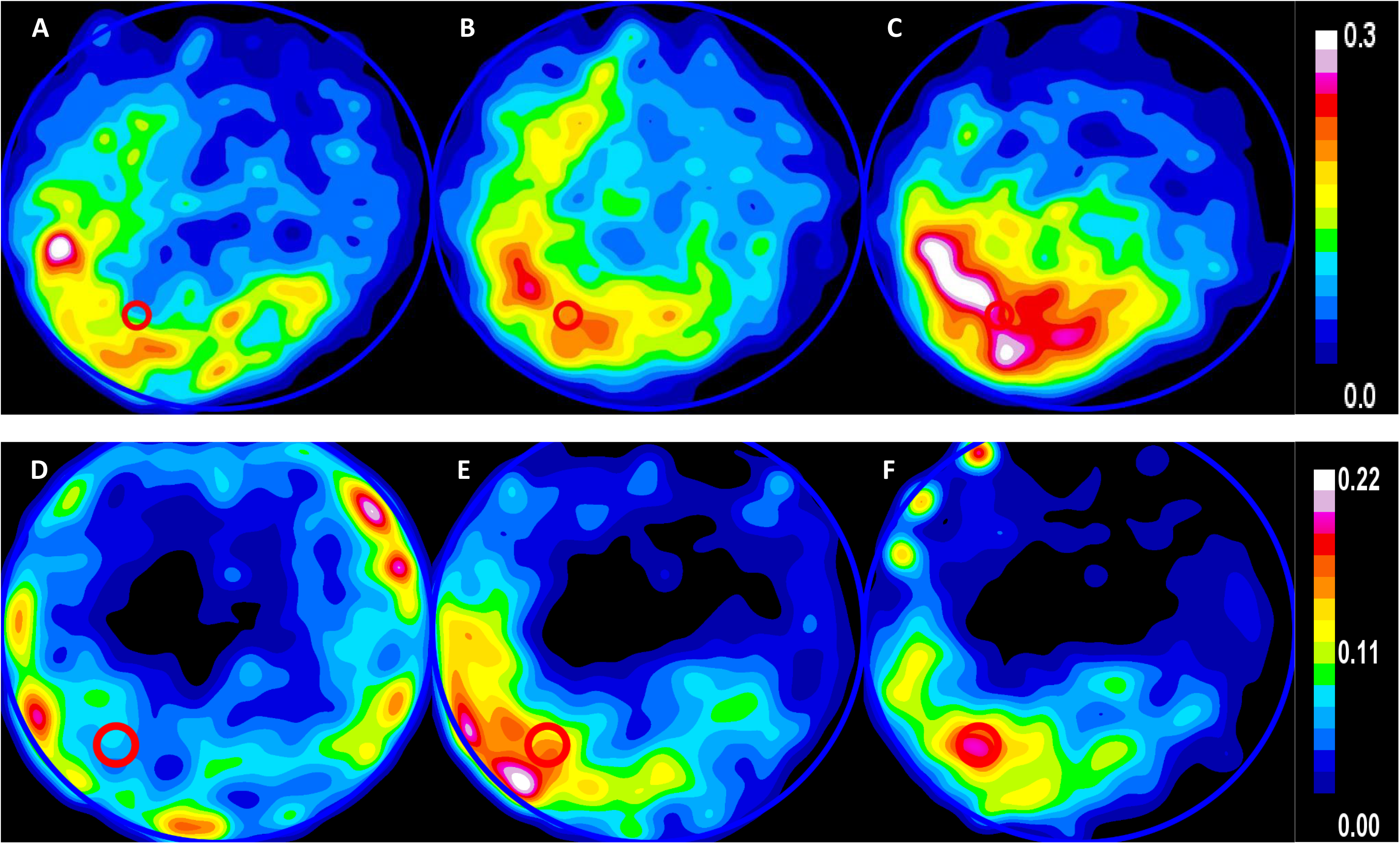
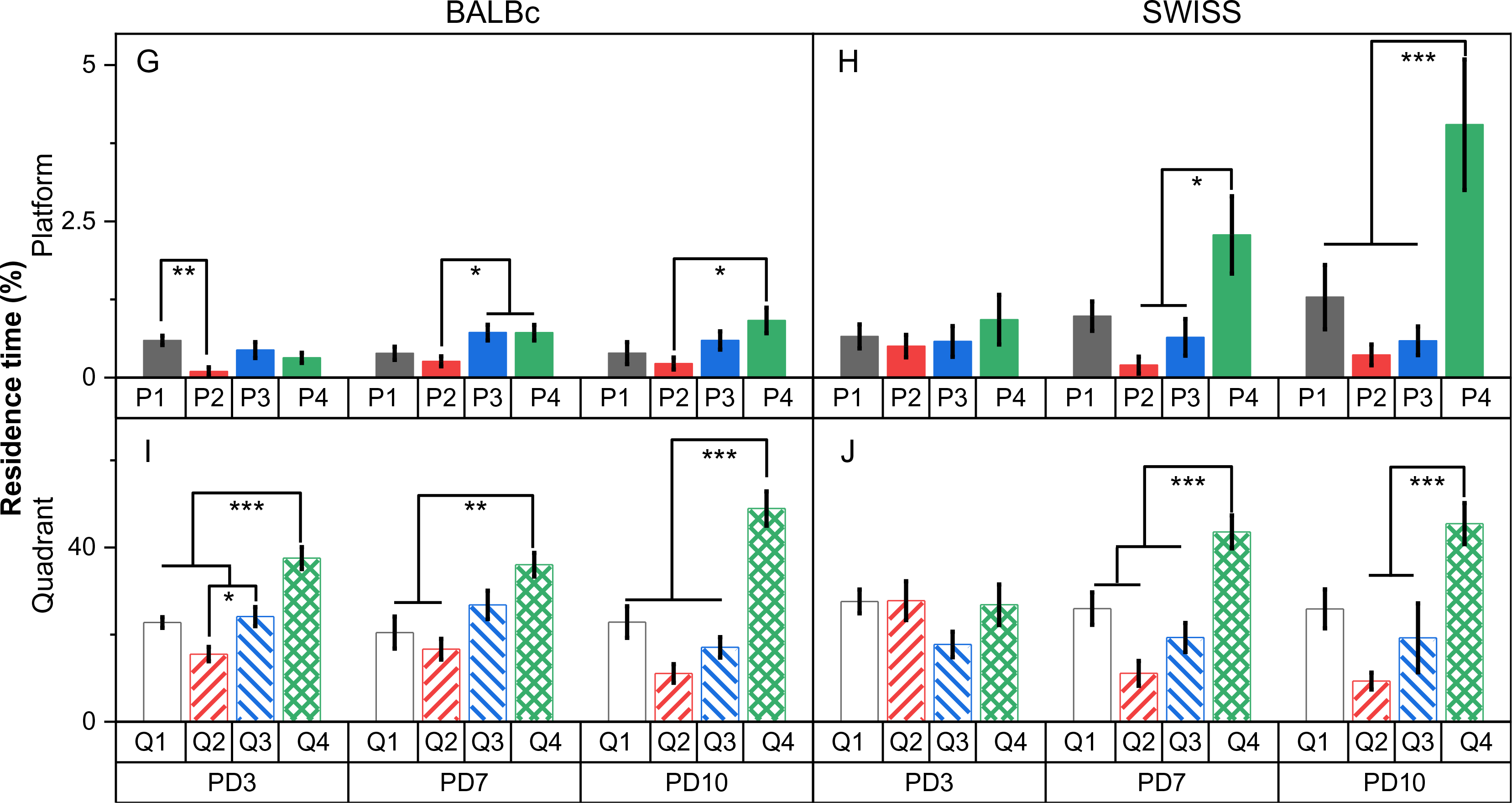
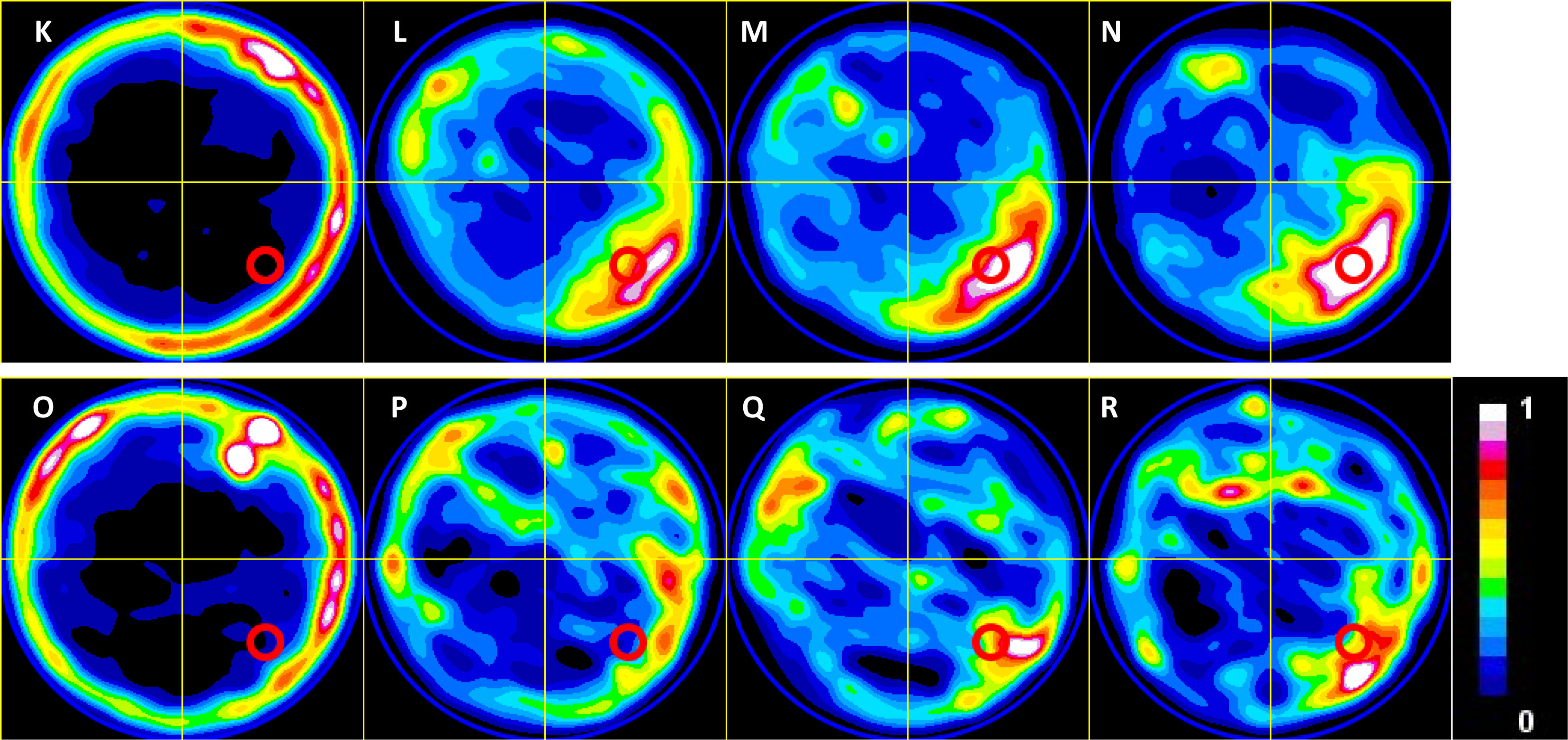
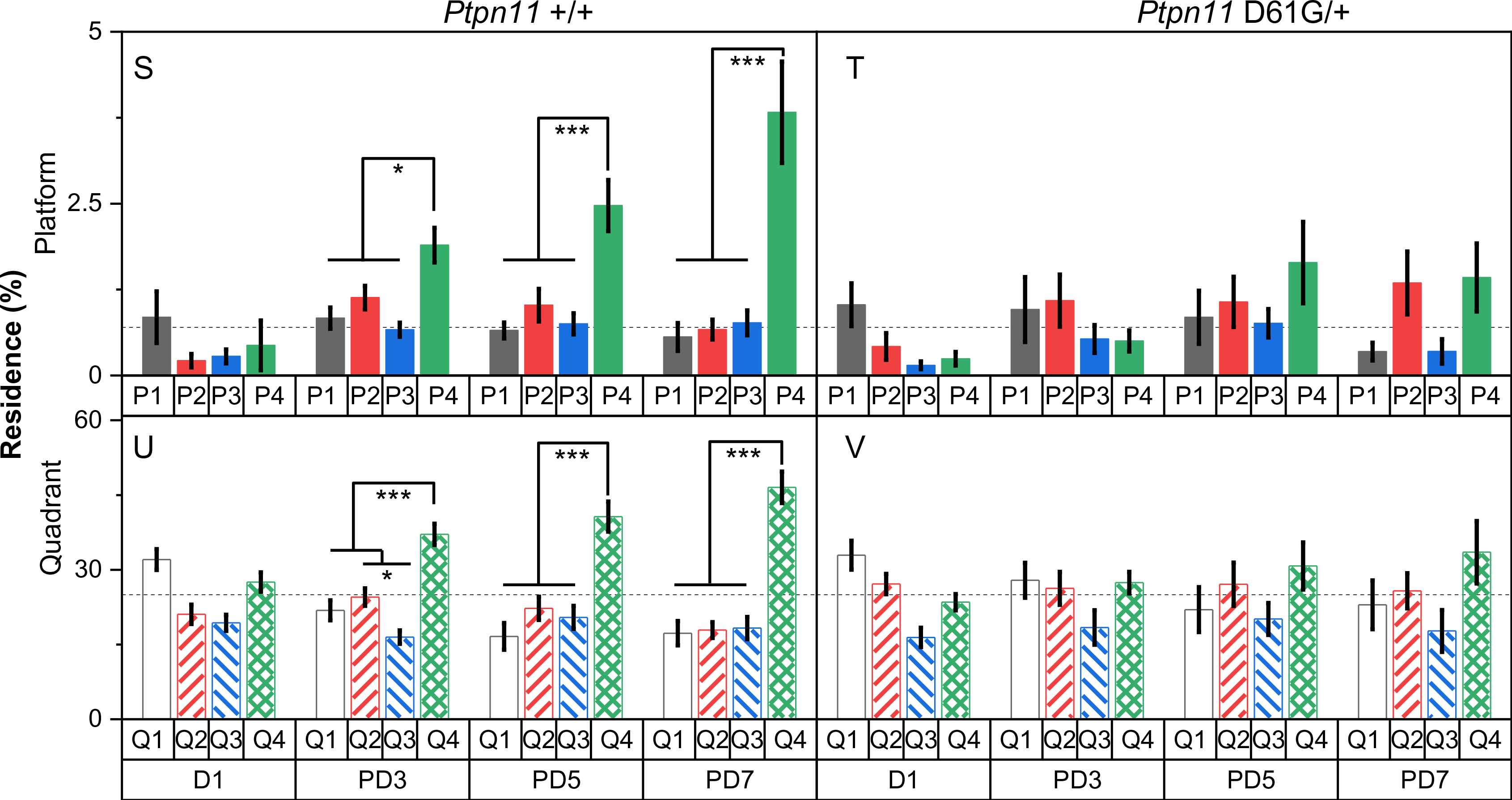
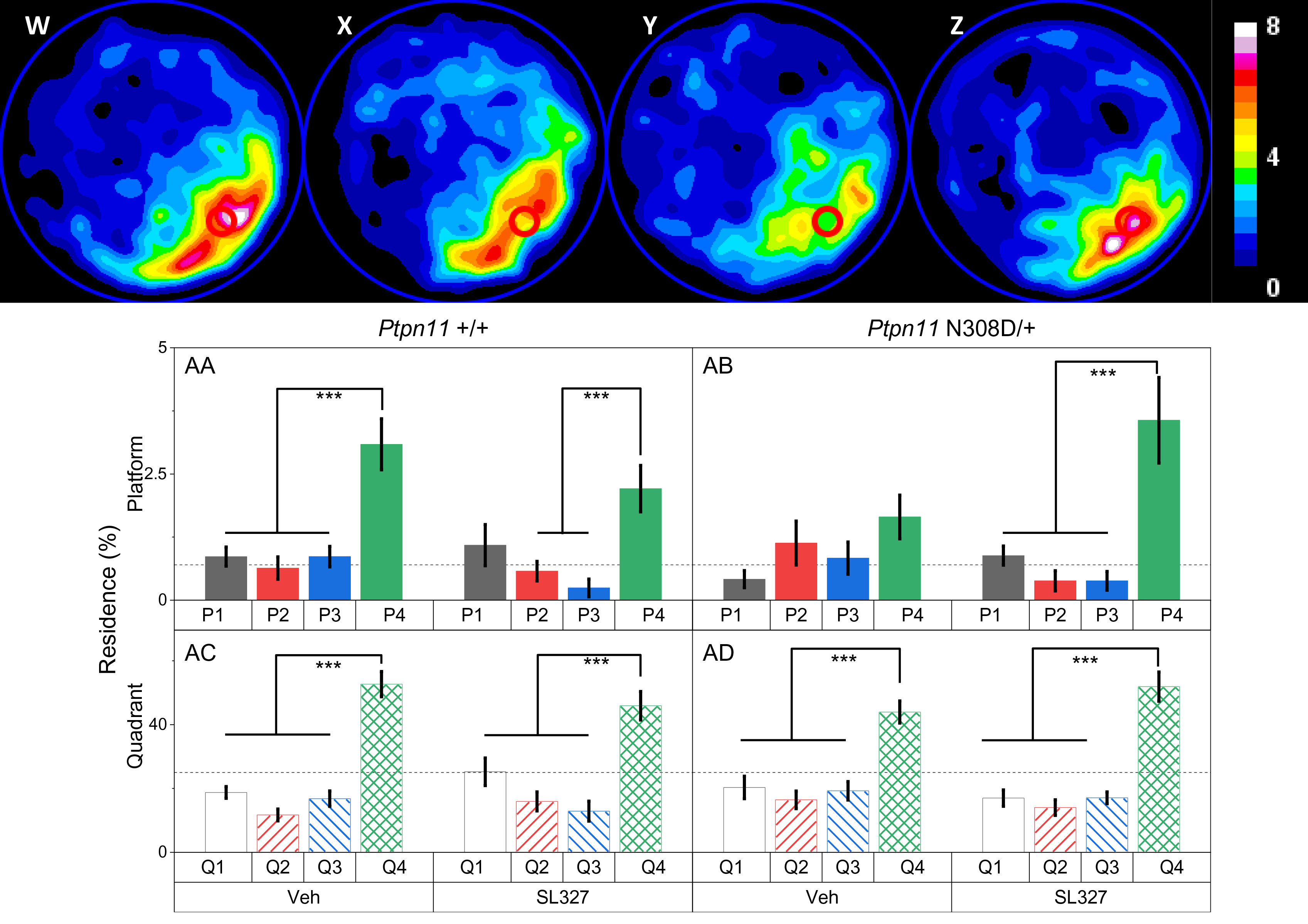
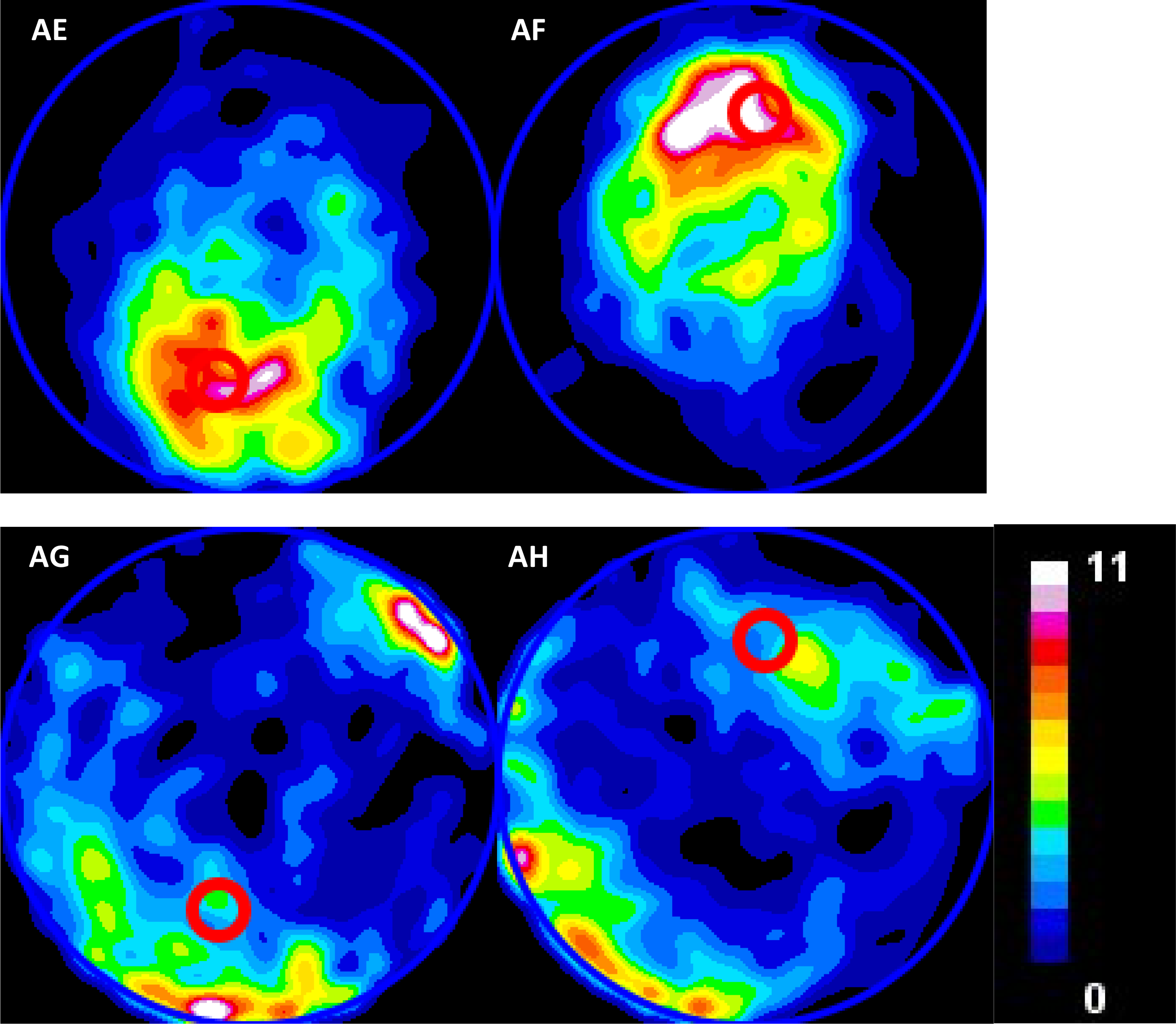
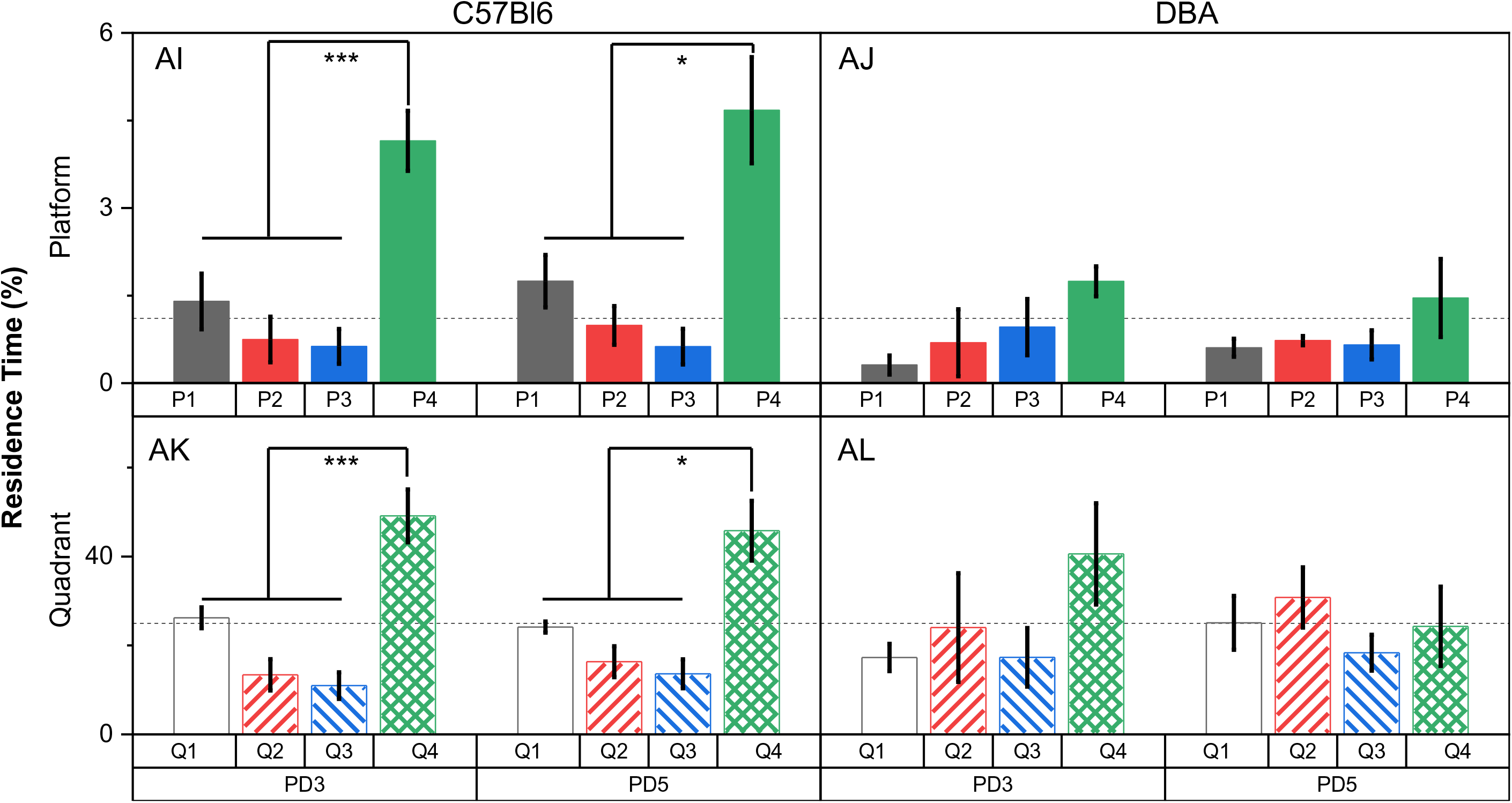
Residence time. The bar graphs represent the residence time measured either as a function of quadrants or as function of platform zones. The data is compared using one-way ANOVA and the means that are significantly different are indicated in the figure with asterisk. The number of asterisks corresponds to degree of significance as described in the main figures. The detailed ANOVA results are presented as S. Table 17 - 24. (A-J) BALB/cJ and SWR/J A-F Heat maps, G-J Plots, (A-J) show the occupancy heat map of BALB/cJ and SWR/J mice on PD3, PD7 and PD10., (K-) *Ptpn11* +/+ and *Ptpn11* D61G/+ K-R Heat maps, S-V, (W-) *Ptpn11* +/+ and *Ptpn11* N308D/+ W-Z Heat maps, AA-AB, (AC-AJ) C57Bl/6J and DBA/2J AC-AF Heat maps, AG-AJ

**Supplementary Figure S5:**
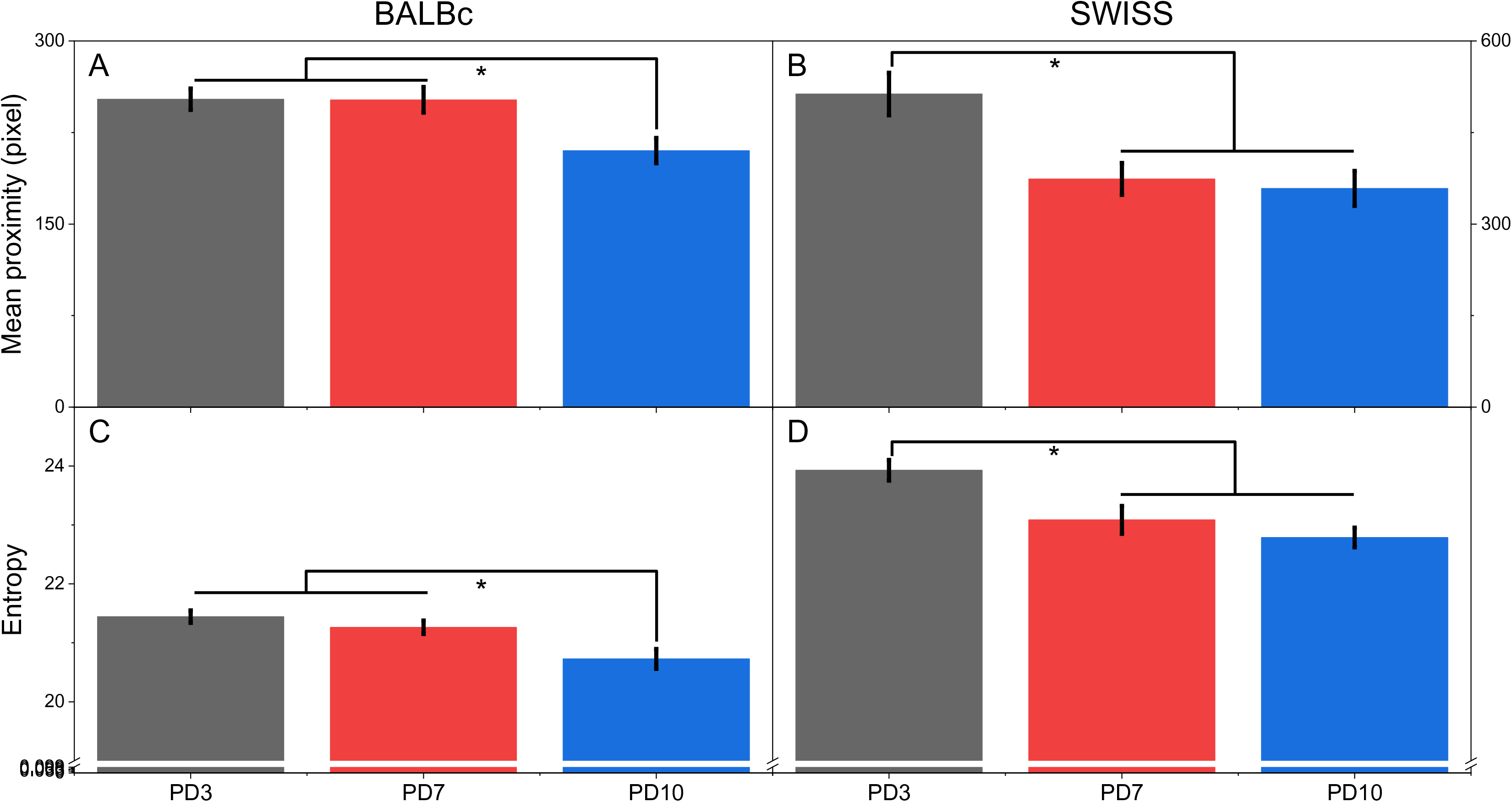
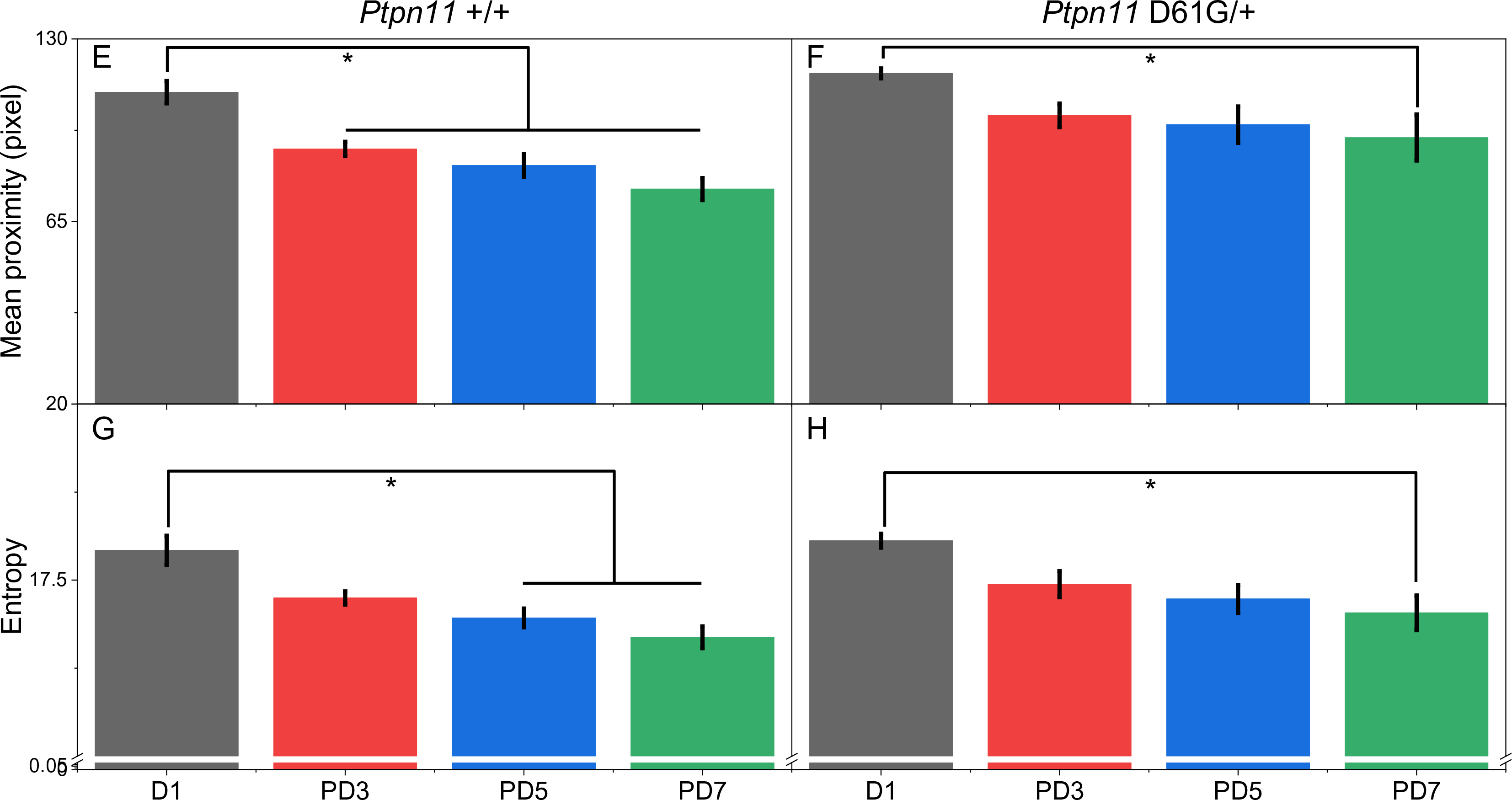
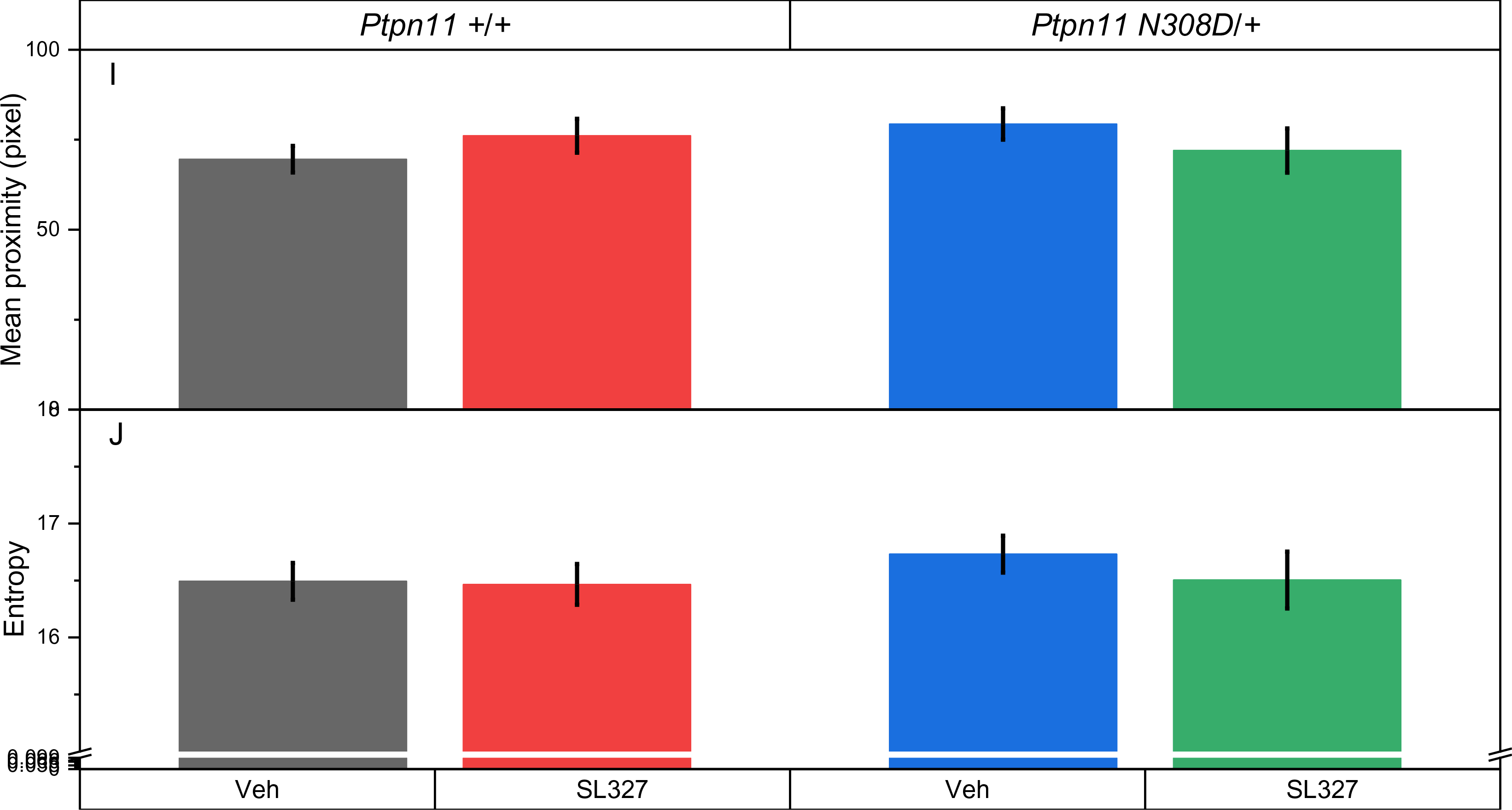
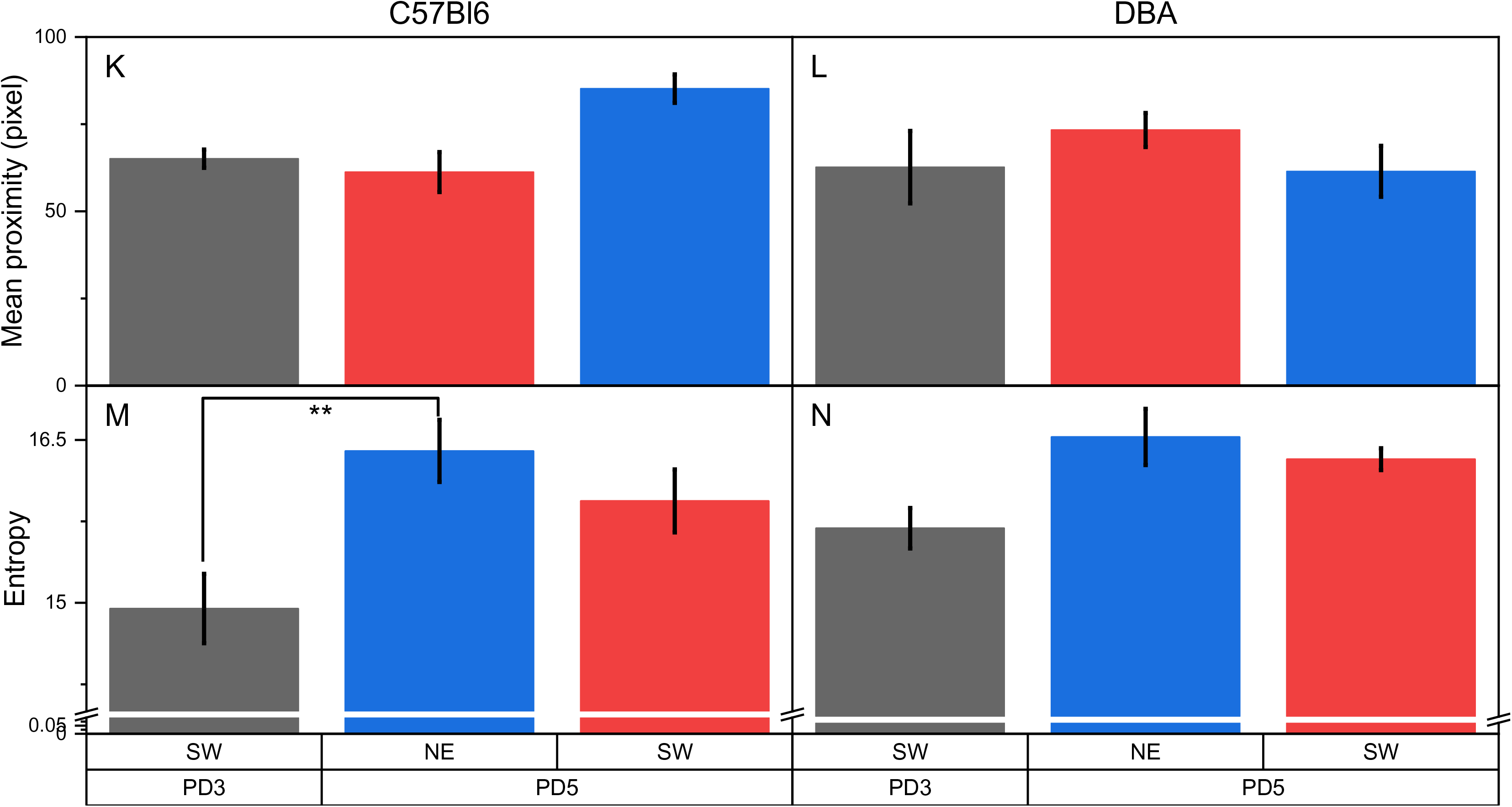
**Proximity measure and entropy measures calculated for datasets analysed in this study.** (A-D) BALB/cJ and SWR/J The bar graphs(X) represent the mean proximity, or the entropy measured for different strains of mice. The data is compared using one-way ANOVA and the means that are significantly different are indicated in the figure with asterisk. The number of asterisks corresponds to degree of significance as described in the main figures. The detailed ANOVA results are presented as S. Tables 25 - 26. (E-H) *Ptpn11* +/+ and *Ptpn11* D61G/+ Two-way ANOVA for mean proximity with genotype as between-subjects factor and training within-subjects factor indicate that differences across genotype: F_1,92_ = 16, p < 0.001 and training: F_3,92_ = 15, p< 0.001 are significant while, genotype × day interaction: F_3,92_ = 0.576, p > 0.05 is not significant. See S. Table 27 for pair-wise comparison. Two-way ANOVA for entropy with genotype as between-subjects factor and training as within-subjects factor indicate that differences across genotype: F_1,92_ = 3.83, p > 0.05, day: F_3,92_ = 16, p< 0.001, genotype × day interaction: F_3,92_ = 0.14, p > 0.05. See S. Table 28 for pair-wise comparison. (I-J) Mean proximity and entropy measures for *Ptpn11* +/+ and *Ptpn11* N308D/+ group of mice. Two-way ANOVA with genotype as between-subjects factor indicated that mean proximity measure is not statistically different between *Ptpn11* +/+ and *Ptpn11* N308D/+ strains (genotype: F_1,38_ = 0.354, p > 0.05), nor across the within-subjects factor treatment between saline (Veh) and SL327 (drug) groups (treatment: F_1,38_ = 0.00757, p > 0.05). The ANOVA Main effect for interaction between genotype and treatment is also not (genotype × treatment interaction: F_1,38_ = 2.13, p > 0.05, S. Table 29). Similarly, the two-way ANOVA with genotype as between-subjects factor and treatment as within-subjects factor, show that the entropy measure was not statistically different between *Ptpn11* +/+ and *Ptpn11* N308D/+ as well as saline and SL327 (genotype: F_1,38_ = 0.53, p > 0.05, treatment: F_1,38_ = 0.45, p > 0.05). The interaction between genotype and treatment is also not statistically significant (genotype × treatment interaction: F_1,38_ = 0.277, p > 0.05. See S. Table 30). Thus, mean proximity and entropy measures are unable to detect the subtle memory deficit, and its rescue by SL327, in *Ptpn11* N308D/+ mutant mice. (K-N) C57Bl/6J and DBA/2J Two-way ANOVA for mean proximity with strain as between-subjects factor and peak as within-subjects factor shows that no significant difference between any factors with no interaction (strain F_1,24_ = 3.61, p > 0.05, peak F_2,24_ = 0.749, p > 0.05, interaction: F_2,24_ = 3.27, p > 0.05). See S. Table 31. Two-way ANOVA for entropy with strain as between-subjects factor and peak as within-subjects factor indicate that the means are different across the, strains (F_2,24_ = 4.441, p < 0.05) but are different for the platform locations as indicated by the peaks (F_2,24_ = 10.97, p < 0.001) with no significant interaction: F_2,24_ = 5.59, p > 0.05. See S. Table 32 for pair-wise comparison.

**Supplementary Figure 6:**
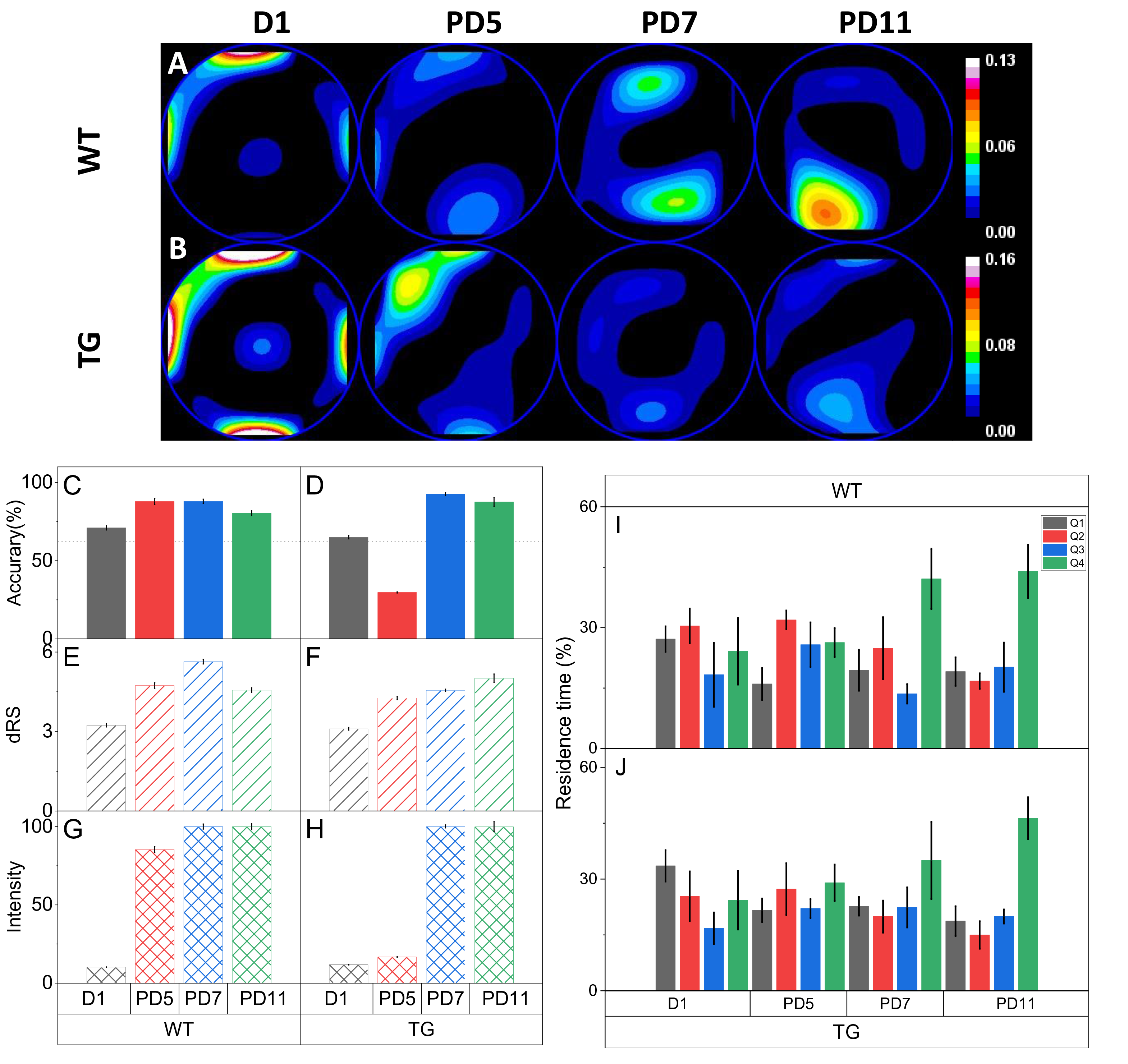
**Divergence is highly sensitive and can detect changes in memory using smaller number of animals compared to conventional residence time-based measures.** (A) shows the convergence heat maps for WT male mice (n = 5) on D1, PD5, PD7 and PD11. The surface on all three probe days (PD5, PD7 and PD11) shows the presence of a distinct convergence peak close to the platform. We infer that WT male mice learn the platform location by PD5. (B) shows the convergence heat maps for D61G male mice (n = 4) on D1, PD5, PD7 and PD11. On PD5, the surface shows a peak away from the centre indicating that the swim patterns are different from D1. However, the field does not convergence to the platform location, indicating the absence of spatial memory specific to the platform. On PD7 and PD11, the surface shows a convergence peak adjacent to the platform location indicating additional training improved the retention of spatial memory for the platform. (C) Accuracy of search centre to the platform location reaches an asymptote maximum on PD5 in WT male mice. A reduction in accuracy seen on PD11 could be due to extinction (during rest days between PD7 and PD11). WT male mice: α = 88 ± 1.9 (PD5), α = 88 ± 1.4 (PD7), α = 80 ± 1.6 (PD11) (D) Accuracy of search centre to the platform location improves as a function of training in TG male mice. From D1 to PD5, the accuracy of search changes from at chance to lower than chance, indicating that the mice have shifted their behaviour from swimming close to the pool walls and are directing their search elsewhere. On PD7, the accuracy of the search centre is comparable to that of its wildtype male littermates. The accuracy of search is maintained on PD11 as well. TG male mice: α = 30 ± 0.4 (PD5), α = 93 ± 0.9 (PD7), α = 88 ± 2.8 (PD11). (E) and (F) shows the uncertainty of search (Horizontal bars). (G) and (H) shows the relative search intensity (Hatched bars). (I) And (J) show the residence time for probe day 5 (PD5), 7 (PD7) and 11 (PD11) with N = 4 (WT), 5 (TG) mice. Q1: Left adjacent, Q2: Opposite, Q3: Right adjacent, Q4: Training/target quadrant.

## Supplementary Tables

**S. Table 1:**
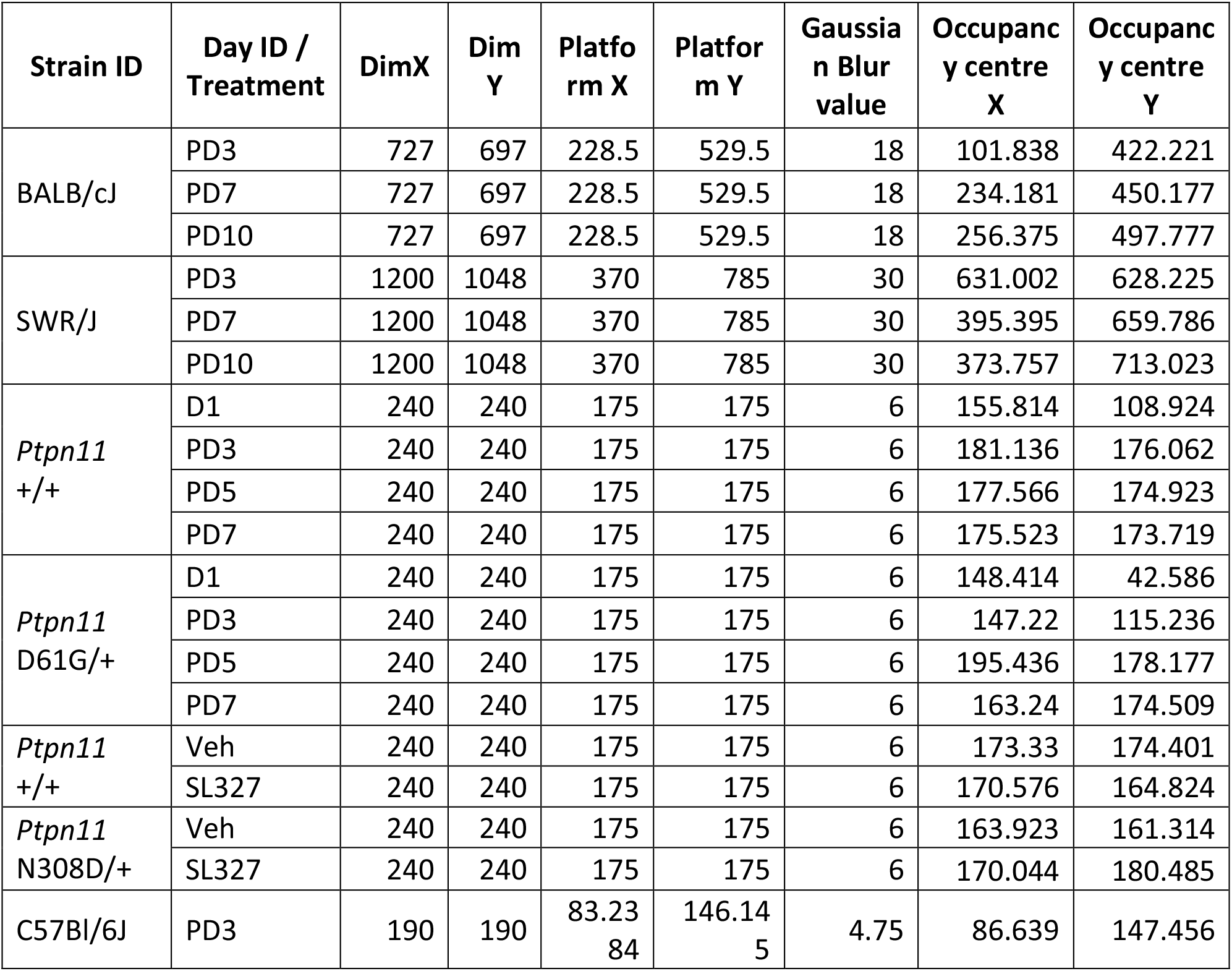
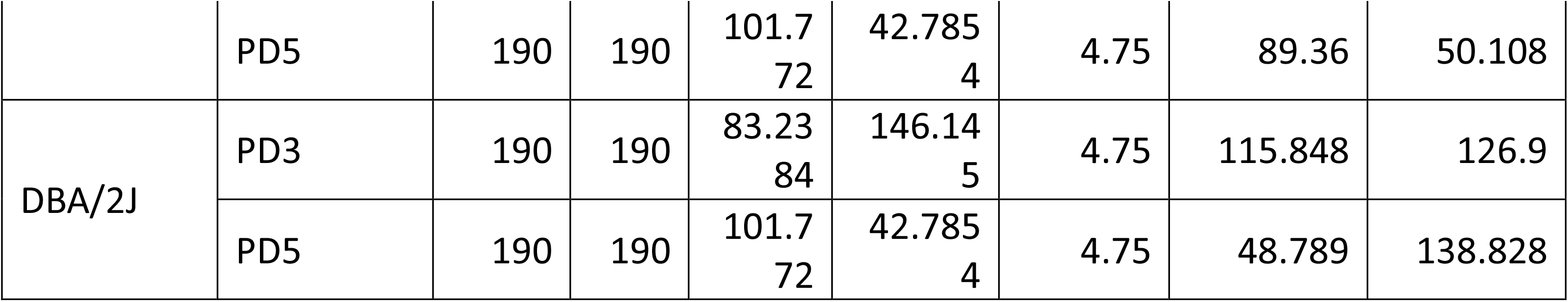
Details utilized for data analysis.

**S. Table 2:** Two-way ANOVA of accuracy measure in BALB/cJ and SWR/J mice.

|  | DF | Sum of Squares | Mean Square | F Value | P Value |
| --- | --- | --- | --- | --- | --- |
| **Genotype** | 1 | 9882.223 | 9882.223 | 466.9975 | 1.22E-27 |
| **Day** | 2 | 3475.977 | 1737.989 | 82.13095 | 8.07E-17 |
| **Interaction** | 2 | 239.3672 | 119.6836 | 5.65581 | 0.00599 |
| **Model** | 5 | 13854.89 | 2770.978 | 130.9462 | 3.29E-28 |
| **Error** | 52 | 1100.382 | 21.16119 |  |  |
| **Corrected Total** | 57 | 14955.27 |  |  |  |

*Bonferroni test:* *Genotype:*
| Genotype | MeanDiff | SEM | t Value | Prob | Alpha | Sig | LCL | UCL |
| --- | --- | --- | --- | --- | --- | --- | --- | --- |
| **SWR/J BALB/CJ** | -26.5274 | 1.20877 | -21.9457 | 5.91E-28 | 0.05 | 1 | -28.953 | -24.1018 |

*Day:*
| Day | MeanDiff | SEM | t Value | Prob | Alpha | Sig | LCL | UCL |
| --- | --- | --- | --- | --- | --- | --- | --- | --- |
| **PD7 PD3** | 10.16935 | 1.4737 | 6.90053 | 2.14E-08 | 0.05 | 1 | 6.52355 | 13.81514 |
| **PD10 PD3** | 19.4354 | 1.4737 | 13.18812 | 9.69E-18 | 0.05 | 1 | 15.78961 | 23.0812 |
| **PD10 PD7** | 9.26606 | 1.49248 | 6.2085 | 2.70E-07 | 0.05 | 1 | 5.57382 | 12.9583 |

*Interactions:*
| Genotype | Day | Genotype | Day | MeanDiff | SEM | t Value | Prob | Alpha | Sig | LCL | UCL |
| --- | --- | --- | --- | --- | --- | --- | --- | --- | --- | --- | --- |
| BALB/CJ | PD7 | BALB/CJ | PD3 | 9.38235 | 2.05724 | 4.56065 | 4.70E-04 | 0.05 | 1 | 3.05249 | 15.71221 |
| BALB/CJ | PD10 | BALB/CJ | PD3 | 14.55459 | 2.05724 | 7.07481 | 5.64E-08 | 0.05 | 1 | 8.22473 | 20.88445 |
| BALB/CJ | PD10 | BALB/CJ | PD7 | 5.17224 | 2.05724 | 2.51416 | 0.22598 | 0.05 | 0 | -1.15762 | 11.5021 |
| SWR/J | PD3 | BALB/CJ | PD3 | -29.0509 | 2.05724 | -14.1213 | 2.97E-18 | 0.05 | 1 | -35.3808 | -22.721 |
| SWR/J | PD3 | BALB/CJ | PD7 | -38.4332 | 2.05724 | -18.6819 | 1.52E-23 | 0.05 | 1 | -44.7631 | -32.1034 |
| SWR/J | PD3 | BALB/CJ | PD10 | -43.6055 | 2.05724 | -21.1961 | 4.52E-26 | 0.05 | 1 | -49.9353 | -37.2756 |
| SWR/J | PD7 | BALB/CJ | PD3 | -19.621 | 2.11361 | -9.28317 | 1.91E-11 | 0.05 | 1 | -26.1244 | -13.1177 |
| SWR/J | PD7 | BALB/CJ | PD7 | -29.0034 | 2.11361 | -13.7222 | 9.67E-18 | 0.05 | 1 | -35.5067 | -22.5001 |
| SWR/J | PD7 | BALB/CJ | PD10 | -34.1756 | 2.11361 | -16.1693 | 9.37E-21 | 0.05 | 1 | -40.6789 | -27.6723 |
| SWR/J | PD7 | SWR/J | PD3 | 9.42985 | 2.11361 | 4.46148 | 6.58E-04 | 0.05 | 1 | 2.92654 | 15.93316 |
| SWR/J | PD10 | BALB/CJ | PD3 | -5.8063 | 2.11361 | -2.7471 | 1.24E-01 | 0.05 | 0 | -12.3096 | 0.69701 |
| SWR/J | PD10 | BALB/CJ | PD7 | -15.1887 | 2.11361 | -7.18611 | 3.75E-08 | 0.05 | 1 | -21.692 | -8.68534 |
| SWR/J | PD10 | BALB/CJ | PD10 | -20.3609 | 2.11361 | -9.63321 | 5.58E-12 | 0.05 | 1 | -26.8642 | -13.8576 |
| SWR/J | PD10 | SWR/J | PD3 | 23.24459 | 2.11361 | 10.99756 | 5.28E-14 | 0.05 | 1 | 16.74128 | 29.7479 |
| SWR/J | PD10 | SWR/J | PD7 | 13.81474 | 2.16852 | 6.37058 | 7.47E-07 | 0.05 | 1 | 7.14248 | 20.487 |

**S. Table 3:** Two-way ANOVA of uncertainty measure in BALB/cJ and SWR/J mice.

|  | DF | Sum of Squares | Mean Square | F Value | P Value |
| --- | --- | --- | --- | --- | --- |
| **Genotype** | 1 | 52.28865 | 52.28865 | 217.2078 | 3.31E-20 |
| **Day** | 2 | 281.9184 | 140.9592 | 585.5467 | 2.20E-36 |
| **Interaction** | 2 | 242.6841 | 121.3421 | 504.0567 | 9.06E-35 |
| **Model** | 5 | 568.7714 | 113.7543 | 472.537 | 4.76E-42 |
| **Error** | 52 | 12.51801 | 0.24073 |  |  |
| **Corrected Total** | 57 | 581.2894 |  |  |  |

*Bonferroni test:* *Genotype:*
| Genotype | MeanDiff | SEM | t Value | Prob | Alpha | Sig | LCL | UCL |
| --- | --- | --- | --- | --- | --- | --- | --- | --- |
| SWR/J BALB/CJ | -2.03313 | 0.12893 | -15.7698 | 1.85E-21 | 0.05 | 1 | -2.29184 | -1.77442 |

*Day:*
| Day | MeanDiff | SEM | t Value | Prob | Alpha | Sig | LCL | UCL |
| --- | --- | --- | --- | --- | --- | --- | --- | --- |
| PD7 PD3 | 5.27921 | 0.15718 | 33.5863 | 1.88E-36 | 0.05 | 1 | 4.89035 | 5.66806 |
| PD10 PD3 | 2.63699 | 0.15718 | 16.77651 | 3.73E-22 | 0.05 | 1 | 2.24813 | 3.02584 |
| PD10 PD7 | -2.64222 | 0.15919 | -16.5983 | 5.96E-22 | 0.05 | 1 | -3.03603 | -2.24841 |

*Interactions:*
| Genotype | Day | Genotype | Day | MeanDiff | SEM | t Value | Prob | Alpha | Sig | LCL | UCL |
| --- | --- | --- | --- | --- | --- | --- | --- | --- | --- | --- | --- |
| BALB/CJ | PD7 | BALB/CJ | PD3 | 1.43146 | 0.21942 | 6.52376 | 4.26E-07 | 0.05 | 1 | 0.75633 | 2.10659 |
| BALB/CJ | PD10 | BALB/CJ | PD3 | 3.24078 | 0.21942 | 14.7696 | 4.54E-19 | 0.05 | 1 | 2.56565 | 3.91591 |
| BALB/CJ | PD10 | BALB/CJ | PD7 | 1.80932 | 0.21942 | 8.24583 | 7.82E-10 | 0.05 | 1 | 1.13419 | 2.48445 |
| SWR/J | PD3 | BALB/CJ | PD3 | -4.03467 | 0.21942 | -18.3877 | 3.11E-23 | 0.05 | 1 | -4.7098 | -3.35954 |
| SWR/J | PD3 | BALB/CJ | PD7 | -5.46613 | 0.21942 | -24.9115 | 2.14E-29 | 0.05 | 1 | -6.14126 | -4.791 |
| SWR/J | PD3 | BALB/CJ | PD10 | -7.27545 | 0.21942 | -33.1573 | 1.79E-35 | 0.05 | 1 | -7.95058 | -6.60032 |
| SWR/J | PD7 | BALB/CJ | PD3 | 5.29566 | 0.22544 | 23.49084 | 3.54E-28 | 0.05 | 1 | 4.60203 | 5.98929 |
| SWR/J | PD7 | BALB/CJ | PD7 | 3.8642 | 0.22544 | 17.14108 | 7.21E-22 | 0.05 | 1 | 3.17057 | 4.55783 |
| SWR/J | PD7 | BALB/CJ | PD10 | 2.05488 | 0.22544 | 9.11517 | 3.46E-11 | 0.05 | 1 | 1.36125 | 2.74851 |
| SWR/J | PD7 | SWR/J | PD3 | 9.33033 | 0.22544 | 41.3881 | 2.66E-40 | 0.05 | 1 | 8.6367 | 10.02396 |
| SWR/J | PD10 | BALB/CJ | PD3 | -2.29271 | 0.22544 | -10.1702 | 8.68E-13 | 0.05 | 1 | -2.98634 | -1.59908 |
| SWR/J | PD10 | BALB/CJ | PD7 | -3.72417 | 0.22544 | -16.5199 | 3.67E-21 | 0.05 | 1 | -4.4178 | -3.03054 |
| SWR/J | PD10 | BALB/CJ | PD10 | -5.53349 | 0.22544 | -24.5458 | 4.34E-29 | 0.05 | 1 | -6.22712 | -4.83986 |
| SWR/J | PD10 | SWR/J | PD3 | 1.74196 | 0.22544 | 7.7271 | 5.17E-09 | 0.05 | 1 | 1.04833 | 2.43559 |
| SWR/J | PD10 | SWR/J | PD7 | -7.58837 | 0.23129 | -32.8087 | 3.02E-35 | 0.05 | 1 | -8.30002 | -6.87672 |

**S. Table 4:** Two-way ANOVA of relative intensity measure in BALB/cJ and SWR/J mice.

|  | DF | Sum of Squares | Mean Square | F Value | P Value |
| --- | --- | --- | --- | --- | --- |
| **Genotype** | 1 | 3796.007 | 3796.007 | 487.5288 | 4.43E-28 |
| **Day** | 2 | 9462.693 | 4731.347 | 607.6563 | 8.73E-37 |
| **Interaction** | 2 | 13007.73 | 6503.866 | 835.3045 | 2.99E-40 |
| **Model** | 5 | 26118.52 | 5223.705 | 670.8909 | 6.25E-46 |
| **Error** | 52 | 404.8835 | 7.78622 |  |  |
| **Corrected Total** | 57 | 26523.41 |  |  |  |

*Bonferroni test:* *Genotype:*
| Genotype | MeanDiff | SEM | t Value | Prob | Alpha | Sig | LCL | UCL |
| --- | --- | --- | --- | --- | --- | --- | --- | --- |
| **SWR/J BALB/CJ** | -16.8857 | 0.73323 | -23.0294 | 6.06E-29 | 0.05 | 1 | -18.357 | -15.4144 |

*Day:*
| Day | MeanDiff | SEM | t Value | Prob | Alpha | Sig | LCL | UCL |
| --- | --- | --- | --- | --- | --- | --- | --- | --- |
| **PD7 PD3** | -14.9947 | 0.89393 | -16.7739 | 3.75E-22 | 0.05 | 1 | -17.2062 | -12.7832 |
| **PD10 PD3** | 15.75263 | 0.89393 | 17.62176 | 4.21E-23 | 0.05 | 1 | 13.54114 | 17.96412 |
| **PD10 PD7** | 30.74737 | 0.90532 | 33.96303 | 1.08E-36 | 0.05 | 1 | 28.5077 | 32.98703 |

Interaction:
| Genotype | Day | Genotype | Day | MeanDiff | SEM | t Value | Prob | Alpha | Sig | LCL | UCL |
| --- | --- | --- | --- | --- | --- | --- | --- | --- | --- | --- | --- |
| BALB/CJ | PD7 | BALB/CJ | PD3 | -35.6 | 1.2479 | -28.528 | 2.97E-32 | 0.05 | 1 | -39.4396 | -31.7604 |
| BALB/CJ | PD10 | BALB/CJ | PD3 | -20.2 | 1.2479 | -16.1873 | 8.93E-21 | 0.05 | 1 | -24.0396 | -16.3604 |
| BALB/CJ | PD10 | BALB/CJ | PD7 | 15.4 | 1.2479 | 12.34077 | 6.74E-16 | 0.05 | 1 | 11.56039 | 19.23961 |
| SWR/J | PD3 | BALB/CJ | PD3 | -54 | 1.2479 | -43.2728 | 2.80E-41 | 0.05 | 1 | -57.8396 | -50.1604 |
| SWR/J | PD3 | BALB/CJ | PD7 | -18.4 | 1.2479 | -14.7448 | 4.87E-19 | 0.05 | 1 | -22.2396 | -14.5604 |
| SWR/J | PD3 | BALB/CJ | PD10 | -33.8 | 1.2479 | -27.0856 | 3.74E-31 | 0.05 | 1 | -37.6396 | -29.9604 |
| SWR/J | PD7 | BALB/CJ | PD3 | -49.1 | 1.28209 | -38.2968 | 1.33E-38 | 0.05 | 1 | -53.0448 | -45.1552 |
| SWR/J | PD7 | BALB/CJ | PD7 | -13.5 | 1.28209 | -10.5297 | 2.55E-13 | 0.05 | 1 | -17.4448 | -9.55517 |
| SWR/J | PD7 | BALB/CJ | PD10 | -28.9 | 1.28209 | -22.5413 | 2.51E-27 | 0.05 | 1 | -32.8448 | -24.9552 |
| SWR/J | PD7 | SWR/J | PD3 | 4.9 | 1.28209 | 3.82188 | 5.33E-03 | 0.05 | 1 | 0.95517 | 8.84483 |
| SWR/J | PD10 | BALB/CJ | PD3 | -1.3 | 1.28209 | -1.01397 | 1.00E+00 | 0.05 | 0 | -5.24483 | 2.64483 |
| SWR/J | PD10 | BALB/CJ | PD7 | 34.3 | 1.28209 | 26.75317 | 6.81E-31 | 0.05 | 1 | 30.35517 | 38.24483 |
| SWR/J | PD10 | BALB/CJ | PD10 | 18.9 | 1.28209 | 14.74154 | 4.92E-19 | 0.05 | 1 | 14.95517 | 22.84483 |
| SWR/J | PD10 | SWR/J | PD3 | 52.7 | 1.28209 | 41.10472 | 3.76E-40 | 0.05 | 1 | 48.75517 | 56.64483 |
| SWR/J | PD10 | SWR/J | PD7 | 47.8 | 1.3154 | 36.33882 | 1.85E-37 | 0.05 | 1 | 43.75269 | 51.84731 |

**S. Table 5:** One-way ANOVA of absolute intensity measure in BALB/cJ and SWR/J mice.

|  | DF | Sum of Squares | Mean Square | F Value | Prob>F |
| --- | --- | --- | --- | --- | --- |
| **Model** | 2 | 2.21E-08 | 1.11E-08 | 4.88E+02 | 6.47E-22 |
| **Error** | 27 | 6.12E-10 | 2.27E-11 |  |  |
| **Total** | 29 | 2.27E-08 |  |  |  |

*Bonferroni test:*
|  | MeanDiff | SEM | t Value | Prob | Alpha | Sig | LCL | UCL |
| --- | --- | --- | --- | --- | --- | --- | --- | --- |
| **PD7 PD3** | -2.17E-05 | 2.13E-06 | -1.02E+01 | 2.85E-10 | 0.05 | 1 | -2.71E-05 | -1.63E-05 |
| **PD10 PD3** | 4.36E-05 | 2.13E-06 | 20.46922 | 1.70E-17 | 0.05 | 1 | 3.82E-05 | 4.90E-05 |
| **PD10 PD7** | 6.53E-05 | 2.13E-06 | 30.65688 | 4.85E-22 | 0.05 | 1 | 5.99E-05 | 7.07E-05 |

|  | DF | Sum of Squares | Mean Square | F Value | Prob>F |
| --- | --- | --- | --- | --- | --- |
| **Model** | 2 | 9.69E-10 | 4.85E-10 | 2.87E+02 | 5.76E-18 |
| **Error** | 25 | 4.22E-11 | 1.69E-12 |  |  |
| **Total** | 27 | 1.01E-09 |  |  |  |

*Bonferroni test:*
|  | MeanDiff | SEM | t Value | Prob | Alpha | Sig | LCL | UCL |
| --- | --- | --- | --- | --- | --- | --- | --- | --- |
| **PD7 PD3** | 1.43E-05 | 5.97E-07 | 2.39E+01 | 2.80E-18 | 0.05 | 1 | 1.28E-05 | 1.58E-05 |
| **PD10 PD3** | 6.50E-06 | 5.97E-07 | 10.88316 | 1.70E-10 | 0.05 | 1 | 4.97E-06 | 8.03E-06 |
| **PD10 PD7** | -7.80E-06 | 6.13E-07 | -12.7291 | 6.02E-12 | 0.05 | 1 | -9.37E-06 | -6.23E-06 |

**S. Table 6:**
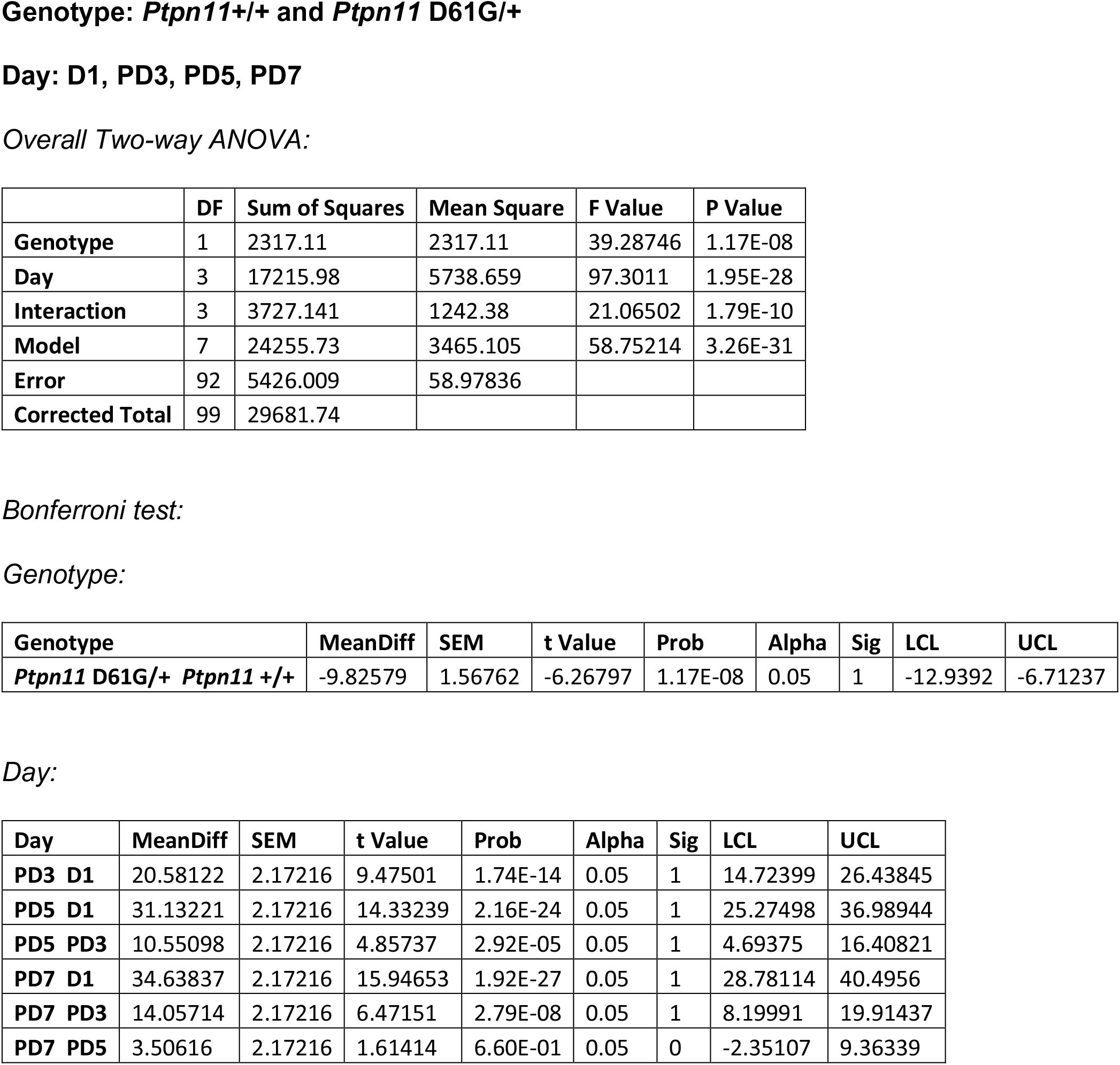
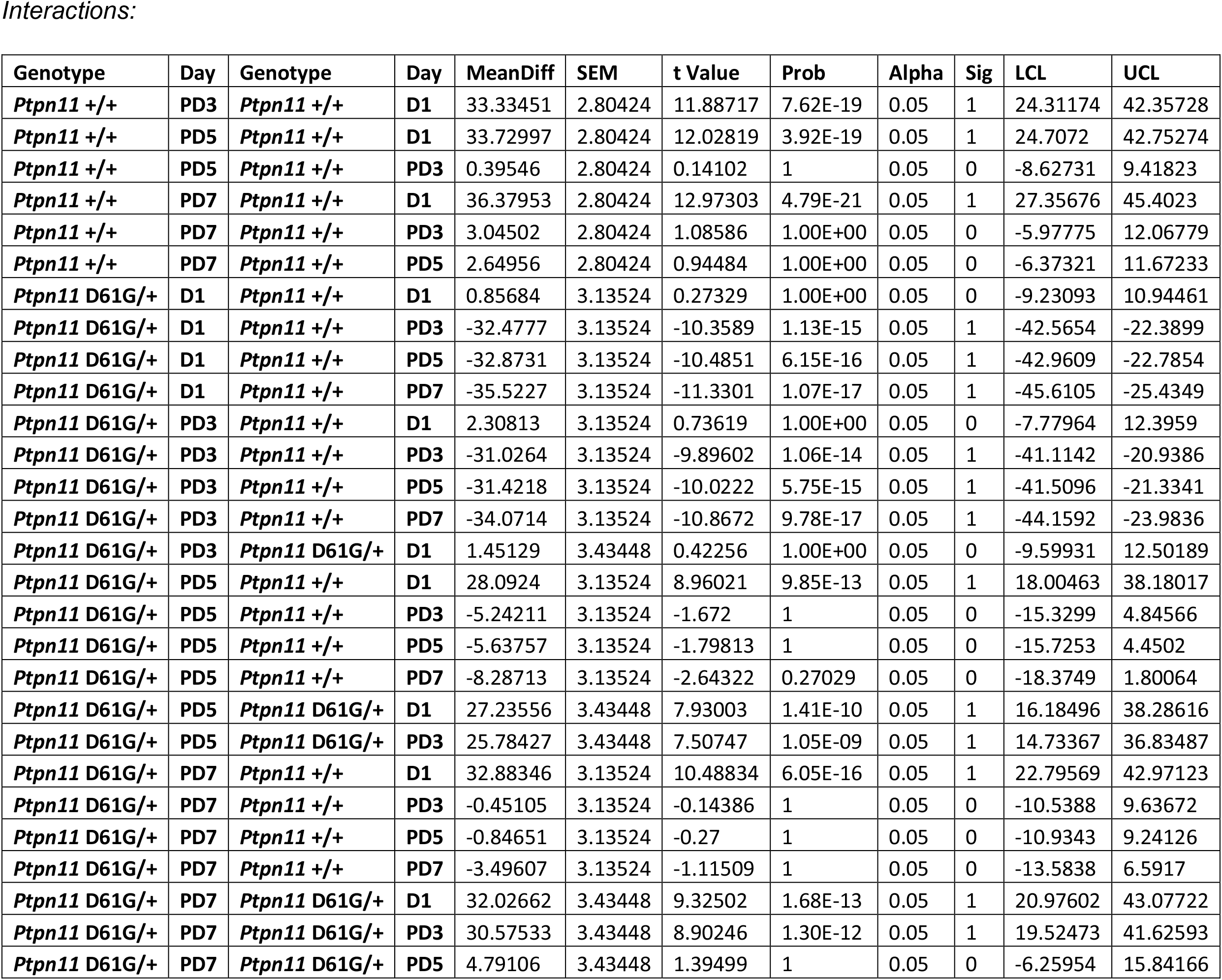
Two-way ANOVA of accuracy measure in *Ptpn11* +/+ and *Ptpn11* D61G/+ mice.

**S. Table 7:** Two-way ANOVA of uncertainty measure in *Ptpn11* +/+ and *Ptpn11* D61G/+ mice.

|  | DF | Sum of Squares | Mean Square | F Value | P Value |
| --- | --- | --- | --- | --- | --- |
| **Genotype** | 1 | 5.93001 | 5.93001 | 1.77958 | 1.85E-01 |
| **Day** | 3 | 14.66631 | 4.88877 | 1.46711 | 2.29E-01 |
| **Interaction** | 3 | 9.95995 | 3.31998 | 0.99632 | 3.98E-01 |
| **Model** | 7 | 30.22296 | 4.31757 | 1.29569 | 2.61E-01 |
| **Error** | 92 | 306.5666 | 3.33225 |  |  |
| **Corrected Total** | 99 | 336.7896 |  |  |  |

**Genotype: *Ptpn11*+/+ and *Ptpn11* D61G/+**

**S. Table 8:** Two-way ANOVA of relative search intensity measure in *Ptpn11* +/+ and *Ptpn11* D61G/+ mice.

|  | DF | Sum of Squares | Mean Square | F Value | P Value |
| --- | --- | --- | --- | --- | --- |
| **Genotype** | 1 | 33390.96 | 33390.96 | 1.35E+07 | 1.85E-239 |
| **Day** | 3 | 100643.6 | 33547.88 | 1.36E+07 | 1.56E-259 |
| **Interaction** | 3 | 33594.36 | 11198.12 | 4.53E+06 | 1.30E-237 |
| **Model** | 7 | 172275.8 | 24610.82 | 9.95E+06 | 1.75E-267 |
| **Error** | 92 | 0.22765 | 2.47E-03 |  |  |
| **Corrected Total** | 99 | 172276 |  |  |  |

*Bonferroni test:* *Genotype:*
| Genotype | MeanDiff | SEM | t Value | Prob | Alpha | Sig | LCL | UCL |
| --- | --- | --- | --- | --- | --- | --- | --- | --- |
| Ptpn11 D61G/+ Ptpn11 +/+ | -37.3 | 0.01015 | -3673.43 | 1.55E-239 | 0.05 | 1 | -37.3202 | -37.2798 |

Day:
| Day | MeanDiff | SEM | t Value | Prob | Alpha | Sig | LCL | UCL |
| --- | --- | --- | --- | --- | --- | --- | --- | --- |
| PD3 D1 | 44.2 | 0.01407 | 3141.486 | 1.68E-232 | 0.05 | 1 | 44.16206 | 44.23794 |
| PD5 D1 | 64.24 | 0.01407 | 4565.816 | 1.85E-247 | 0.05 | 1 | 64.20206 | 64.27794 |
| PD5 PD3 | 20.04 | 0.01407 | 1424.33 | 7.28E-201 | 0.05 | 1 | 20.00206 | 20.07794 |
| PD7 D1 | 88.44 | 0.01407 | 6285.815 | 2.97E-260 | 0.05 | 1 | 88.40206 | 88.47794 |
| PD7 PD3 | 44.24 | 0.01407 | 3144.329 | 1.55E-232 | 0.05 | 1 | 44.20206 | 44.27794 |
| PD7 PD5 | 24.2 | 0.01407 | 1719.999 | 2.09E-208 | 0.05 | 1 | 24.16206 | 24.23794 |

Interactions:
| Genotype | Day | Genotype | Day | MeanDiff | SEM | t Value | Prob | Alpha | Sig | LCL | UCL |
| --- | --- | --- | --- | --- | --- | --- | --- | --- | --- | --- | --- |
| Ptpn11 +/+ | PD3 | Ptpn11 +/+ | D1 | 78 | 0.01816 | 4294.208 | 2.45E-244 | 0.05 | 1 | 77.94156 | 78.05844 |
| Ptpn11 +/+ | PD5 | Ptpn11 +/+ | D1 | 87.6 | 0.01816 | 4822.726 | 5.56E-249 | 0.05 | 1 | 87.54156 | 87.65844 |
| Ptpn11 +/+ | PD5 | Ptpn11 +/+ | PD3 | 9.6 | 0.01816 | 528.5179 | 1.44E-160 | 0.05 | 1 | 9.54156 | 9.65844 |
| Ptpn11 +/+ | PD7 | Ptpn11 +/+ | D1 | 87.6 | 0.01816 | 4822.726 | 5.56E-249 | 0.05 | 1 | 87.54156 | 87.65844 |
| Ptpn11 +/+ | PD7 | Ptpn11 +/+ | PD3 | 9.6 | 0.01816 | 528.5179 | 1.44E-160 | 0.05 | 1 | 9.54156 | 9.65844 |
| Ptpn11 +/+ | PD7 | Ptpn11 +/+ | PD5 | 2.84E-14 | 0.01816 | 1.56E-12 | 1.00E+00 | 0.05 | 0 | -0.05844 | 0.05844 |
| Ptpn11 D61G/+ | D1 | Ptpn11 +/+ | D1 | -2.1 | 0.02031 | -103.408 | 1.55E-95 | 0.05 | 1 | -2.16534 | -2.03466 |
| Ptpn11 D61G/+ | D1 | Ptpn11 +/+ | PD3 | -80.1 | 0.02031 | -3944.26 | 6.18E-241 | 0.05 | 1 | -80.1653 | -80.0347 |
| Ptpn11 D61G/+ | D1 | Ptpn11 +/+ | PD5 | -89.7 | 0.02031 | -4416.98 | 1.83E-245 | 0.05 | 1 | -89.7653 | -89.6347 |
| Ptpn11 D61G/+ | D1 | Ptpn11 +/+ | PD7 | -89.7 | 0.02031 | -4416.98 | 1.83E-245 | 0.05 | 1 | -89.7653 | -89.6347 |
| Ptpn11 D61G/+ | PD3 | Ptpn11 +/+ | D1 | -8.6 | 0.02031 | -423.479 | 1.03E-151 | 0.05 | 1 | -8.66534 | -8.53466 |
| Ptpn11 D61G/+ | PD3 | Ptpn11 +/+ | PD3 | -86.6 | 0.02031 | -4264.34 | 4.67E-244 | 0.05 | 1 | -86.6653 | -86.5347 |
| Ptpn11 D61G/+ | PD3 | Ptpn11 +/+ | PD5 | -96.2 | 0.02031 | -4737.06 | 2.90E-248 | 0.05 | 1 | -96.2653 | -96.1347 |
| Ptpn11 D61G/+ | PD3 | Ptpn11 +/+ | PD7 | -96.2 | 0.02031 | -4737.06 | 2.90E-248 | 0.05 | 1 | -96.2653 | -96.1347 |
| Ptpn11 D61G/+ | PD3 | Ptpn11 D61G/+ | D1 | -6.5 | 0.02225 | -292.184 | 6.81E-137 | 0.05 | 1 | -6.57158 | -6.42842 |
| Ptpn11 D61G/+ | PD5 | Ptpn11 +/+ | D1 | 27.1 | 0.02031 | 1334.451 | 1.37E-197 | 0.05 | 1 | 27.03466 | 27.16534 |
| Ptpn11 D61G/+ | PD5 | Ptpn11 +/+ | PD3 | -50.9 | 0.02031 | -2506.41 | 8.50E-223 | 0.05 | 1 | -50.9653 | -50.8347 |
| Ptpn11 D61G/+ | PD5 | Ptpn11 +/+ | PD5 | -60.5 | 0.02031 | -2979.13 | 1.04E-229 | 0.05 | 1 | -60.5653 | -60.4347 |
| Ptpn11 D61G/+ | PD5 | Ptpn11 +/+ | PD7 | -60.5 | 0.02031 | -2979.13 | 1.04E-229 | 0.05 | 1 | -60.5653 | -60.4347 |
| Ptpn11 D61G/+ | PD5 | Ptpn11 D61G/+ | D1 | 29.2 | 0.02225 | 1312.58 | 6.28E-197 | 0.05 | 1 | 29.12842 | 29.27158 |
| Ptpn11 D61G/+ | PD5 | Ptpn11 D61G/+ | PD3 | 35.7 | 0.02225 | 1604.764 | 5.78E-205 | 0.05 | 1 | 35.62842 | 35.77158 |
| Ptpn11 D61G/+ | PD7 | Ptpn11 +/+ | D1 | 87.6 | 0.02031 | 4313.577 | 1.62E-244 | 0.05 | 1 | 87.53466 | 87.66534 |
| Ptpn11 D61G/+ | PD7 | Ptpn11 +/+ | PD3 | 9.6 | 0.02031 | 472.7208 | 4.15E-156 | 0.05 | 1 | 9.53466 | 9.66534 |
| Ptpn11 D61G/+ | PD7 | Ptpn11 +/+ | PD5 | 7.11E-14 | 0.02031 | 3.50E-12 | 1 | 0.05 | 0 | -0.06534 | 0.06534 |
| Ptpn11 D61G/+ | PD7 | Ptpn11 +/+ | PD7 | 4.26E-14 | 0.02031 | 2.10E-12 | 1 | 0.05 | 0 | -0.06534 | 0.06534 |
| Ptpn11 D61G/+ | PD7 | Ptpn11 D61G/+ | D1 | 89.7 | 0.02225 | 4032.137 | 8.11E-242 | 0.05 | 1 | 89.62842 | 89.77158 |
| Ptpn11 D61G/+ | PD7 | Ptpn11 D61G/+ | PD3 | 96.2 | 0.02225 | 4324.321 | 1.29E-244 | 0.05 | 1 | 96.12842 | 96.27158 |
| Ptpn11 D61G/+ | PD7 | Ptpn11 D61G/+ | PD5 | 60.5 | 0.02225 | 2719.557 | 4.62E-226 | 0.05 | 1 | 60.42842 | 60.57158 |

**Genotype: *Ptpn11*+/+ and *Ptpn11* D61G/+**

**S. Table 9:**
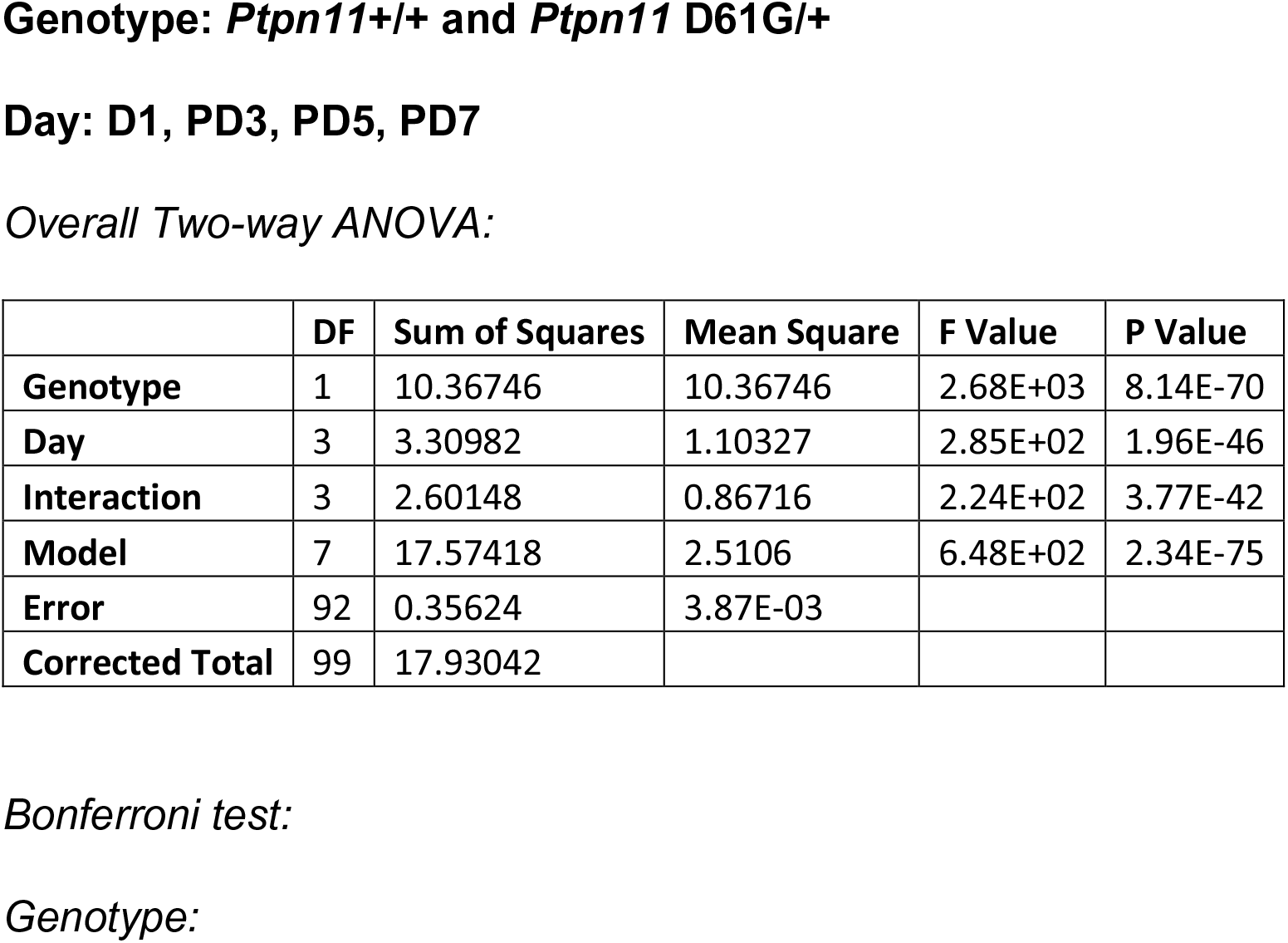
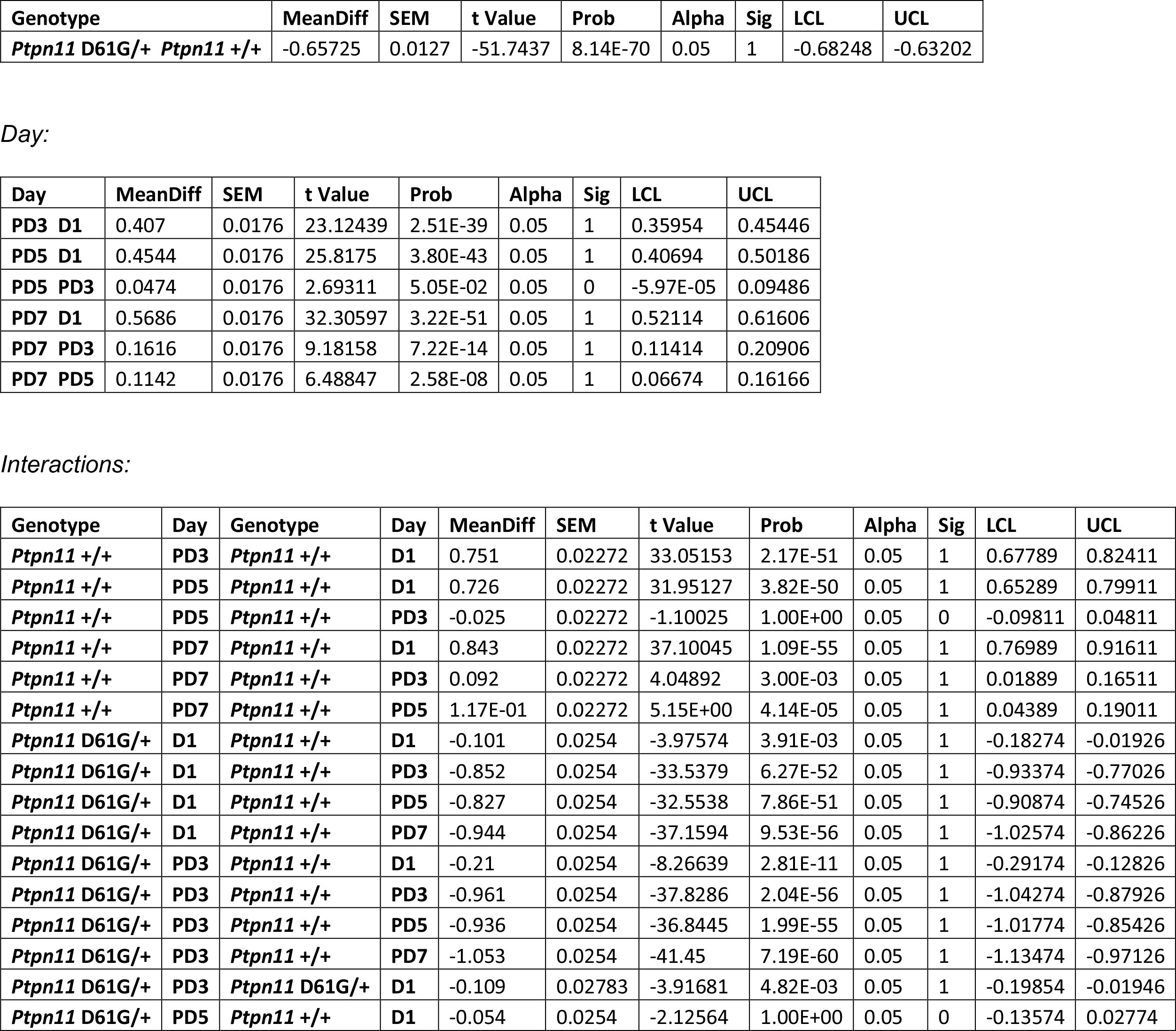
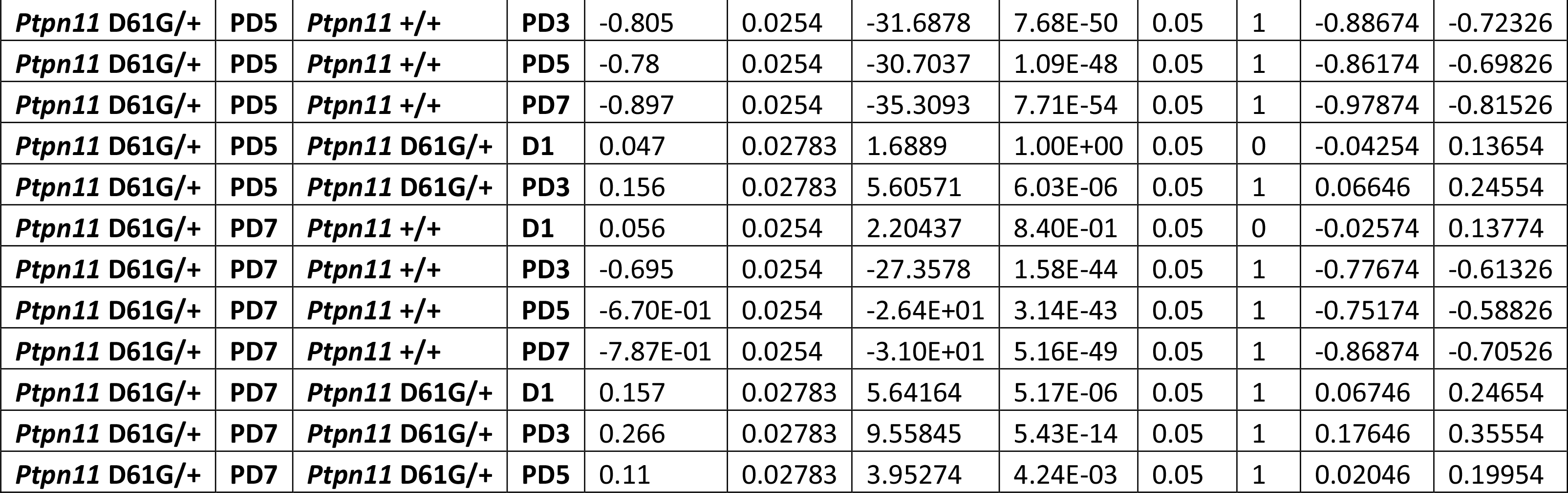
Two-way ANOVA of absolute search intensity measure in *Ptpn11* +/+ and *Ptpn11* D61G/+ mice.

|  | DF | Sum of Squares | Mean Square | F Value | P Value |
| --- | --- | --- | --- | --- | --- |
| **Genotype** | 1 | 10.36746 | 10.36746 | 2.68E+03 | 8.14E-70 |
| **Day** | 3 | 3.30982 | 1.10327 | 2.85E+02 | 1.96E-46 |
| **Interaction** | 3 | 2.60148 | 0.86716 | 2.24E+02 | 3.77E-42 |
| **Model** | 7 | 17.57418 | 2.5106 | 6.48E+02 | 2.34E-75 |
| **Error** | 92 | 0.35624 | 3.87E-03 |  |  |
| **Corrected Total** | 99 | 17.93042 |  |  |  |

*Bonferroni test:* *Genotype:*
| Genotype | MeanDiff | SEM | t Value | Prob | Alpha | Sig | LCL | UCL |
| --- | --- | --- | --- | --- | --- | --- | --- | --- |
| *Ptpn11* D61G/+ *Ptpn11* +/+ | -0.65725 | 0.0127 | -51.7437 | 8.14E-70 | 0.05 | 1 | -0.68248 | -0.63202 |

Day:
| Day | MeanDiff | SEM | t Value | Prob | Alpha | Sig | LCL | UCL |
| --- | --- | --- | --- | --- | --- | --- | --- | --- |
| PD3 D1 | 0.407 | 0.0176 | 23.12439 | 2.51E-39 | 0.05 | 1 | 0.35954 | 0.45446 |
| PD5 D1 | 0.4544 | 0.0176 | 25.8175 | 3.80E-43 | 0.05 | 1 | 0.40694 | 0.50186 |
| PD5 PD3 | 0.0474 | 0.0176 | 2.69311 | 5.05E-02 | 0.05 | 0 | -5.97E-05 | 0.09486 |
| PD7 D1 | 0.5686 | 0.0176 | 32.30597 | 3.22E-51 | 0.05 | 1 | 0.52114 | 0.61606 |
| PD7 PD3 | 0.1616 | 0.0176 | 9.18158 | 7.22E-14 | 0.05 | 1 | 0.11414 | 0.20906 |
| PD7 PD5 | 0.1142 | 0.0176 | 6.48847 | 2.58E-08 | 0.05 | 1 | 0.06674 | 0.16166 |

Interactions:
| Genotype | Day | Genotype | Day | MeanDiff | SEM | t Value | Prob | Alpha | Sig | LCL | UCL |
| --- | --- | --- | --- | --- | --- | --- | --- | --- | --- | --- | --- |
| *Ptpn11* +/+ | PD3 | *Ptpn11* +/+ | D1 | 0.751 | 0.02272 | 33.05153 | 2.17E-51 | 0.05 | 1 | 0.67789 | 0.82411 |
| *Ptpn11* +/+ | PD5 | *Ptpn11* +/+ | D1 | 0.726 | 0.02272 | 31.95127 | 3.82E-50 | 0.05 | 1 | 0.65289 | 0.79911 |
| *Ptpn11* +/+ | PD5 | *Ptpn11* +/+ | PD3 | -0.025 | 0.02272 | -1.10025 | 1.00E+00 | 0.05 | 0 | -0.09811 | 0.04811 |
| *Ptpn11* +/+ | PD7 | *Ptpn11* +/+ | D1 | 0.843 | 0.02272 | 37.10045 | 1.09E-55 | 0.05 | 1 | 0.76989 | 0.91611 |
| *Ptpn11* +/+ | PD7 | *Ptpn11* +/+ | PD3 | 0.092 | 0.02272 | 4.04892 | 3.00E-03 | 0.05 | 1 | 0.01889 | 0.16511 |
| *Ptpn11* +/+ | PD7 | *Ptpn11* +/+ | PD5 | 1.17E-01 | 0.02272 | 5.15E+00 | 4.14E-05 | 0.05 | 1 | 0.04389 | 0.19011 |
| *Ptpn11* D61G/+ | D1 | *Ptpn11* +/+ | D1 | -0.101 | 0.0254 | -3.97574 | 3.91E-03 | 0.05 | 1 | -0.18274 | -0.01926 |
| *Ptpn11* D61G/+ | D1 | *Ptpn11* +/+ | PD3 | -0.852 | 0.0254 | -33.5379 | 6.27E-52 | 0.05 | 1 | -0.93374 | -0.77026 |
| *Ptpn11* D61G/+ | D1 | *Ptpn11* +/+ | PD5 | -0.827 | 0.0254 | -32.5538 | 7.86E-51 | 0.05 | 1 | -0.90874 | -0.74526 |
| *Ptpn11* D61G/+ | D1 | *Ptpn11* +/+ | PD7 | -0.944 | 0.0254 | -37.1594 | 9.53E-56 | 0.05 | 1 | -1.02574 | -0.86226 |
| *Ptpn11* D61G/+ | PD3 | *Ptpn11* +/+ | D1 | -0.21 | 0.0254 | -8.26639 | 2.81E-11 | 0.05 | 1 | -0.29174 | -0.12826 |
| *Ptpn11* D61G/+ | PD3 | *Ptpn11* +/+ | PD3 | -0.961 | 0.0254 | -37.8286 | 2.04E-56 | 0.05 | 1 | -1.04274 | -0.87926 |
| *Ptpn11* D61G/+ | PD3 | *Ptpn11* +/+ | PD5 | -0.936 | 0.0254 | -36.8445 | 1.99E-55 | 0.05 | 1 | -1.01774 | -0.85426 |
| *Ptpn11* D61G/+ | PD3 | *Ptpn11* +/+ | PD7 | -1.053 | 0.0254 | -41.45 | 7.19E-60 | 0.05 | 1 | -1.13474 | -0.97126 |
| *Ptpn11* D61G/+ | PD3 | *Ptpn11* D61G/+ | D1 | -0.109 | 0.02783 | -3.91681 | 4.82E-03 | 0.05 | 1 | -0.19854 | -0.01946 |
| *Ptpn11* D61G/+ | PD5 | *Ptpn11* +/+ | D1 | -0.054 | 0.0254 | -2.12564 | 1.00E+00 | 0.05 | 0 | -0.13574 | 0.02774 |
| *Ptpn11* D61G/+ | PD5 | *Ptpn11* +/+ | PD3 | -0.805 | 0.0254 | -31.6878 | 7.68E-50 | 0.05 | 1 | -0.88674 | -0.72326 |
| *Ptpn11* D61G/+ | PD5 | *Ptpn11* +/+ | PD5 | -0.78 | 0.0254 | -30.7037 | 1.09E-48 | 0.05 | 1 | -0.86174 | -0.69826 |
| *Ptpn11* D61G/+ | PD5 | *Ptpn11* +/+ | PD7 | -0.897 | 0.0254 | -35.3093 | 7.71E-54 | 0.05 | 1 | -0.97874 | -0.81526 |
| *Ptpn11* D61G/+ | PD5 | *Ptpn11* D61G/+ | D1 | 0.047 | 0.02783 | 1.6889 | 1.00E+00 | 0.05 | 0 | -0.04254 | 0.13654 |
| *Ptpn11* D61G/+ | PD5 | *Ptpn11* D61G/+ | PD3 | 0.156 | 0.02783 | 5.60571 | 6.03E-06 | 0.05 | 1 | 0.06646 | 0.24554 |
| *Ptpn11* D61G/+ | PD7 | *Ptpn11* +/+ | D1 | 0.056 | 0.0254 | 2.20437 | 8.40E-01 | 0.05 | 0 | -0.02574 | 0.13774 |
| *Ptpn11* D61G/+ | PD7 | *Ptpn11* +/+ | PD3 | -0.695 | 0.0254 | -27.3578 | 1.58E-44 | 0.05 | 1 | -0.77674 | -0.61326 |
| *Ptpn11* D61G/+ | PD7 | *Ptpn11* +/+ | PD5 | -6.70E-01 | 0.0254 | -2.64E+01 | 3.14E-43 | 0.05 | 1 | -0.75174 | -0.58826 |
| *Ptpn11* D61G/+ | PD7 | *Ptpn11* +/+ | PD7 | -7.87E-01 | 0.0254 | -3.10E+01 | 5.16E-49 | 0.05 | 1 | -0.86874 | -0.70526 |
| *Ptpn11* D61G/+ | PD7 | *Ptpn11* D61G/+ | D1 | 0.157 | 0.02783 | 5.64164 | 5.17E-06 | 0.05 | 1 | 0.06746 | 0.24654 |
| *Ptpn11* D61G/+ | PD7 | *Ptpn11* D61G/+ | PD3 | 0.266 | 0.02783 | 9.55845 | 5.43E-14 | 0.05 | 1 | 0.17646 | 0.35554 |
| *Ptpn11* D61G/+ | PD7 | *Ptpn11* D61G/+ | PD5 | 0.11 | 0.02783 | 3.95274 | 4.24E-03 | 0.05 | 1 | 0.02046 | 0.19954 |

**S. Table 10:**
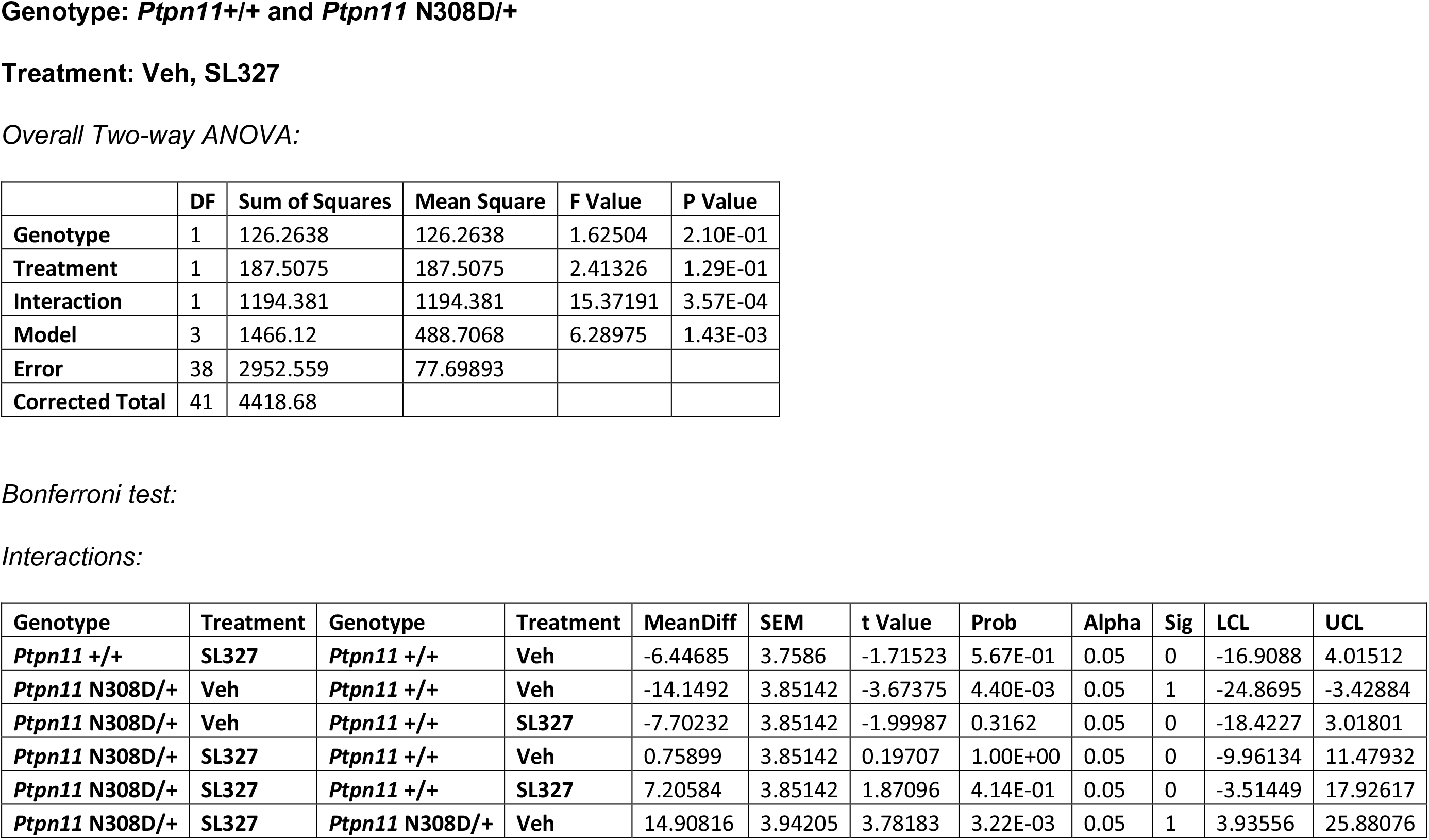
Two-way ANOVA of accuracy measure in *Ptpn11* +/+ and *Ptpn11* N308D/+ mice.

**S. Table 11:** Two-way ANOVA of uncertainty measure in *Ptpn11* +/+ and *Ptpn11* N308D/+ mice.

|  | DF | Sum of Squares | Mean Square | F Value | P Value |
| --- | --- | --- | --- | --- | --- |
| **Genotype** | 1 | 0.85302 | 0.85302 | 3.32122 | 7.63E-02 |
| **Treatment** | 1 | 0.01761 | 0.01761 | 0.06857 | 7.95E-01 |
| **Interaction** | 1 | 14.4231 | 14.4231 | 56.15618 | 5.33E-09 |
| **Model** | 3 | 15.37466 | 5.12489 | 19.95369 | 6.23E-08 |
| **Error** | 38 | 9.75988 | 0.25684 |  |  |
| **Corrected Total** | 41 | 25.13454 |  |  |  |

Bonferroni test: Interactions:
| Genotype | Treatment | Genotype | Treatment | MeanDiff | SEM | t Value | Prob | Alpha | Sig | LCL | UCL |
| --- | --- | --- | --- | --- | --- | --- | --- | --- | --- | --- | --- |
| *Ptpn11* +/+ | SL327 | *Ptpn11* +/+ | Veh | 1.21435 | 0.2161 | 5.61946 | 1.14E-05 | 0.05 | 1 | 0.61285 | 1.81585 |
| *Ptpn11* N308D/+ | Veh | *Ptpn11* +/+ | Veh | 1.4587 | 0.22143 | 6.58752 | 5.36E-07 | 0.05 | 1 | 0.84234 | 2.07506 |
| *Ptpn11* N308D/+ | Veh | *Ptpn11* +/+ | SL327 | 0.24435 | 0.22143 | 1.10349 | 1 | 0.05 | 0 | -0.37201 | 0.86071 |
| *Ptpn11* N308D/+ | SL327 | *Ptpn11* +/+ | Veh | 0.32635 | 0.22143 | 1.4738 | 8.93E-01 | 0.05 | 0 | -0.29001 | 0.94271 |
| *Ptpn11* N308D/+ | SL327 | *Ptpn11* +/+ | SL327 | -0.888 | 0.22143 | -4.01023 | 1.64E-03 | 0.05 | 1 | -1.50436 | -0.27164 |
| *Ptpn11* N308D/+ | SL327 | *Ptpn11* N308D/+ | Veh | -1.13235 | 0.22664 | -4.99615 | 8.06E-05 | 0.05 | 1 | -1.76321 | -0.50149 |

Genotype: *Ptpn11*+/+ and *Ptpn11* N308D/+
Treatment: Veh, SL327
Overall Two-way ANOVA:

**S. Table 12:** Two-way ANOVA of relative search intensity measure in *Ptpn11* +/+ and *Ptpn11* N308D/+ mice.

|  | DF | Sum of Squares | Mean Square | F Value | P Value |
| --- | --- | --- | --- | --- | --- |
| **Genotype** | 1 | 35.85476 | 35.85476 | 0.36005 | 5.52E-01 |
| **Treatment** | 1 | 35.85476 | 35.85476 | 0.36005 | 5.52E-01 |
| Interaction | 1 | 35.85476 | 35.85476 | 0.36005 | 5.52E-01 |
| Model | 3 | 111.1498 | 37.04992 | 0.37205 | 7.74E-01 |
| Error | 38 | 3784.136 | 9.96E+01 |  |  |
| Corrected Total | 41 | 3895.286 |  |  |  |

**S. Table 13:**
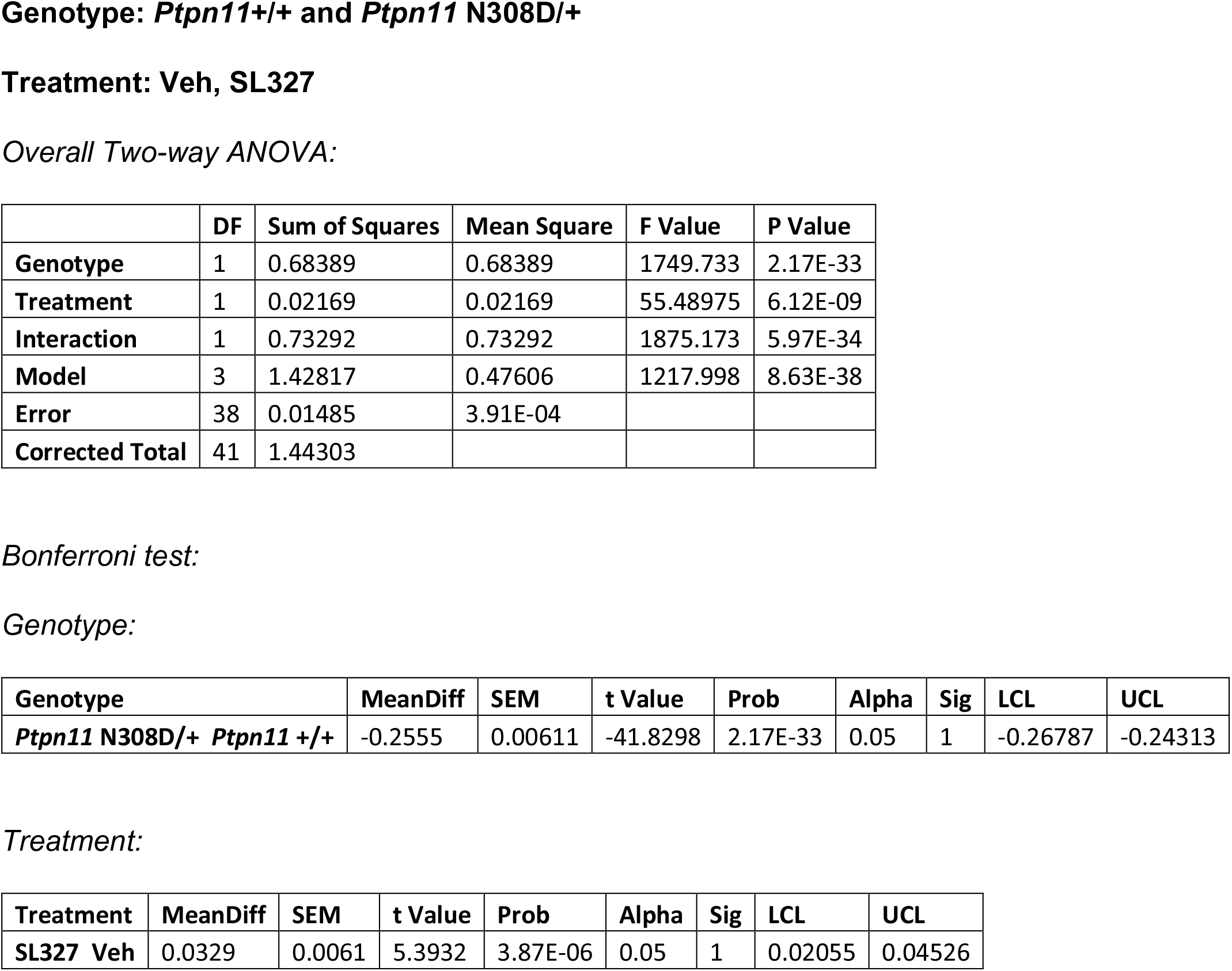
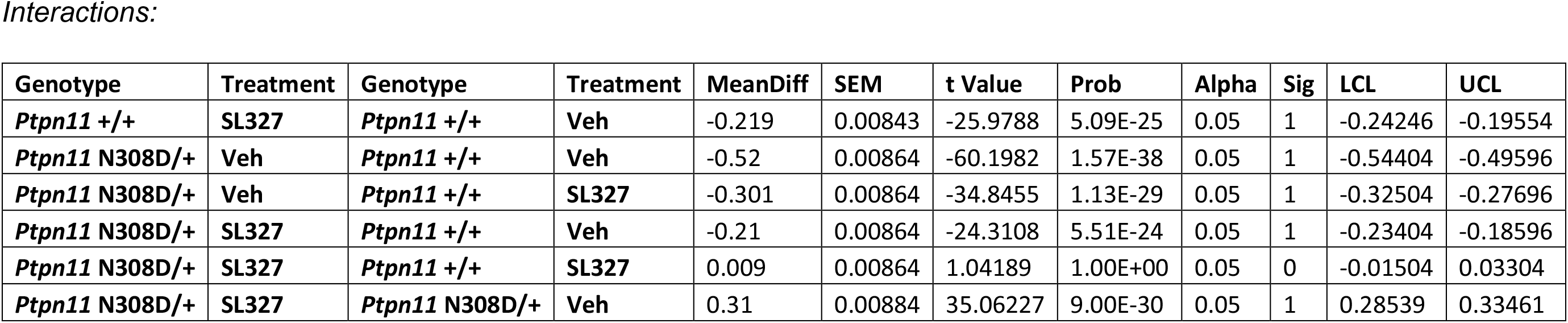
Two-way ANOVA of absolute search intensity measure in *Ptpn11* +/+ and *Ptpn11* N308D/+ mice.

**S. Table 14(a):** One-way ANOVA of accuracy measure in C57Bl/6J and DBA/2J mice.

|  | DF | Sum of Squares | Mean Square | F Value | Prob>F |
| --- | --- | --- | --- | --- | --- |
| Model | 2 | 0.12519 | 0.0626 | 3.43E-03 | 0.99657 |
| Error | 12 | 218.8674 | 18.23895 |  |  |
| Total | 14 | 218.9926 |  |  |  |

|  | DF | Sum of Squares | Mean Square | F Value | Prob>F |
| --- | --- | --- | --- | --- | --- |
| Model | 1 | 48.76434 | 48.76434 | 1.09E+00 | 0.32783 |
| Error | 8 | 359.235 | 44.90437 |  |  |
| Total | 9 | 407.9993 |  |  |  |

**S. Table 14(b):** Two-way ANOVA of accuracy measure in C57Bl/6J and DBA/2J mice.

|  | DF | Sum of Squares | Mean Square | F Value | P Value |
| --- | --- | --- | --- | --- | --- |
| **Genotype** | 1 | 20.46507 | 20.46507 | 0.78459 | 3.89E-01 |
| **Peaks** | 1 | 23.39631 | 23.39631 | 0.89696 | 3.58E-01 |
| **Interaction** | 1 | 25.38838 | 25.38838 | 0.97334 | 3.39E-01 |
| **Model** | 3 | 69.24976 | 23.08325 | 0.88496 | 4.70E-01 |
| **Error** | 16 | 417.342 | 2.61E+01 |  |  |
| **Corrected Total** | 19 | 486.5918 |  |  |  |

**S. Table 15(a):**
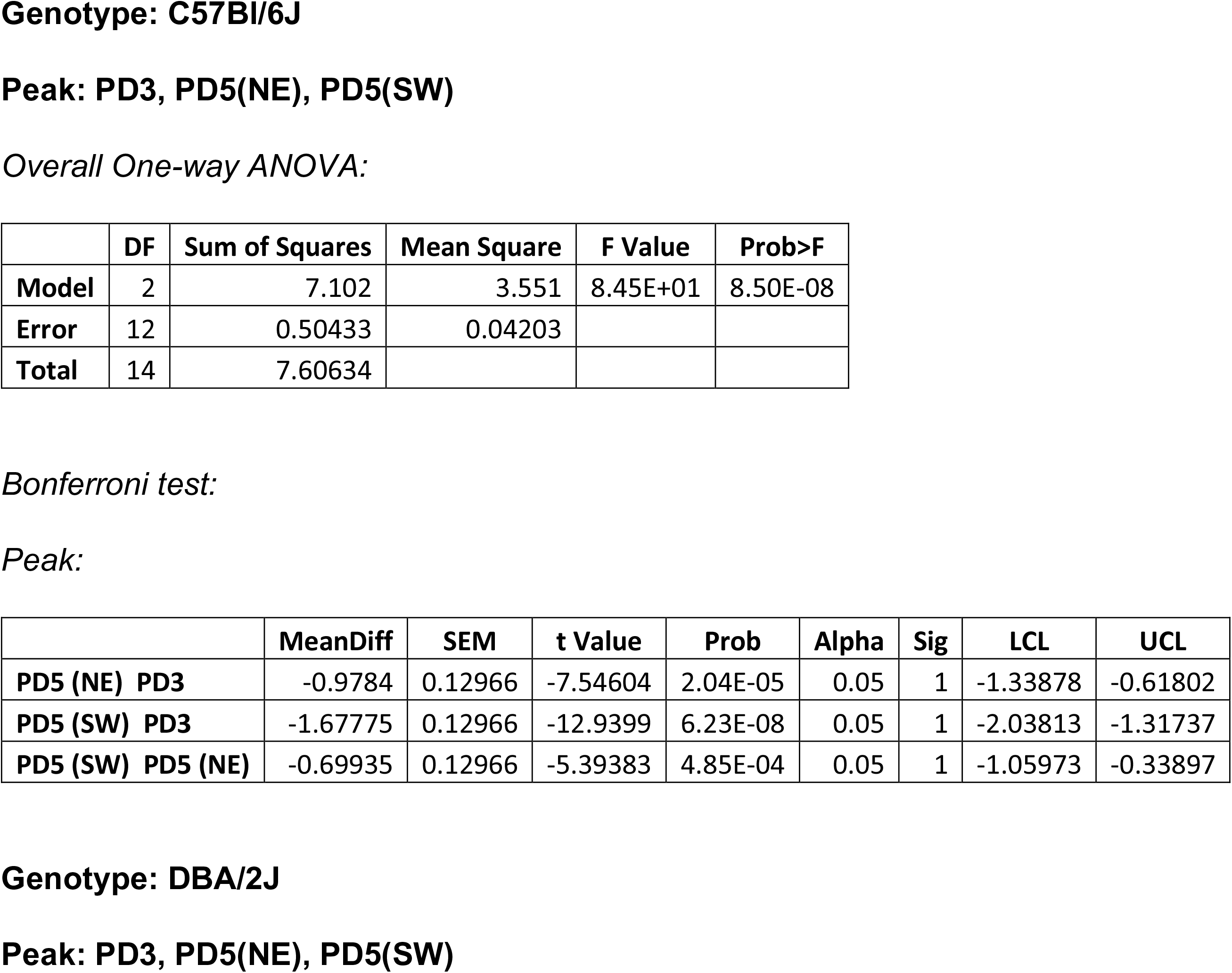
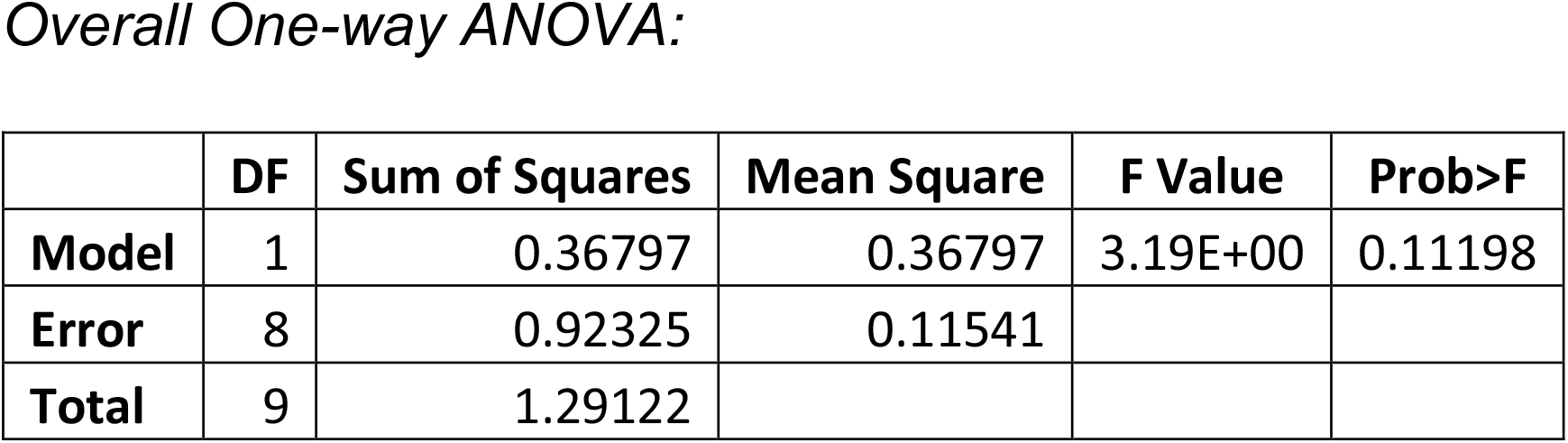
One-way ANOVA of accuracy measure in C57Bl/6J and DBA/2J mice.

|  | DF | Sum of Squares | Mean Square | F Value | Prob>F |
| --- | --- | --- | --- | --- | --- |
| **Model** | 2 | 7.102 | 3.551 | 8.45E+01 | 8.50E-08 |
| **Error** | 12 | 0.50433 | 0.04203 |  |  |
| **Total** | 14 | 7.60634 |  |  |  |

*Bonferroni test:* *Peak:*
|  | MeanDiff | SEM | t Value | Prob | Alpha | Sig | LCL | UCL |
| --- | --- | --- | --- | --- | --- | --- | --- | --- |
| **PD5 (NE) PD3** | -0.9784 | 0.12966 | -7.54604 | 2.04E-05 | 0.05 | 1 | -1.33878 | -0.61802 |
| **PD5 (SW) PD3** | -1.67775 | 0.12966 | -12.9399 | 6.23E-08 | 0.05 | 1 | -2.03813 | -1.31737 |
| **PD5 (SW) PD5 (NE)** | -0.69935 | 0.12966 | -5.39383 | 4.85E-04 | 0.05 | 1 | -1.05973 | -0.33897 |
| **Model** | 1 | 0.36797 | 0.36797 | 3.19E+00 | 0.11198 |
| **Error** | 8 | 0.92325 | 0.11541 |  |  |
| **Total** | 9 | 1.29122 |  |  |  |

**S. Table 15(b):**
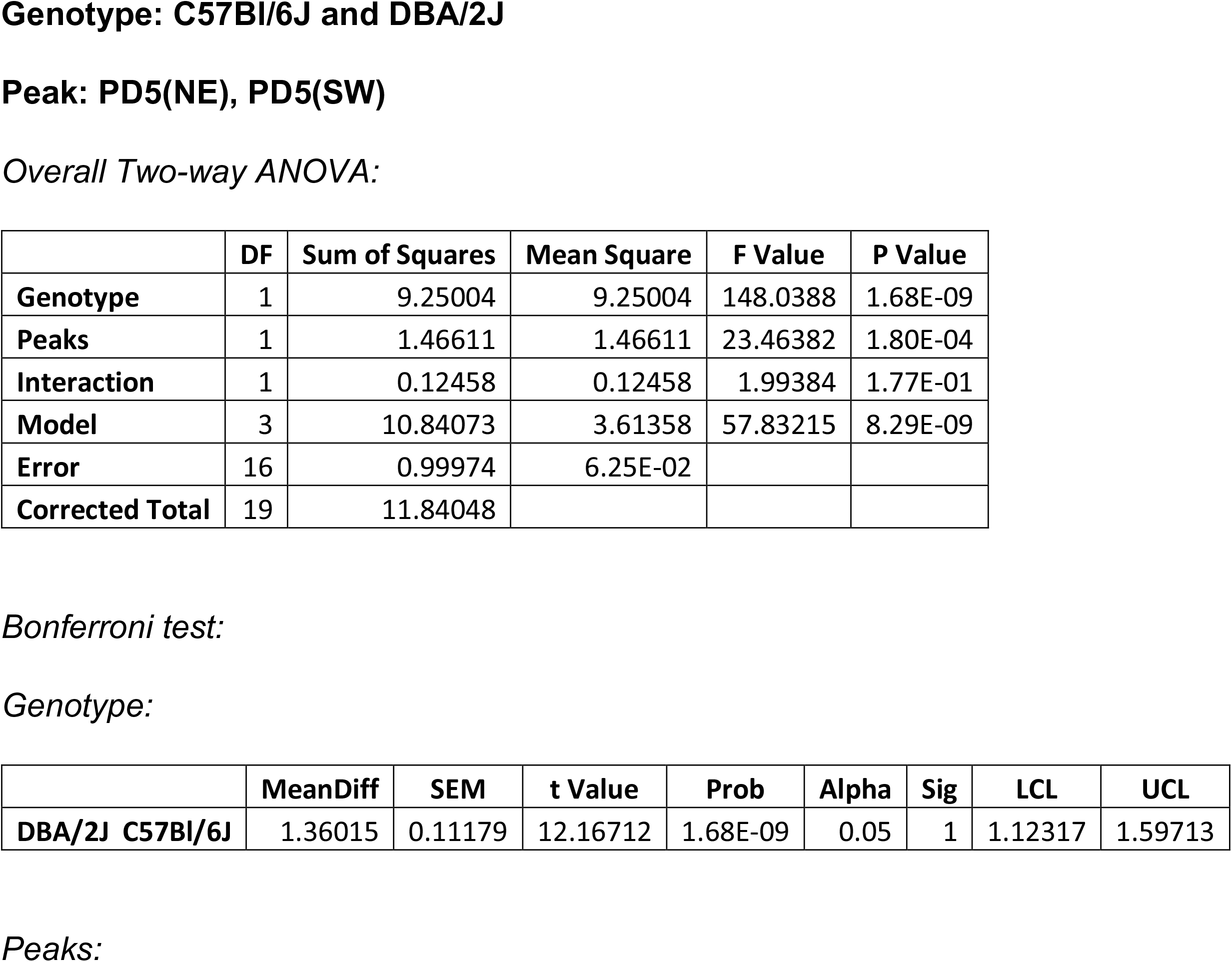
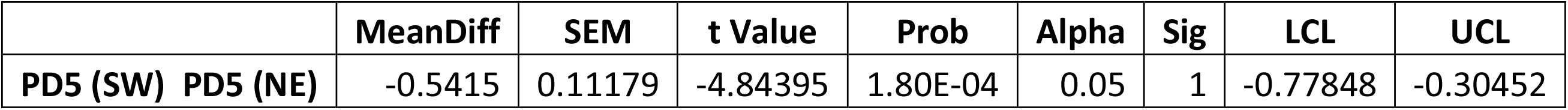
Two-way ANOVA of uncertainty measure in C57Bl/6J and DBA/2J mice.

**S. Table 16(a):** One-way ANOVA of accuracy measure in C57Bl/6J and DBA/2J mice.

|  | DF | Sum of Squares | Mean Square | F Value | Prob>F |
| --- | --- | --- | --- | --- | --- |
| **Model** | 2 | 25433.43 | 12716.72 | 5.92E+02 | 1.02E-12 |
| **Error** | 12 | 257.7107 | 21.47589 |  |  |
| **Total** | 14 | 25691.14 |  |  |  |

*Bonferroni test:* *Peak:*
|  | MeanDiff | SEM | t Value | Prob | Alpha | Sig | LCL | UCL |
| --- | --- | --- | --- | --- | --- | --- | --- | --- |
| PD5 (NE) PD3 | 0.1 | 2.93093 | 0.03412 | 1.00E+00 | 0.05 | 0 | -8.04644 | 8.24644 |
| PD5 (SW) PD3 | -87.3 | 2.93093 | -29.7858 | 3.85E-12 | 0.05 | 1 | -95.4464 | -79.1536 |
| PD5 (SW) PD5 (NE) | -87.4 | 2.93093 | -29.8199 | 3.79E-12 | 0.05 | 1 | -95.5464 | -79.2536 |

|  | DF | Sum of Squares | Mean Square | F Value | Prob>F |
| --- | --- | --- | --- | --- | --- |
| **Model** | 1 | 731.025 | 731.025 | 3.80E+01 | 2.70E-04 |
| **Error** | 8 | 153.8777 | 19.23471 |  |  |
| **Total** | 9 | 884.9027 |  |  |  |

*Bonferroni test:* *Peak:*
|  | MeanDiff | SEM | t Value | Prob | Alpha | Sig | LCL | UCL |
| --- | --- | --- | --- | --- | --- | --- | --- | --- |
| PD5 (SW) PD5 (NE) | -17.1 | 2.77379 | -6.16486 | 2.70E-04 | 0.05 | 1 | -23.4964 | -10.7036 |

**S. Table 16(b):**
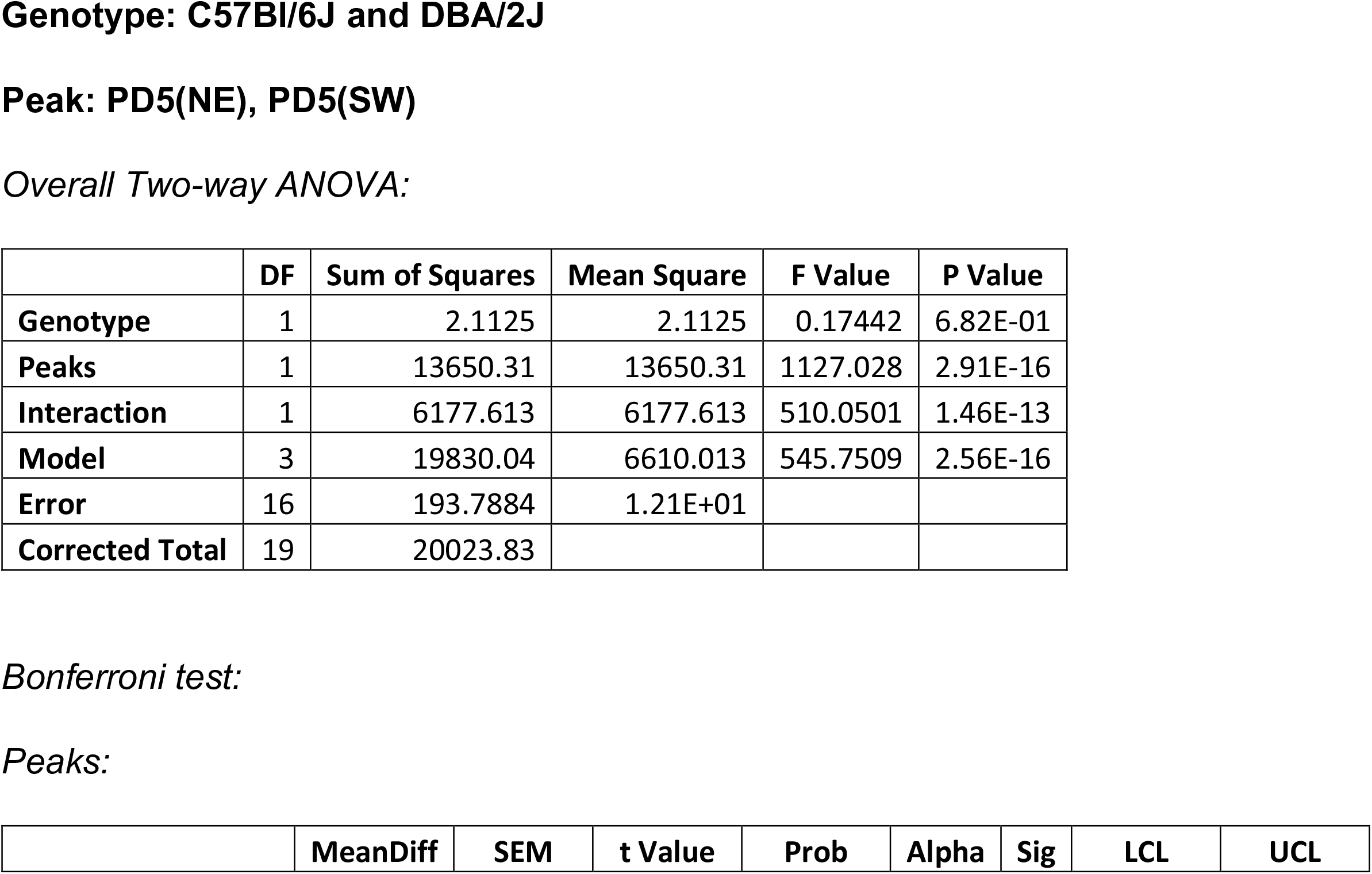
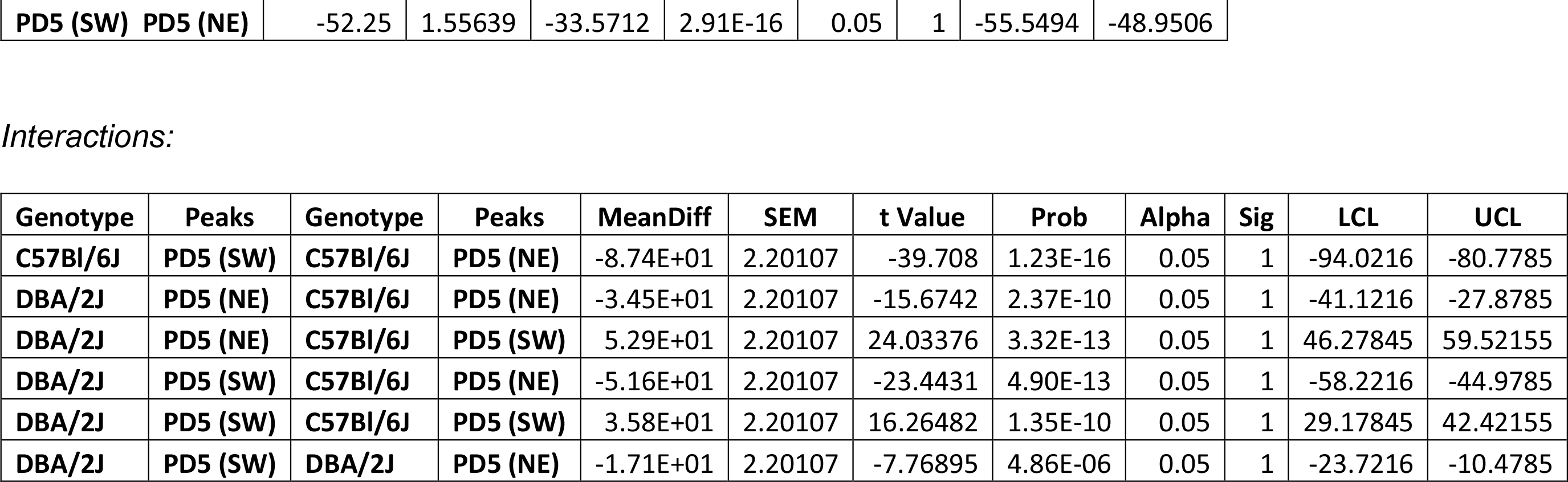
Two-way ANOVA of relative intensity measure in C57Bl/6J and DBA/2J mice.

**S. Table 17:**
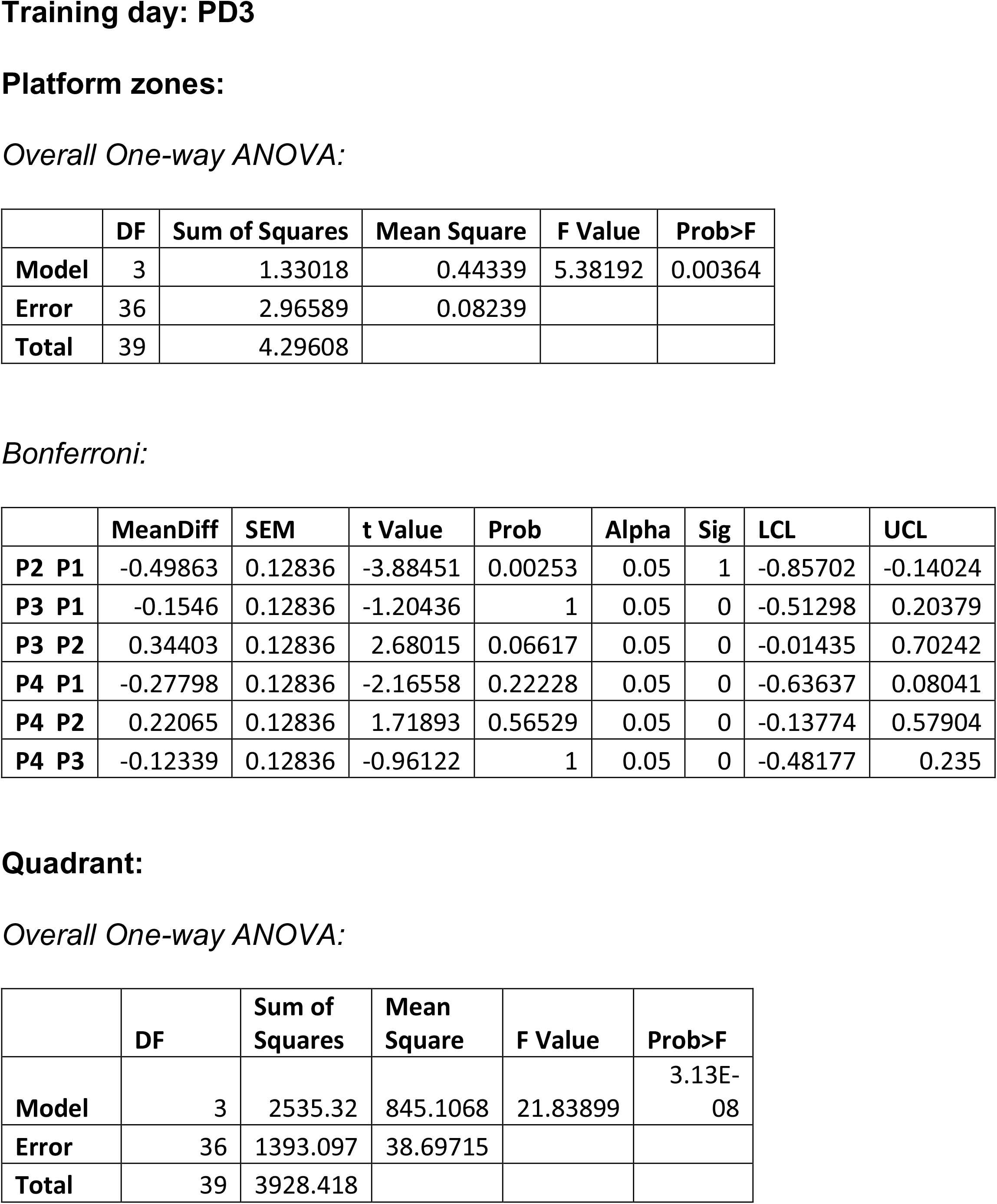
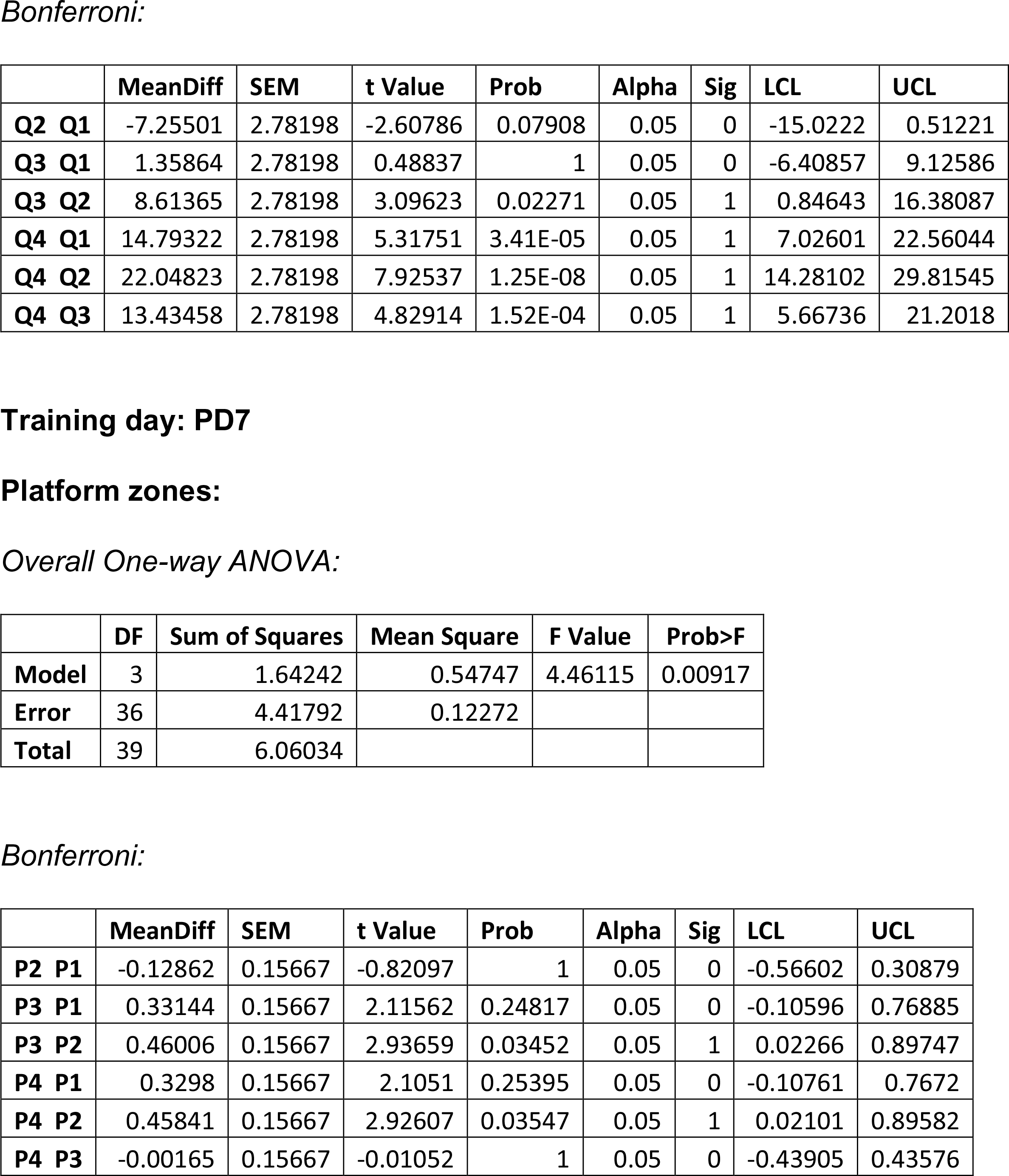
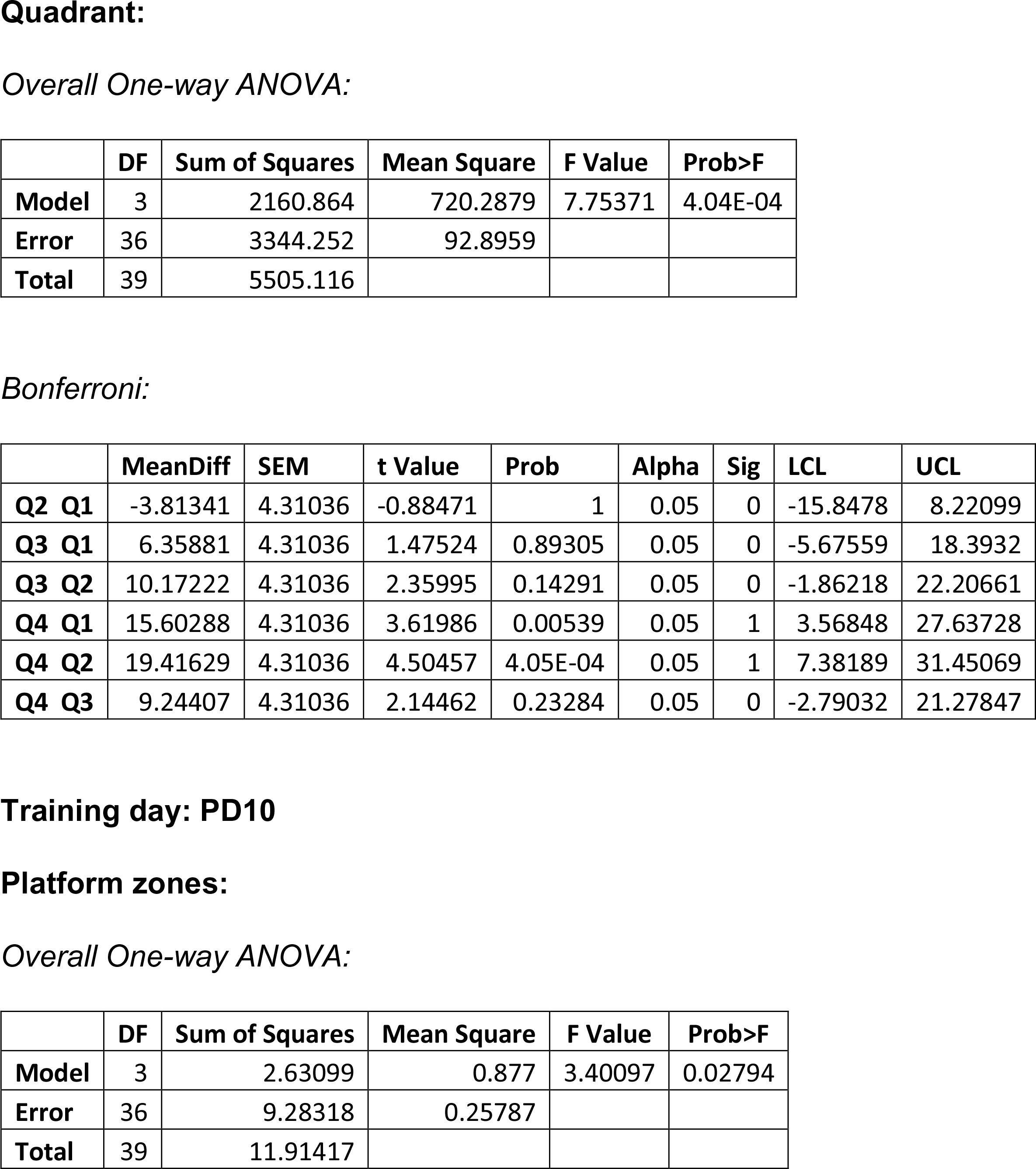
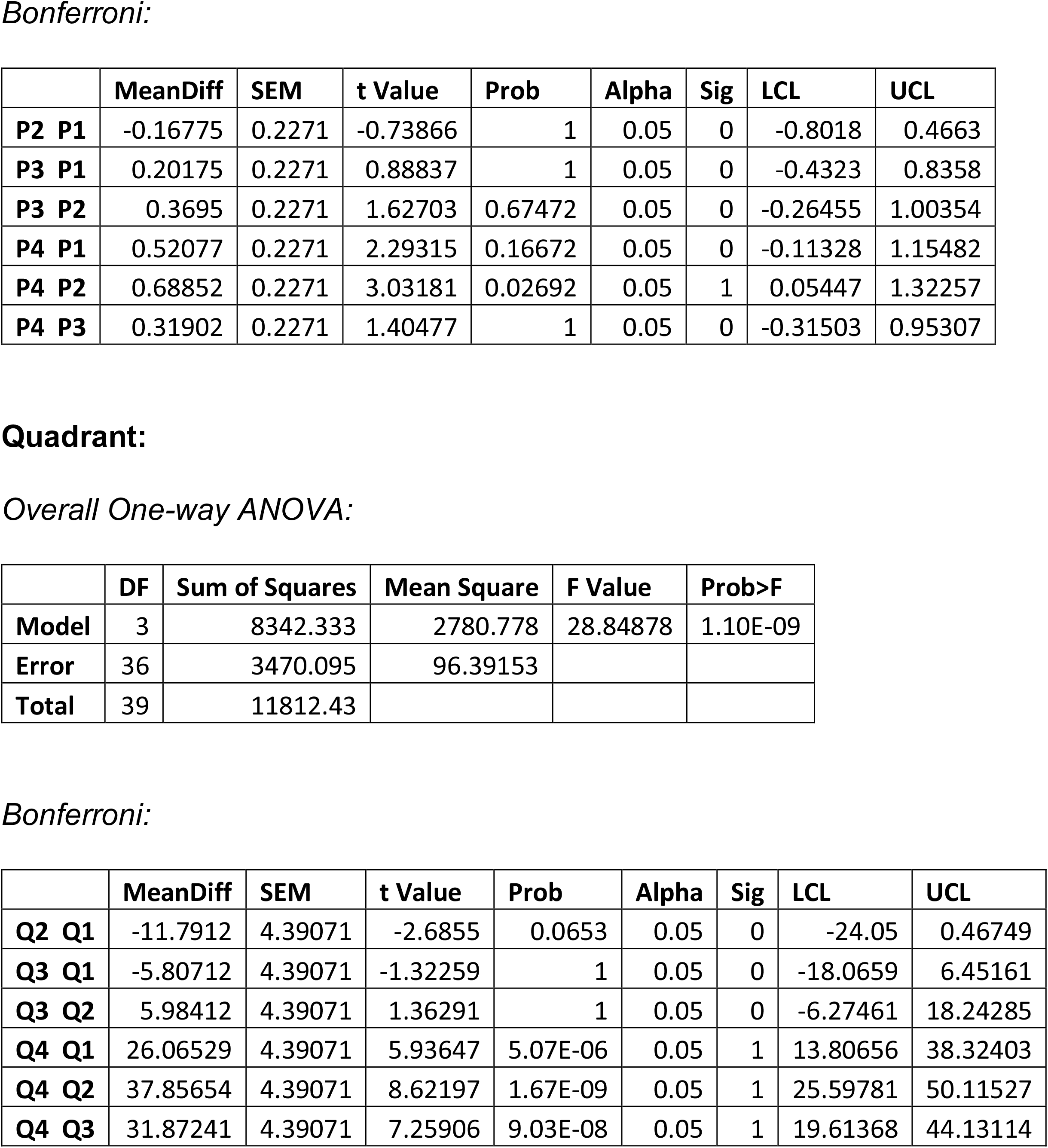
One-way ANOVA of residence time in BALB/cJ (PD3, PD7, PD10)

**S. Table 18:**
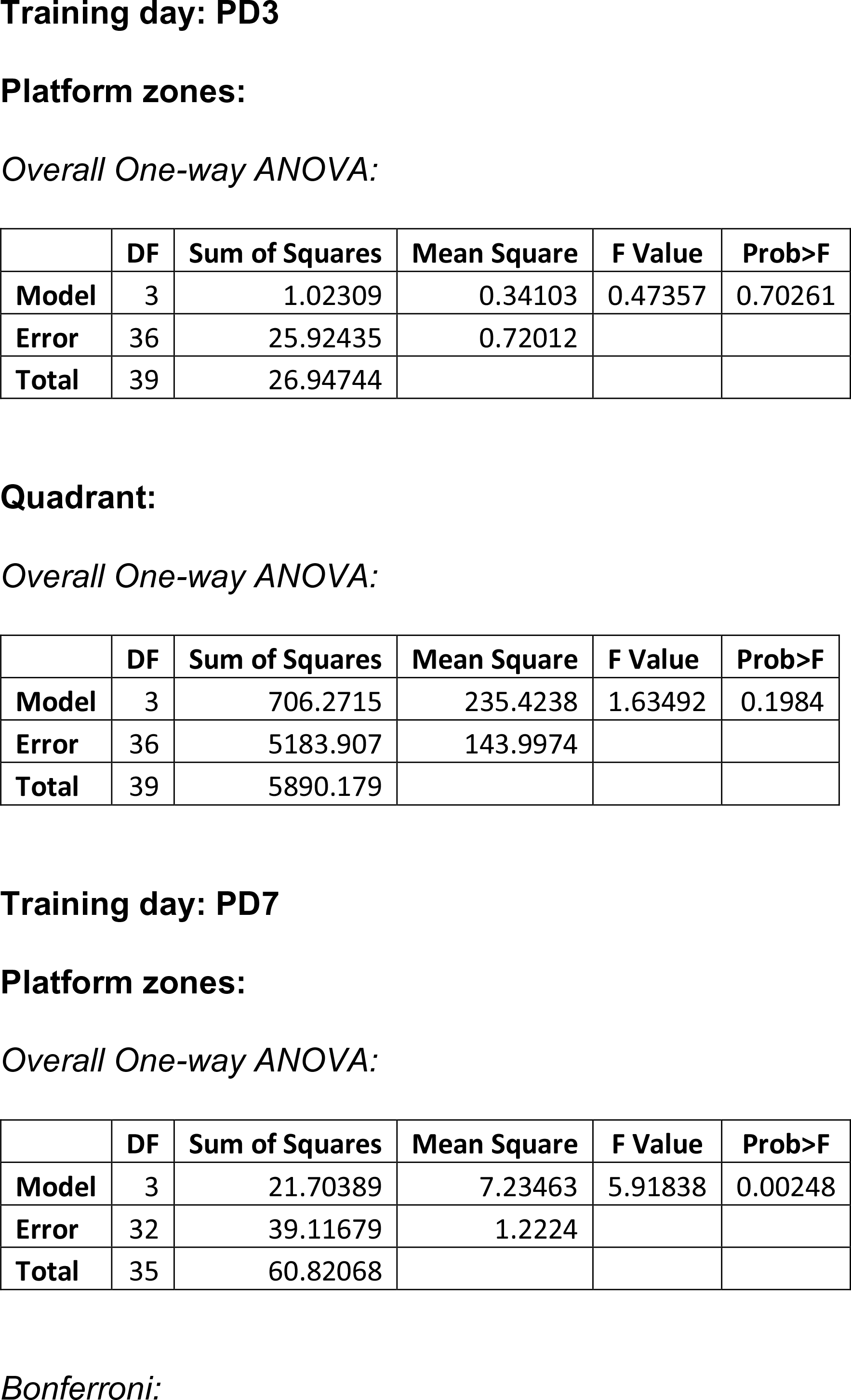
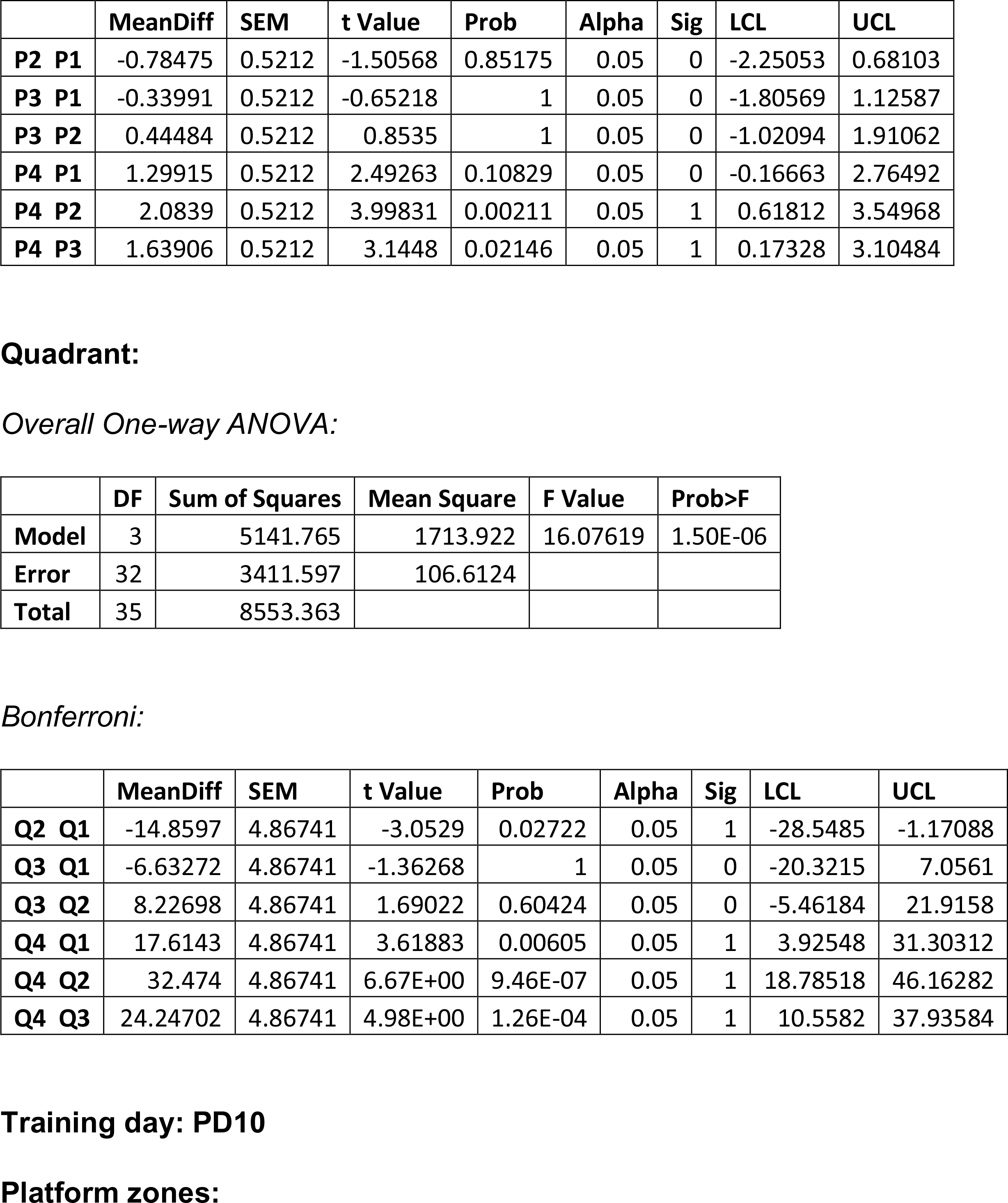
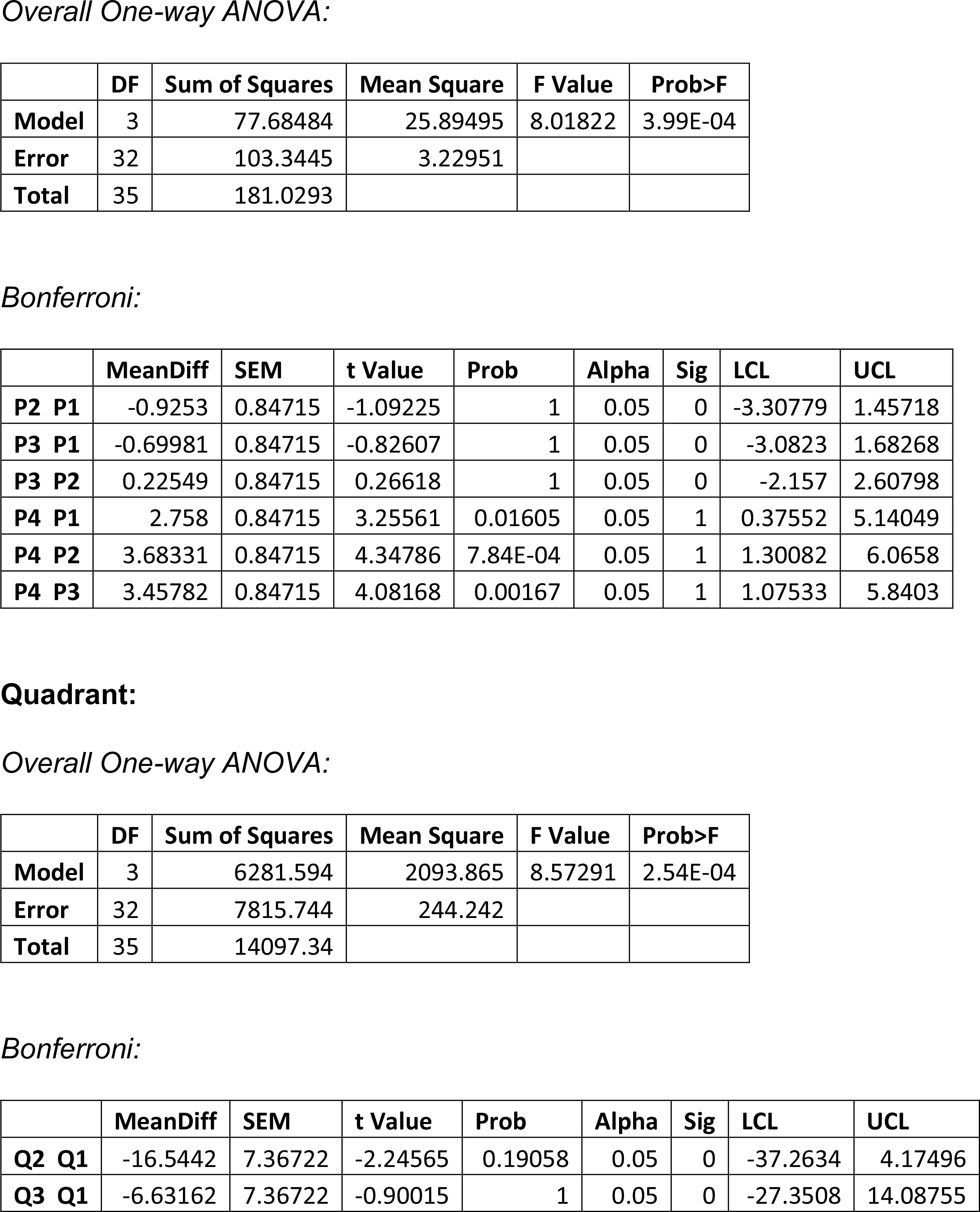
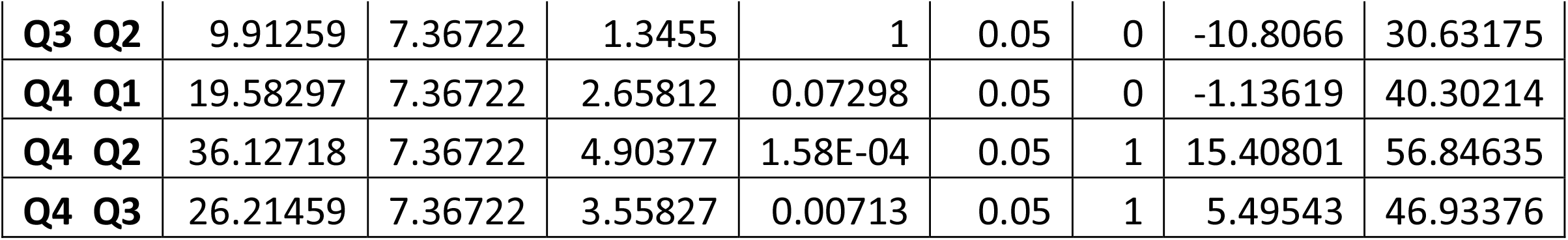
One-way ANOVA of residence time in SWR/J (PD3, PD7, PD10)

|  | DF | Sum of Squares | Mean Square | F Value | Prob>F |
| --- | --- | --- | --- | --- | --- |
| Model | 3 | 1.02309 | 0.34103 | 0.47357 | 0.70261 |
| Error | 36 | 25.92435 | 0.72012 |  |  |
| Total | 39 | 26.94744 |  |  |  |

Quadrant: Overall One-way ANOVA:
|  | DF | Sum of Squares | Mean Square | F Value | Prob>F |
| --- | --- | --- | --- | --- | --- |
| Model | 3 | 706.2715 | 235.4238 | 1.63492 | 0.1984 |
| Error | 36 | 5183.907 | 143.9974 |  |  |
| Total | 39 | 5890.179 |  |  |  |

|  | DF | Sum of Squares | Mean Square | F Value | Prob>F |
| --- | --- | --- | --- | --- | --- |
| Model | 3 | 21.70389 | 7.23463 | 5.91838 | 0.00248 |
| Error | 32 | 39.11679 | 1.2224 |  |  |
| Total | 35 | 60.82068 |  |  |  |

Bonferroni:
|  | MeanDiff | SEM | t Value | Prob | Alpha | Sig | LCL | UCL |
| --- | --- | --- | --- | --- | --- | --- | --- | --- |
| **P2 P1** | -0.78475 | 0.5212 | -1.50568 | 0.85175 | 0.05 | 0 | -2.25053 | 0.68103 |
| **P3 P1** | -0.33991 | 0.5212 | -0.65218 | 1 | 0.05 | 0 | -1.80569 | 1.12587 |
| **P3 P2** | 0.44484 | 0.5212 | 0.8535 | 1 | 0.05 | 0 | -1.02094 | 1.91062 |
| **P4 P1** | 1.29915 | 0.5212 | 2.49263 | 0.10829 | 0.05 | 0 | -0.16663 | 2.76492 |
| **P4 P2** | 2.0839 | 0.5212 | 3.99831 | 0.00211 | 0.05 | 1 | 0.61812 | 3.54968 |
| **P4 P3** | 1.63906 | 0.5212 | 3.1448 | 0.02146 | 0.05 | 1 | 0.17328 | 3.10484 |

**Quadrant:** *Overall One-way ANOVA:*
|  | DF | Sum of Squares | Mean Square | F Value | Prob>F |
| --- | --- | --- | --- | --- | --- |
| **Model** | 3 | 5141.765 | 1713.922 | 16.07619 | 1.50E-06 |
| **Error** | 32 | 3411.597 | 106.6124 |  |  |
| **Total** | 35 | 8553.363 |  |  |  |

*Bonferroni:*
|  | MeanDiff | SEM | t Value | Prob | Alpha | Sig | LCL | UCL |
| --- | --- | --- | --- | --- | --- | --- | --- | --- |
| **Q2 Q1** | -14.8597 | 4.86741 | -3.0529 | 0.02722 | 0.05 | 1 | -28.5485 | -1.17088 |
| **Q3 Q1** | -6.63272 | 4.86741 | -1.36268 | 1 | 0.05 | 0 | -20.3215 | 7.0561 |
| **Q3 Q2** | 8.22698 | 4.86741 | 1.69022 | 0.60424 | 0.05 | 0 | -5.46184 | 21.9158 |
| **Q4 Q1** | 17.6143 | 4.86741 | 3.61883 | 0.00605 | 0.05 | 1 | 3.92548 | 31.30312 |
| **Q4 Q2** | 32.474 | 4.86741 | 6.67E+00 | 9.46E-07 | 0.05 | 1 | 18.78518 | 46.16282 |
| **Q4 Q3** | 24.24702 | 4.86741 | 4.98E+00 | 1.26E-04 | 0.05 | 1 | 10.5582 | 37.93584 |
| **Model** | 3 | 77.68484 | 25.89495 | 8.01822 | 3.99E-04 |
| **Error** | 32 | 103.3445 | 3.22951 |  |  |
| **Total** | 35 | 181.0293 |  |  |  |

Bonferroni:
|  | MeanDiff | SEM | t Value | Prob | Alpha | Sig | LCL | UCL |
| --- | --- | --- | --- | --- | --- | --- | --- | --- |
| **P2 P1** | -0.9253 | 0.84715 | -1.09225 | 1 | 0.05 | 0 | -3.30779 | 1.45718 |
| **P3 P1** | -0.69981 | 0.84715 | -0.82607 | 1 | 0.05 | 0 | -3.0823 | 1.68268 |
| **P3 P2** | 0.22549 | 0.84715 | 0.26618 | 1 | 0.05 | 0 | -2.157 | 2.60798 |
| **P4 P1** | 2.758 | 0.84715 | 3.25561 | 0.01605 | 0.05 | 1 | 0.37552 | 5.14049 |
| **P4 P2** | 3.68331 | 0.84715 | 4.34786 | 7.84E-04 | 0.05 | 1 | 1.30082 | 6.0658 |
| **P4 P3** | 3.45782 | 0.84715 | 4.08168 | 0.00167 | 0.05 | 1 | 1.07533 | 5.8403 |

Quadrant: Overall One-way ANOVA:
|  | DF | Sum of Squares | Mean Square | F Value | Prob>F |
| --- | --- | --- | --- | --- | --- |
| **Model** | 3 | 6281.594 | 2093.865 | 8.57291 | 2.54E-04 |
| **Error** | 32 | 7815.744 | 244.242 |  |  |
| **Total** | 35 | 14097.34 |  |  |  |

Bonferroni:
|  | MeanDiff | SEM | t Value | Prob | Alpha | Sig | LCL | UCL |
| --- | --- | --- | --- | --- | --- | --- | --- | --- |
| **Q2 Q1** | -16.5442 | 7.36722 | -2.24565 | 0.19058 | 0.05 | 0 | -37.2634 | 4.17496 |
| **Q3 Q1** | -6.63162 | 7.36722 | -0.90015 | 1 | 0.05 | 0 | -27.3508 | 14.08755 |
| **Q3 Q2** | 9.91259 | 7.36722 | 1.3455 | 1 | 0.05 | 0 | -10.8066 | 30.63175 |
| **Q4 Q1** | 19.58297 | 7.36722 | 2.65812 | 0.07298 | 0.05 | 0 | -1.13619 | 40.30214 |
| **Q4 Q2** | 36.12718 | 7.36722 | 4.90377 | 1.58E-04 | 0.05 | 1 | 15.40801 | 56.84635 |
| **Q4 Q3** | 26.21459 | 7.36722 | 3.55827 | 0.00713 | 0.05 | 1 | 5.49543 | 46.93376 |

**S. Table 19:** One-way ANOVA of residence time in *Ptpn11* +/+ (PD3, PD5, PD7)

|  | DF | Sum of Squares | Mean Square | F Value | Prob>F |
| --- | --- | --- | --- | --- | --- |
| **Model** | 3 | 13.37722 | 4.45907 | 9.36383 | 4.11E-05 |
| **Error** | 56 | 26.66729 | 0.4762 |  |  |
| **Total** | 59 | 40.04451 |  |  |  |

Bonferroni:
|  | MeanDiff | SEM | t Value | Prob | Alpha | Sig | LCL | UCL |
| --- | --- | --- | --- | --- | --- | --- | --- | --- |
| **P2 P1** | 0.2995 | 0.25198 | 1.1886 | 1 | 0.05 | 0 | -0.38972 | 0.98872 |
| **P3 P1** | -0.16894 | 0.25198 | -0.67045 | 1 | 0.05 | 0 | -0.85816 | 0.52028 |
| **P3 P2** | -0.46844 | 0.25198 | -1.85905 | 0.40967 | 0.05 | 0 | -1.15766 | 0.22078 |
| **P4 P1** | 1.06284 | 0.25198 | 4.21796 | 5.46E-04 | 0.05 | 1 | 0.37362 | 1.75206 |
| **P4 P2** | 0.76333 | 0.25198 | 3.02936 | 0.02222 | 0.05 | 1 | 0.07412 | 1.45255 |
| **P4 P3** | 1.23178 | 0.25198 | 4.8884 | 5.35E-05 | 0.05 | 1 | 0.54256 | 1.921 |

Quadrant: Overall One-way ANOVA:
|  | DF | Sum of Squares | Mean Square | F Value | Prob>F |
| --- | --- | --- | --- | --- | --- |
| **Model** | 3 | 3443.372 | 1147.791 | 22.70877 | 9.52E-10 |
| **Error** | 56 | 2830.46 | 50.54393 |  |  |
| **Total** | 59 | 6273.832 |  |  |  |

*Bonferroni:*
|  | MeanDiff | SEM | t Value | Prob | Alpha | Sig | LCL | UCL |
| --- | --- | --- | --- | --- | --- | --- | --- | --- |
| **Q2 Q1** | 2.65812 | 2.596 | 1.02393 | 1 | 0.05 | 0 | -4.4425 | 9.75874 |
| **Q3 Q1** | -5.38142 | 2.596 | -2.07297 | 0.25673 | 0.05 | 0 | -12.482 | 1.7192 |
| **Q3 Q2** | -8.03954 | 2.596 | -3.0969 | 0.01832 | 0.05 | 1 | -15.1402 | -0.93892 |
| **Q4 Q1** | 15.25824 | 2.596 | 5.87761 | 1.44E-06 | 0.05 | 1 | 8.15762 | 22.35886 |
| **Q4 Q2** | 12.60012 | 2.596 | 4.85368 | 6.05E-05 | 0.05 | 1 | 5.4995 | 19.70074 |
| **Q4 Q3** | 20.63967 | 2.596 | 7.95058 | 5.59E-10 | 0.05 | 1 | 13.53905 | 27.74028 |

|  | DF | Sum of Squares | Mean Square | F Value | Prob>F |
| --- | --- | --- | --- | --- | --- |
| **Model** | 3 | 32.1798 | 10.7266 | 12.37014 | 2.57E-06 |
| **Error** | 56 | 48.55965 | 0.86714 |  |  |
| **Total** | 59 | 80.73945 |  |  |  |

*Bonferroni:*
|  | MeanDiff | SEM | t Value | Prob | Alpha | Sig | LCL | UCL |
| --- | --- | --- | --- | --- | --- | --- | --- | --- |
| **P2 P1** | 0.36778 | 0.34003 | 1.08162 | 1 | 0.05 | 0 | -0.56227 | 1.29783 |
| **P3 P1** | 0.09652 | 0.34003 | 0.28385 | 1 | 0.05 | 0 | -0.83353 | 1.02656 |
| **P3 P2** | -0.27127 | 0.34003 | -0.79778 | 1 | 0.05 | 0 | -1.20131 | 0.65878 |
| **P4 P1** | 1.81713 | 0.34003 | 5.34409 | 1.03E-05 | 0.05 | 1 | 0.88708 | 2.74718 |
| **P4 P2** | 1.44935 | 0.34003 | 4.26246 | 4.70E-04 | 0.05 | 1 | 0.5193 | 2.3794 |
| **P4 P3** | 1.72062 | 0.34003 | 5.06024 | 2.90E-05 | 0.05 | 1 | 0.79057 | 2.65067 |

**Quadrant:** *Overall One-way ANOVA:*
|  | DF | Sum of Squares | Mean Square | F Value | Prob>F |
| --- | --- | --- | --- | --- | --- |
| **Model** | 3 | 5163.906 | 1721.302 | 16.91657 | 6.07E-08 |
| **Error** | 56 | 5698.135 | 101.7524 |  |  |
| **Total** | 59 | 10862.04 |  |  |  |

*Bonferroni:*
|  | MeanDiff | SEM | t Value | Prob | Alpha | Sig | LCL | UCL |
| --- | --- | --- | --- | --- | --- | --- | --- | --- |
| **Q2 Q1** | 5.6225 | 3.68334 | 1.52647 | 0.79513 | 0.05 | 0 | -4.45225 | 15.69724 |
| **Q3 Q1** | 3.81986 | 3.68334 | 1.03706 | 1 | 0.05 | 0 | -6.25489 | 13.89461 |
| **Q3 Q2** | -1.80264 | 3.68334 | -0.4894 | 1 | 0.05 | 0 | -11.8774 | 8.27211 |
| **Q4 Q1** | 24.05282 | 3.68334 | 6.53017 | 1.24E-07 | 0.05 | 1 | 13.97808 | 34.12757 |
| **Q4 Q2** | 18.43033 | 3.68334 | 5.0037 | 3.55E-05 | 0.05 | 1 | 8.35558 | 28.50507 |
| **Q4 Q3** | 20.23296 | 3.68334 | 5.4931 | 5.99E-06 | 0.05 | 1 | 10.15822 | 30.30771 |

|  | DF | Sum of Squares | Mean Square | F Value | Prob>F |
| --- | --- | --- | --- | --- | --- |
| **Model** | 3 | 112.9265 | 37.64218 | 15.70201 | 1.58E-07 |
| **Error** | 56 | 134.2479 | 2.39728 |  |  |
| **Total** | 59 | 247.1745 |  |  |  |

*Bonferroni:*
|  | MeanDiff | SEM | t Value | Prob | Alpha | Sig | LCL | UCL |
| --- | --- | --- | --- | --- | --- | --- | --- | --- |
| **P2 P1** | 0.10764 | 0.56537 | 0.19038 | 1 | 0.05 | 0 | -1.43876 | 1.65404 |
| **P3 P1** | 0.20613 | 0.56537 | 0.36459 | 1 | 0.05 | 0 | -1.34027 | 1.75253 |
| **P3 P2** | 0.09849 | 0.56537 | 0.17421 | 1 | 0.05 | 0 | -1.44791 | 1.64489 |
| **P4 P1** | 3.26838 | 0.56537 | 5.781 | 2.07E-06 | 0.05 | 1 | 1.72198 | 4.81478 |
| **P4 P2** | 3.16074 | 0.56537 | 5.59062 | 4.18E-06 | 0.05 | 1 | 1.61434 | 4.70714 |
| **P4 P3** | 3.06225 | 0.56537 | 5.41641 | 7.94E-06 | 0.05 | 1 | 1.51585 | 4.60865 |

**Quadrant:** *Overall One-way ANOVA:*
|  | DF | Sum of Squares | Mean Square | F Value | Prob>F |
| --- | --- | --- | --- | --- | --- |
| **Model** | 3 | 9295.567 | 3098.522 | 35.35832 | 5.90E-13 |
| **Error** | 56 | 4907.395 | 87.63206 |  |  |
| **Total** | 59 | 14202.96 |  |  |  |

*Bonferroni:*
|  | MeanDiff | SEM | t Value | Prob | Alpha | Sig | LCL | UCL |
| --- | --- | --- | --- | --- | --- | --- | --- | --- |
| **Q2 Q1** | 0.62958 | 3.41823 | 0.18418 | 1 | 0.05 | 0 | -8.72003 | 9.97918 |
| **Q3 Q1** | 1.04187 | 3.41823 | 0.3048 | 1 | 0.05 | 0 | -8.30773 | 10.39148 |
| **Q3 Q2** | 0.4123 | 3.41823 | 0.12062 | 1 | 0.05 | 0 | -8.93731 | 9.7619 |
| **Q4 Q1** | 29.28933 | 3.41823 | 8.56858 | 5.43E-11 | 0.05 | 1 | 19.93973 | 38.63894 |
| **Q4 Q2** | 28.65976 | 3.41823 | 8.38439 | 1.09E-10 | 0.05 | 1 | 19.31015 | 38.00936 |
| **Q4 Q3** | 28.24746 | 3.41823 | 8.26378 | 1.71E-10 | 0.05 | 1 | 18.89786 | 37.59707 |

**S. Table 20:** One-way ANOVA of residence time in *Ptpn11* D61G/+ (PD3, PD5, PD7)

|  | DF | Sum of Squares | Mean Square | F Value | Prob>F |
| --- | --- | --- | --- | --- | --- |
| **Model** | 3 | 2.6682 | 0.8894 | 0.82426 | 0.48915 |
| **Error** | 36 | 38.84521 | 1.07903 |  |  |
| **Total** | 39 | 41.51341 |  |  |  |

**Quadrant:** *Overall One-way ANOVA:*
|  | DF | Sum of Squares | Mean Square | F Value | Prob>F |
| --- | --- | --- | --- | --- | --- |
| **Model** | 3 | 591.4742 | 197.1581 | 1.97745 | 0.13475 |
| **Error** | 36 | 3589.308 | 99.70299 |  |  |
| **Total** | 39 | 4180.782 |  |  |  |

|  | DF | Sum of Squares | Mean Square | F Value | Prob>F |
| --- | --- | --- | --- | --- | --- |
| **Model** | 3 | 4.72747 | 1.57582 | 0.92963 | 0.43637 |
| **Error** | 36 | 61.02423 | 1.69512 |  |  |
| Total | 39 | 65.7517 |  |  |  |

Quadrant: Overall One-way ANOVA:
|  | DF | Sum of Squares | Mean Square | F Value | Prob>F |
| --- | --- | --- | --- | --- | --- |
| Model | 3 | 705.9599 | 235.32 | 1.30865 | 0.28658 |
| Error | 36 | 6473.501 | 179.8195 |  |  |
| Total | 39 | 7179.461 |  |  |  |

|  | DF | Sum of Squares | Mean Square | F Value | Prob>F |
| --- | --- | --- | --- | --- | --- |
| Model | 3 | 10.79882 | 3.59961 | 2.85701 | 0.05051 |
| Error | 36 | 45.35719 | 1.25992 |  |  |
| Total | 39 | 56.15601 |  |  |  |

Quadrant: Overall One-way ANOVA:
|  | DF | Sum of Squares | Mean Square | F Value | Prob>F |
| --- | --- | --- | --- | --- | --- |
| Model | 3 | 1304.115 | 434.7049 | 1.85952 | 0.15394 |
| Error | 36 | 8415.821 | 233.7728 |  |  |
| Total | 39 | 9719.935 |  |  |  |

**S. Table 21:** One-way ANOVA of residence time in *Ptpn11* +/+ (Veh and SL327)

|  | DF | Sum of Squares | Mean Square | F Value | Prob>F |
| --- | --- | --- | --- | --- | --- |
| Model | 3 | 44.0418 | 14.6806 | 14.0022 | 2.19E-06 |
| Error | 40 | 41.93797 | 1.04845 |  |  |
| Total | 43 | 85.97977 |  |  |  |

Bonferroni:
|  | MeanDiff | SEM | t Value | Prob | Alpha | Sig | LCL | UCL |
| --- | --- | --- | --- | --- | --- | --- | --- | --- |
| P2 P1 | -0.22735 | 0.43661 | -0.52071 | 1 | 0.05 | 0 | -1.43932 | 0.98462 |
| P3 P1 | 1.76E-04 | 0.43661 | 4.04E-04 | 1 | 0.05 | 0 | -1.21179 | 1.21214 |
| P3 P2 | 0.22752 | 0.43661 | 0.52112 | 1 | 0.05 | 0 | -0.98444 | 1.43949 |
| P4 P1 | 2.2248 | 0.43661 | 5.09564 | 5.24E-05 | 0.05 | 1 | 1.01283 | 3.43677 |
| P4 P2 | 2.45215 | 0.43661 | 5.61636 | 9.81E-06 | 0.05 | 1 | 1.24018 | 3.66412 |
| P4 P3 | 2.22463 | 0.43661 | 5.09524 | 5.24E-05 | 0.05 | 1 | 1.01266 | 3.43659 |

Quadrant: Overall One-way ANOVA:
|  | DF | Sum of Squares | Mean Square | F Value | Prob>F |
| --- | --- | --- | --- | --- | --- |
| Model | 3 | 11546.21 | 3848.735 | 49.60291 | 1.52E-13 |
| Error | 40 | 3103.637 | 77.59091 |  |  |
| Total | 43 | 14649.84 |  |  |  |

*Bonferroni:*
|  | MeanDiff | SEM | t Value | Prob | Alpha | Sig | LCL | UCL |
| --- | --- | --- | --- | --- | --- | --- | --- | --- |
| **Q2 Q1** | -7.04366 | 3.75599 | -1.87532 | 0.40837 | 0.05 | 0 | -17.4698 | 3.38245 |
| **Q3 Q1** | -1.93791 | 3.75599 | -0.51595 | 1 | 0.05 | 0 | -12.364 | 8.48821 |
| **Q3 Q2** | 5.10576 | 3.75599 | 1.35937 | 1 | 0.05 | 0 | -5.32036 | 15.53188 |
| **Q4 Q1** | 33.94169 | 3.75599 | 9.03669 | 1.98E-10 | 0.05 | 1 | 23.51557 | 44.36781 |
| **Q4 Q2** | 40.98535 | 3.75599 | 10.91201 | 8.81E-13 | 0.05 | 1 | 30.55923 | 51.41147 |
| **Q4 Q3** | 35.87959 | 3.75599 | 9.55264 | 4.26E-11 | 0.05 | 1 | 25.45348 | 46.30571 |

|  | DF | Sum of Squares | Mean Square | F Value | Prob>F |
| --- | --- | --- | --- | --- | --- |
| **Model** | 3 | 24.46334 | 8.15445 | 6.63545 | 9.62E-04 |
| **Error** | 40 | 49.15683 | 1.22892 |  |  |
| **Total** | 43 | 73.62017 |  |  |  |

*Bonferroni:*
|  | MeanDiff | SEM | t Value | Prob | Alpha | Sig | LCL | UCL |
| --- | --- | --- | --- | --- | --- | --- | --- | --- |
| **P2 P1** | -0.5145 | 0.47269 | -1.08843 | 1 | 0.05 | 0 | -1.82663 | 0.79764 |
| **P3 P1** | -0.84748 | 0.47269 | -1.79286 | 0.48336 | 0.05 | 0 | -2.15961 | 0.46466 |
| **P3 P2** | -0.33298 | 0.47269 | -0.70443 | 1 | 0.05 | 0 | -1.64512 | 0.97916 |
| **P4 P1** | 1.12053 | 0.47269 | 2.37052 | 0.13603 | 0.05 | 0 | -0.19161 | 2.43267 |
| **P4 P2** | 1.63503 | 0.47269 | 3.45895 | 0.00781 | 0.05 | 1 | 0.32289 | 2.94716 |
| **P4 P3** | 1.96801 | 0.47269 | 4.16338 | 9.72E-04 | 0.05 | 1 | 0.65587 | 3.28014 |

Quadrant: Overall One-way ANOVA:
|  | DF | Sum of Squares | Mean Square | F Value | Prob>F |
| --- | --- | --- | --- | --- | --- |
| Model | 3 | 7333.376 | 2444.459 | 15.55633 | 7.40E-07 |
| Error | 40 | 6285.436 | 157.1359 |  |  |
| Total | 43 | 13618.81 |  |  |  |

Bonferroni:
|  | MeanDiff | SEM | t Value | Prob | Alpha | Sig | LCL | UCL |
| --- | --- | --- | --- | --- | --- | --- | --- | --- |
| Q2 Q1 | -9.2759 | 5.34511 | -1.7354 | 0.54221 | 0.05 | 0 | -24.1132 | 5.5614 |
| Q3 Q1 | -12.3508 | 5.34511 | -2.31068 | 0.15653 | 0.05 | 0 | -27.1881 | 2.48647 |
| Q3 Q2 | -3.07493 | 5.34511 | -0.57528 | 1 | 0.05 | 0 | -17.9122 | 11.76238 |
| Q4 Q1 | 20.69546 | 5.34511 | 3.87185 | 0.00234 | 0.05 | 1 | 5.85816 | 35.53276 |
| Q4 Q2 | 29.97136 | 5.34511 | 5.60725 | 1.01E-05 | 0.05 | 1 | 15.13406 | 44.80866 |
| Q4 Q3 | 33.04629 | 5.34511 | 6.18253 | 1.57E-06 | 0.05 | 1 | 18.20898 | 47.88359 |

Treatment: Veh
Platform zones:
Overall One-way ANOVA:

**S. Table 22:**
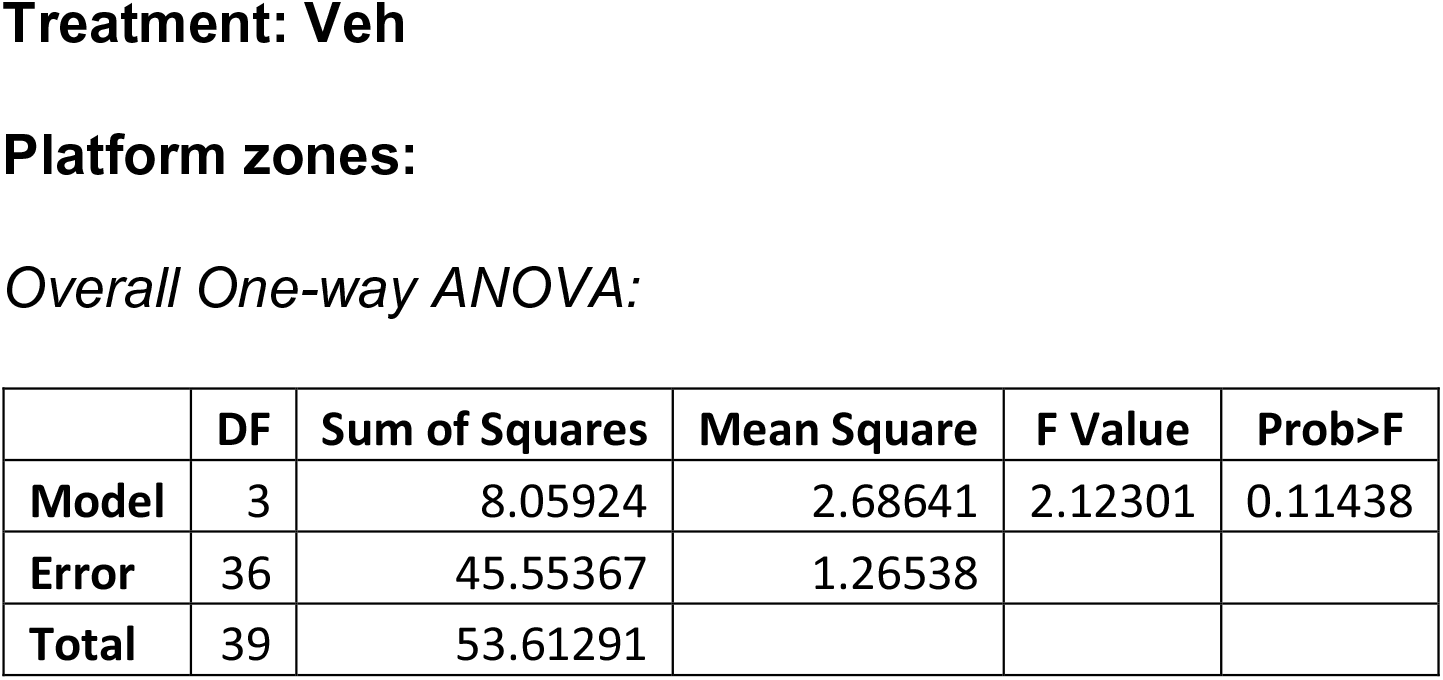
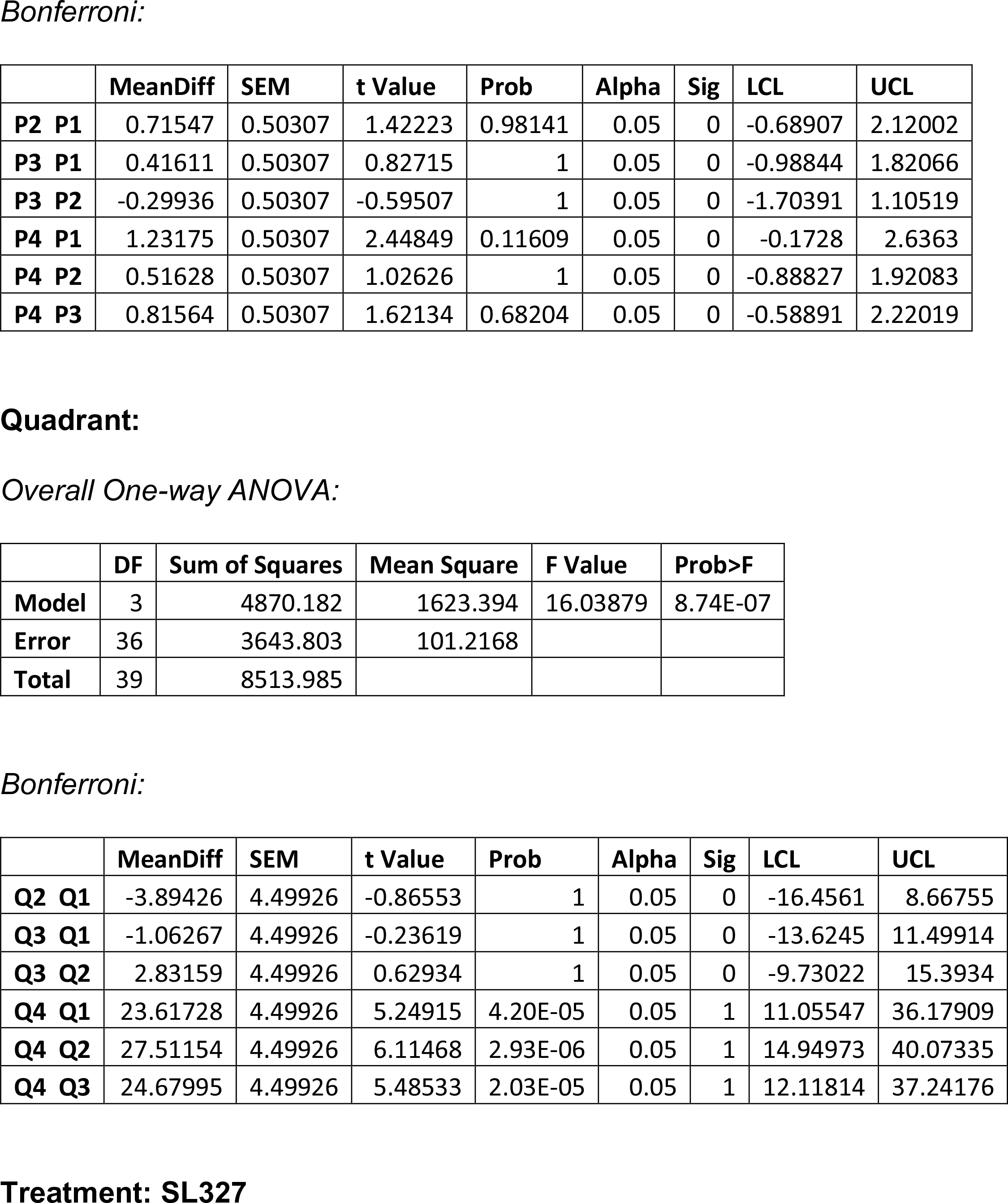
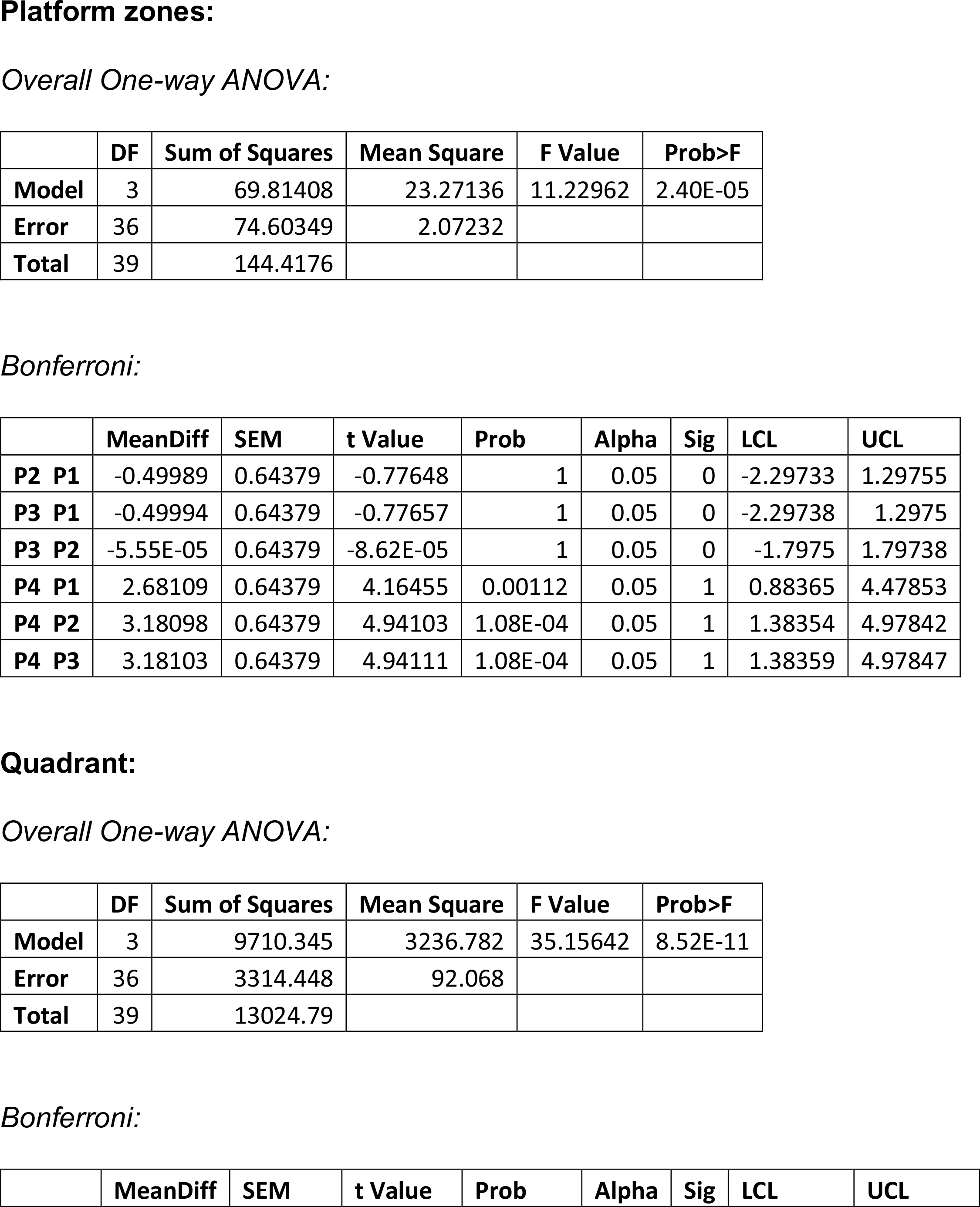
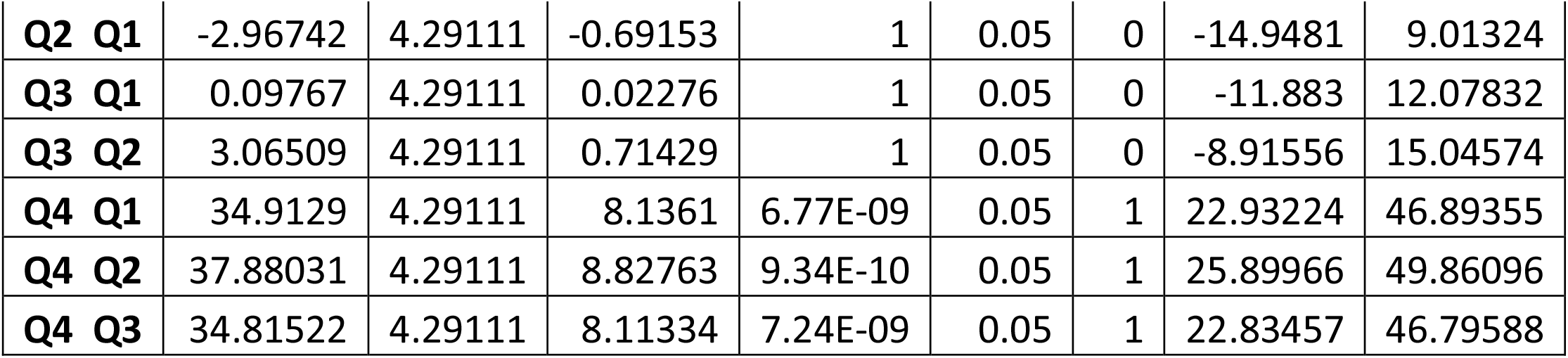
One-way ANOVA of residence time in *Ptpn11* N308D/+ (Veh and SL327)

|  | DF | Sum of Squares | Mean Square | F Value | Prob>F |
| --- | --- | --- | --- | --- | --- |
| Model | 3 | 8.05924 | 2.68641 | 2.12301 | 0.11438 |
| Error | 36 | 45.55367 | 1.26538 |  |  |
| Total | 39 | 53.61291 |  |  |  |

*Bonferroni:*
|  | MeanDiff | SEM | t Value | Prob | Alpha | Sig | LCL | UCL |
| --- | --- | --- | --- | --- | --- | --- | --- | --- |
| **P2 P1** | 0.71547 | 0.50307 | 1.42223 | 0.98141 | 0.05 | 0 | -0.68907 | 2.12002 |
| **P3 P1** | 0.41611 | 0.50307 | 0.82715 | 1 | 0.05 | 0 | -0.98844 | 1.82066 |
| **P3 P2** | -0.29936 | 0.50307 | -0.59507 | 1 | 0.05 | 0 | -1.70391 | 1.10519 |
| **P4 P1** | 1.23175 | 0.50307 | 2.44849 | 0.11609 | 0.05 | 0 | -0.1728 | 2.6363 |
| **P4 P2** | 0.51628 | 0.50307 | 1.02626 | 1 | 0.05 | 0 | -0.88827 | 1.92083 |
| **P4 P3** | 0.81564 | 0.50307 | 1.62134 | 0.68204 | 0.05 | 0 | -0.58891 | 2.22019 |

**Quadrant:** *Overall One-way ANOVA:*
|  | DF | Sum of Squares | Mean Square | F Value | Prob>F |
| --- | --- | --- | --- | --- | --- |
| **Model** | 3 | 4870.182 | 1623.394 | 16.03879 | 8.74E-07 |
| **Error** | 36 | 3643.803 | 101.2168 |  |  |
| **Total** | 39 | 8513.985 |  |  |  |

*Bonferroni:*
|  | MeanDiff | SEM | t Value | Prob | Alpha | Sig | LCL | UCL |
| --- | --- | --- | --- | --- | --- | --- | --- | --- |
| **Q2 Q1** | -3.89426 | 4.49926 | -0.86553 | 1 | 0.05 | 0 | -16.4561 | 8.66755 |
| **Q3 Q1** | -1.06267 | 4.49926 | -0.23619 | 1 | 0.05 | 0 | -13.6245 | 11.49914 |
| **Q3 Q2** | 2.83159 | 4.49926 | 0.62934 | 1 | 0.05 | 0 | -9.73022 | 15.3934 |
| **Q4 Q1** | 23.61728 | 4.49926 | 5.24915 | 4.20E-05 | 0.05 | 1 | 11.05547 | 36.17909 |
| **Q4 Q2** | 27.51154 | 4.49926 | 6.11468 | 2.93E-06 | 0.05 | 1 | 14.94973 | 40.07335 |
| **Q4 Q3** | 24.67995 | 4.49926 | 5.48533 | 2.03E-05 | 0.05 | 1 | 12.11814 | 37.24176 |
| Model | 3 | 69.81408 | 23.27136 | 11.22962 | 2.40E-05 |
| Error | 36 | 74.60349 | 2.07232 |  |  |
| Total | 39 | 144.4176 |  |  |  |

Bonferroni:
|  | MeanDiff | SEM | t Value | Prob | Alpha | Sig | LCL | UCL |
| --- | --- | --- | --- | --- | --- | --- | --- | --- |
| P2 P1 | -0.49989 | 0.64379 | -0.77648 | 1 | 0.05 | 0 | -2.29733 | 1.29755 |
| P3 P1 | -0.49994 | 0.64379 | -0.77657 | 1 | 0.05 | 0 | -2.29738 | 1.2975 |
| P3 P2 | -5.55E-05 | 0.64379 | -8.62E-05 | 1 | 0.05 | 0 | -1.7975 | 1.79738 |
| P4 P1 | 2.68109 | 0.64379 | 4.16455 | 0.00112 | 0.05 | 1 | 0.88365 | 4.47853 |
| P4 P2 | 3.18098 | 0.64379 | 4.94103 | 1.08E-04 | 0.05 | 1 | 1.38354 | 4.97842 |
| P4 P3 | 3.18103 | 0.64379 | 4.94111 | 1.08E-04 | 0.05 | 1 | 1.38359 | 4.97847 |

Quadrant: Overall One-way ANOVA:
|  | DF | Sum of Squares | Mean Square | F Value | Prob>F |
| --- | --- | --- | --- | --- | --- |
| Model | 3 | 9710.345 | 3236.782 | 35.15642 | 8.52E-11 |
| Error | 36 | 3314.448 | 92.068 |  |  |
| Total | 39 | 13024.79 |  |  |  |

Bonferroni:
|  | MeanDiff | SEM | t Value | Prob | Alpha | Sig | LCL | UCL |
| --- | --- | --- | --- | --- | --- | --- | --- | --- |
| **Q2 Q1** | -2.96742 | 4.29111 | -0.69153 | 1 | 0.05 | 0 | -14.9481 | 9.01324 |
| **Q3 Q1** | 0.09767 | 4.29111 | 0.02276 | 1 | 0.05 | 0 | -11.883 | 12.07832 |
| **Q3 Q2** | 3.06509 | 4.29111 | 0.71429 | 1 | 0.05 | 0 | -8.91556 | 15.04574 |
| **Q4 Q1** | 34.9129 | 4.29111 | 8.1361 | 6.77E-09 | 0.05 | 1 | 22.93224 | 46.89355 |
| **Q4 Q2** | 37.88031 | 4.29111 | 8.82763 | 9.34E-10 | 0.05 | 1 | 25.89966 | 49.86096 |
| **Q4 Q3** | 34.81522 | 4.29111 | 8.11334 | 7.24E-09 | 0.05 | 1 | 22.83457 | 46.79588 |

**S. Table 23:**
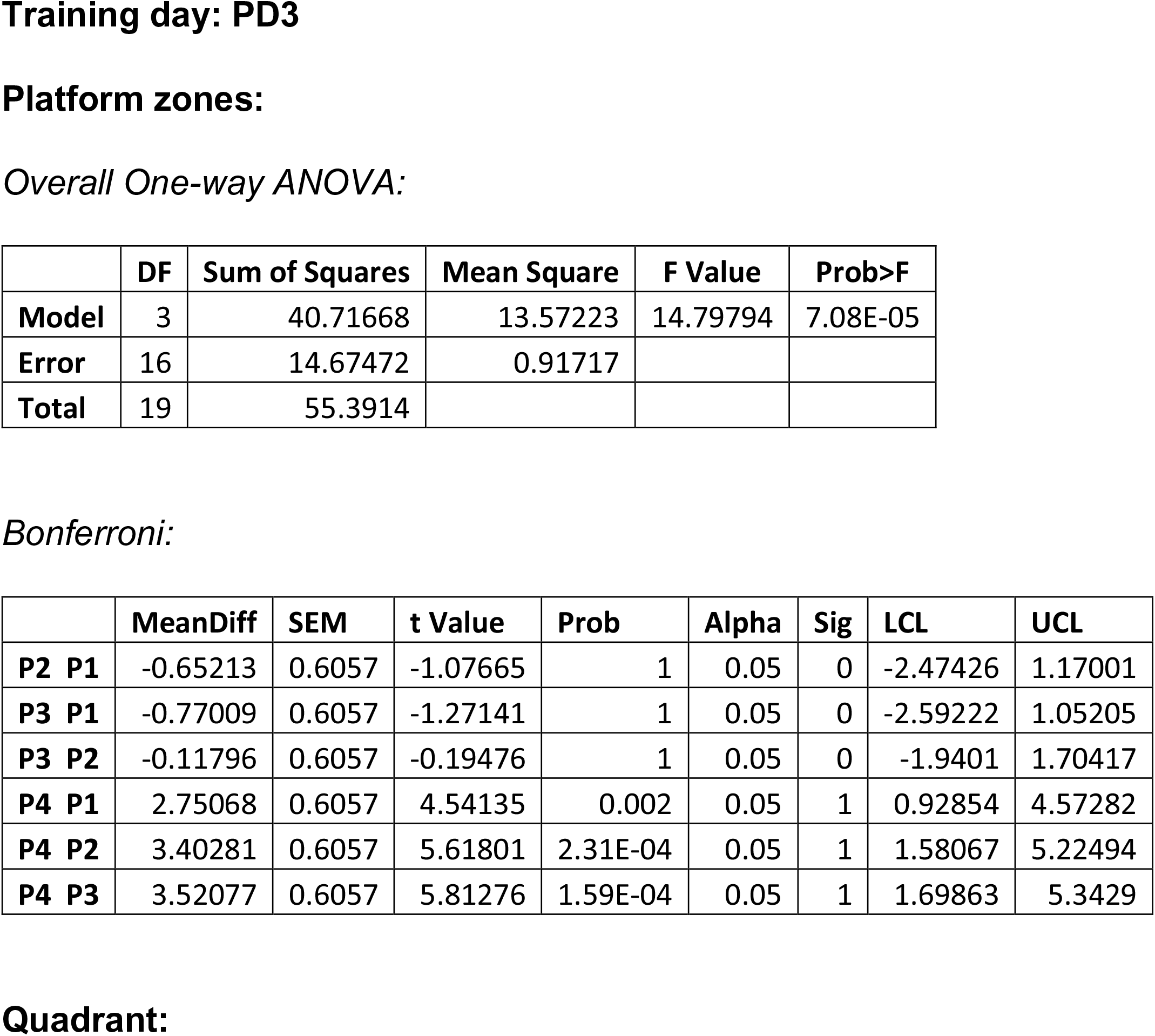
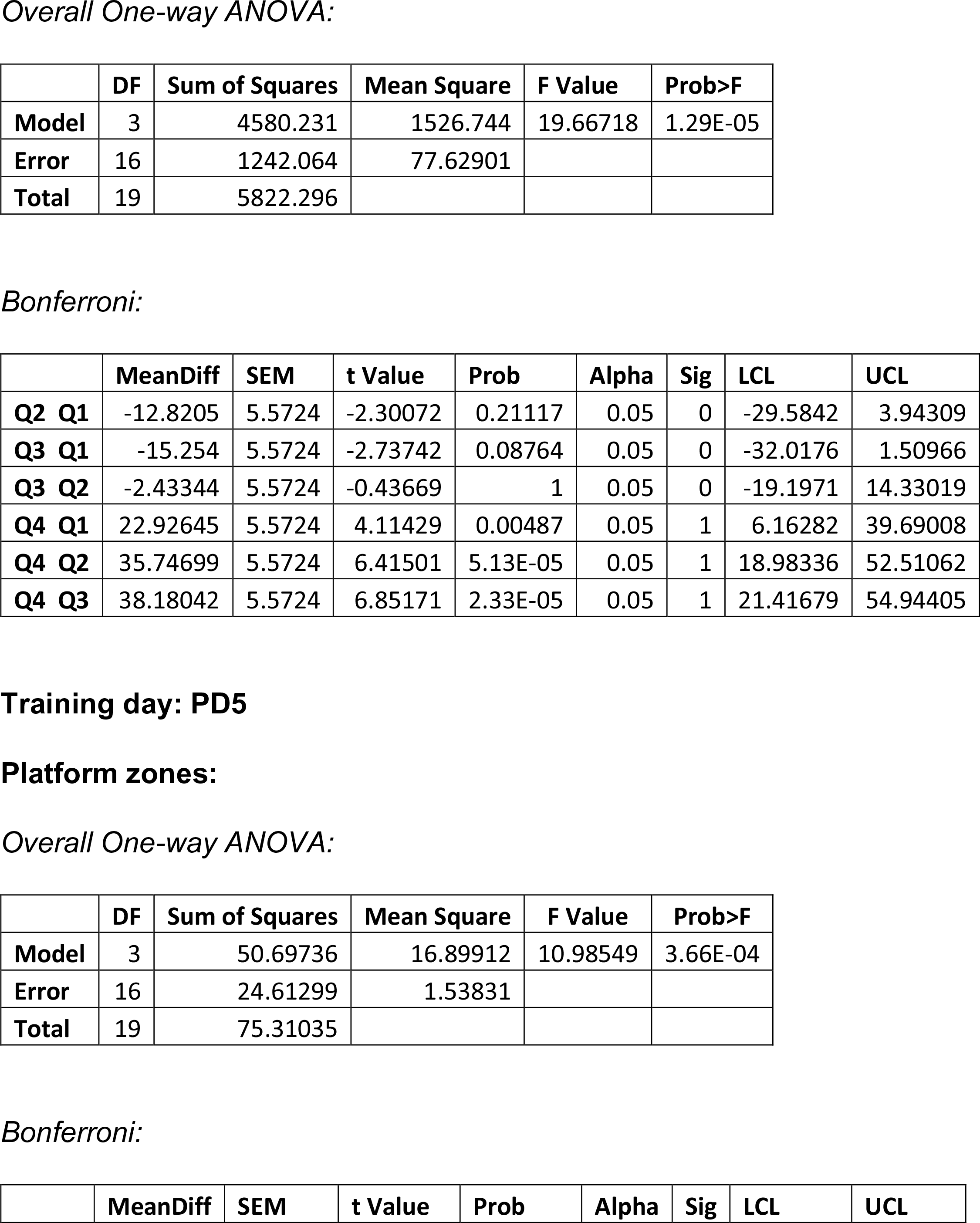
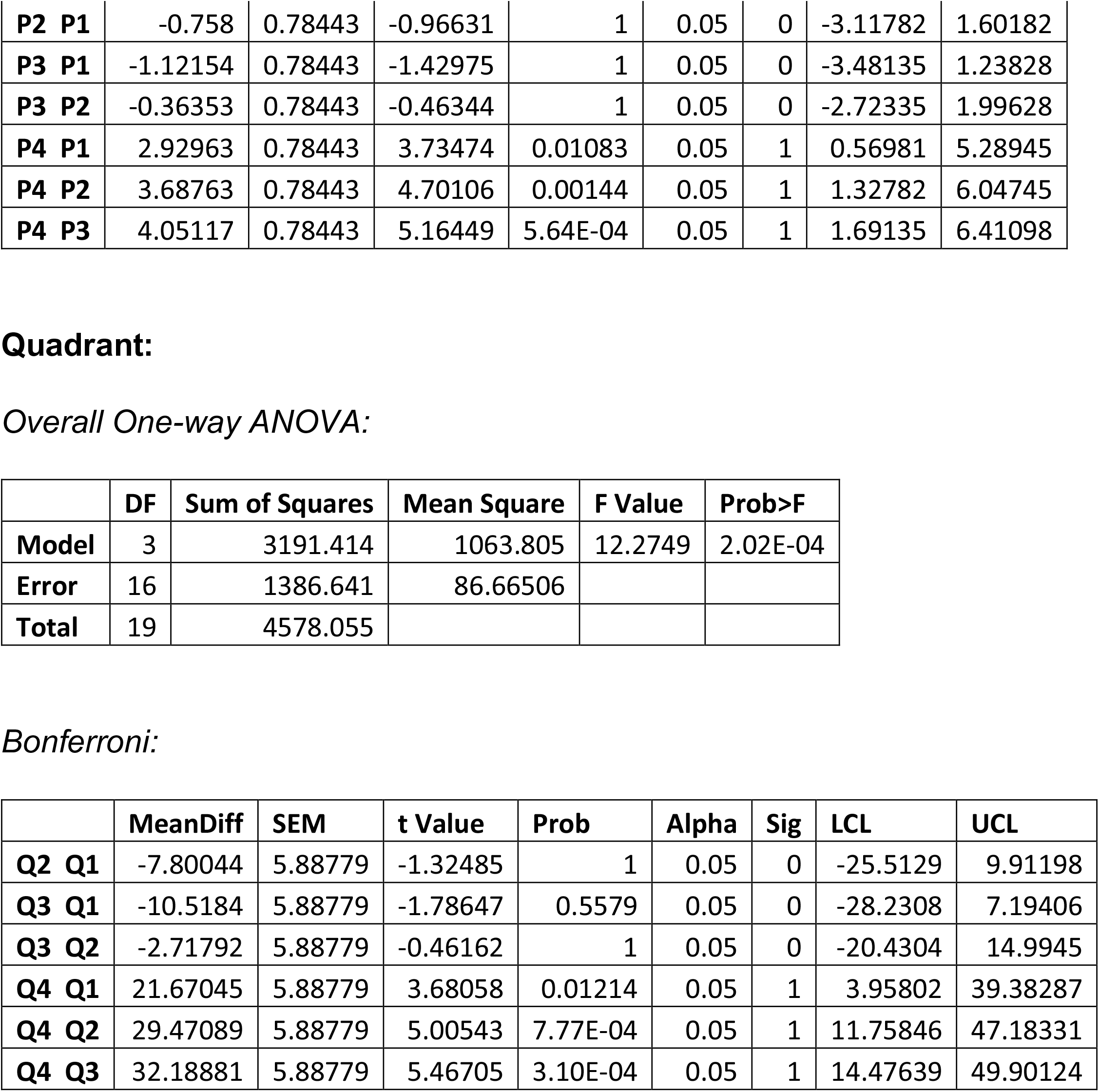
One-way ANOVA of residence time in C57Bl/6J (PD3, PD5)

**S. Table 24:** One-way ANOVA of residence time in DBA/2J (PD3, PD5)

|  | DF | Sum of Squares | Mean Square | F Value | Prob>F |
| --- | --- | --- | --- | --- | --- |
| Model | 3 | 5.51122 | 1.83707 | 2.25156 | 0.12168 |
| Error | 16 | 13.0546 | 0.81591 |  |  |
| Total | 19 | 18.56582 |  |  |  |

Quadrant: Overall One-way ANOVA:
|  | DF | Sum of Squares | Mean Square | F Value | Prob>F |
| --- | --- | --- | --- | --- | --- |
| Model | 3 | 1810.085 | 603.3615 | 1.45408 | 0.26444 |
| Error | 16 | 6639.095 | 414.9435 |  |  |
| Total | 19 | 8449.18 |  |  |  |

|  | DF | Sum of Squares | Mean Square | F Value | Prob>F |
| --- | --- | --- | --- | --- | --- |
| Model | 3 | 2.4072 | 0.8024 | 1.18561 | 0.34654 |
| Error | 16 | 10.82855 | 0.67678 |  |  |
| Total | 19 | 13.23574 |  |  |  |

Quadrant: Overall One-way ANOVA:
|  | DF | Sum of Squares | Mean Square | F Value | Prob>F |
| --- | --- | --- | --- | --- | --- |
| Model | 3 | 388.9418 | 129.6473 | 0.5827 | 0.6349 |
| Error | 16 | 3559.875 | 222.4922 |  |  |
| Total | 19 | 3948.816 |  |  |  |

**S. Table 25:**
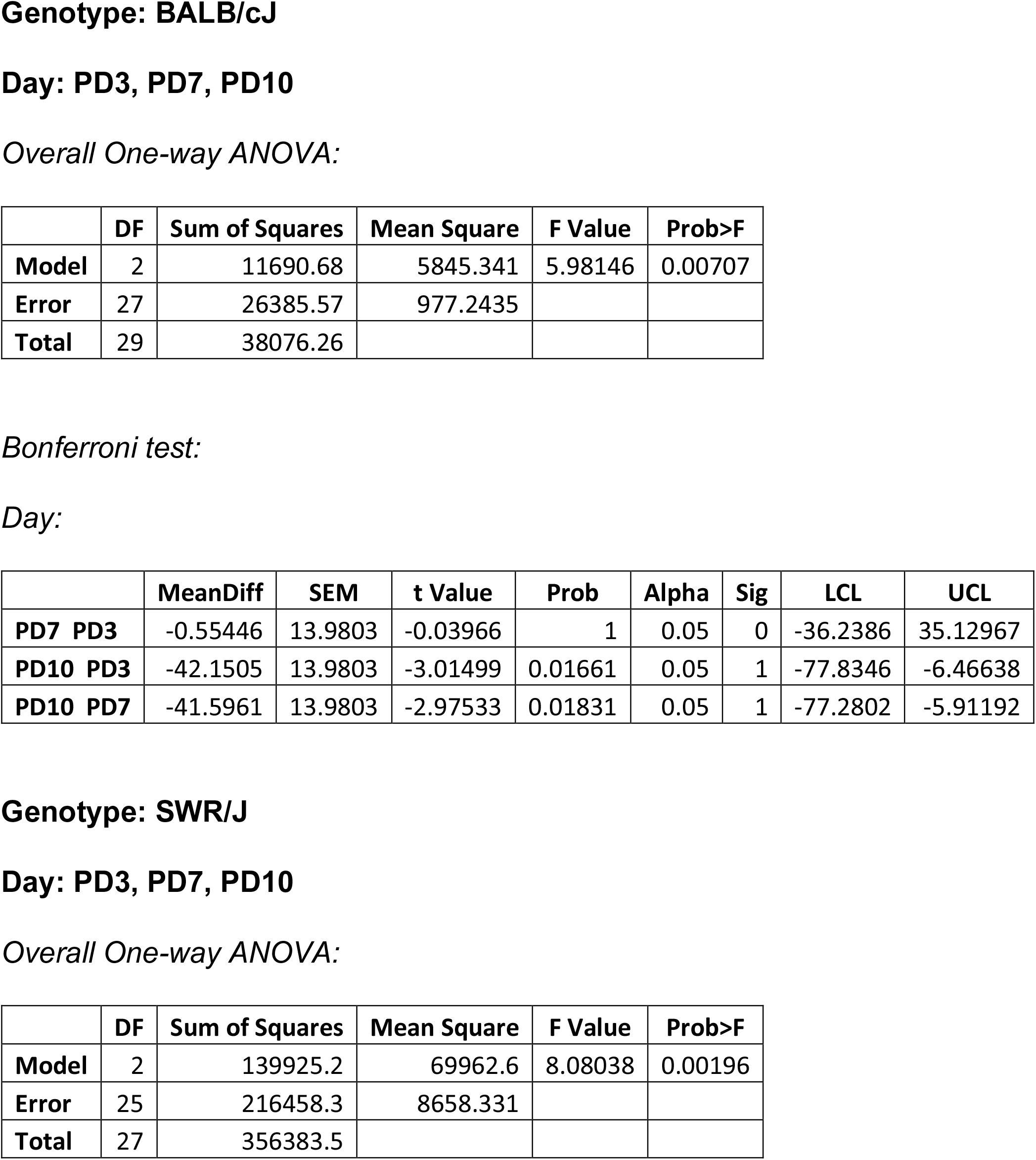
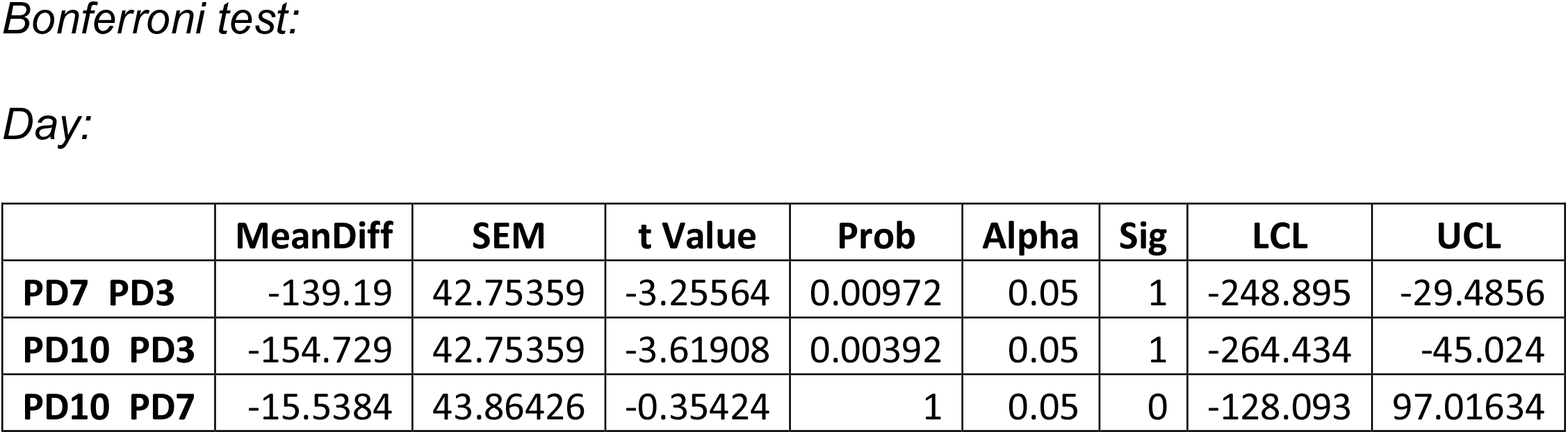
One-way ANOVA of mean proximity in BALB/cJ and SWR/J mice.

**S. Table 26:**
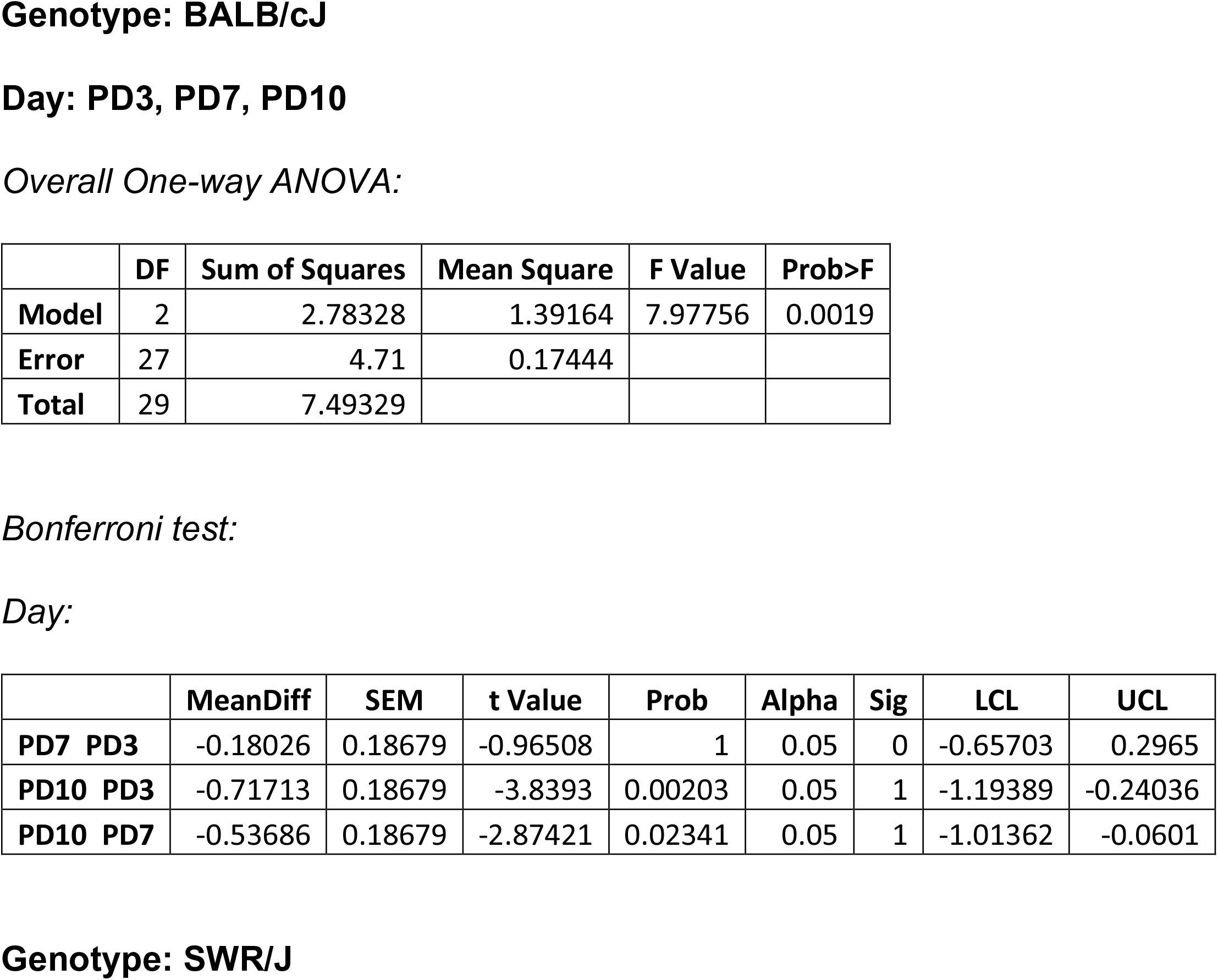
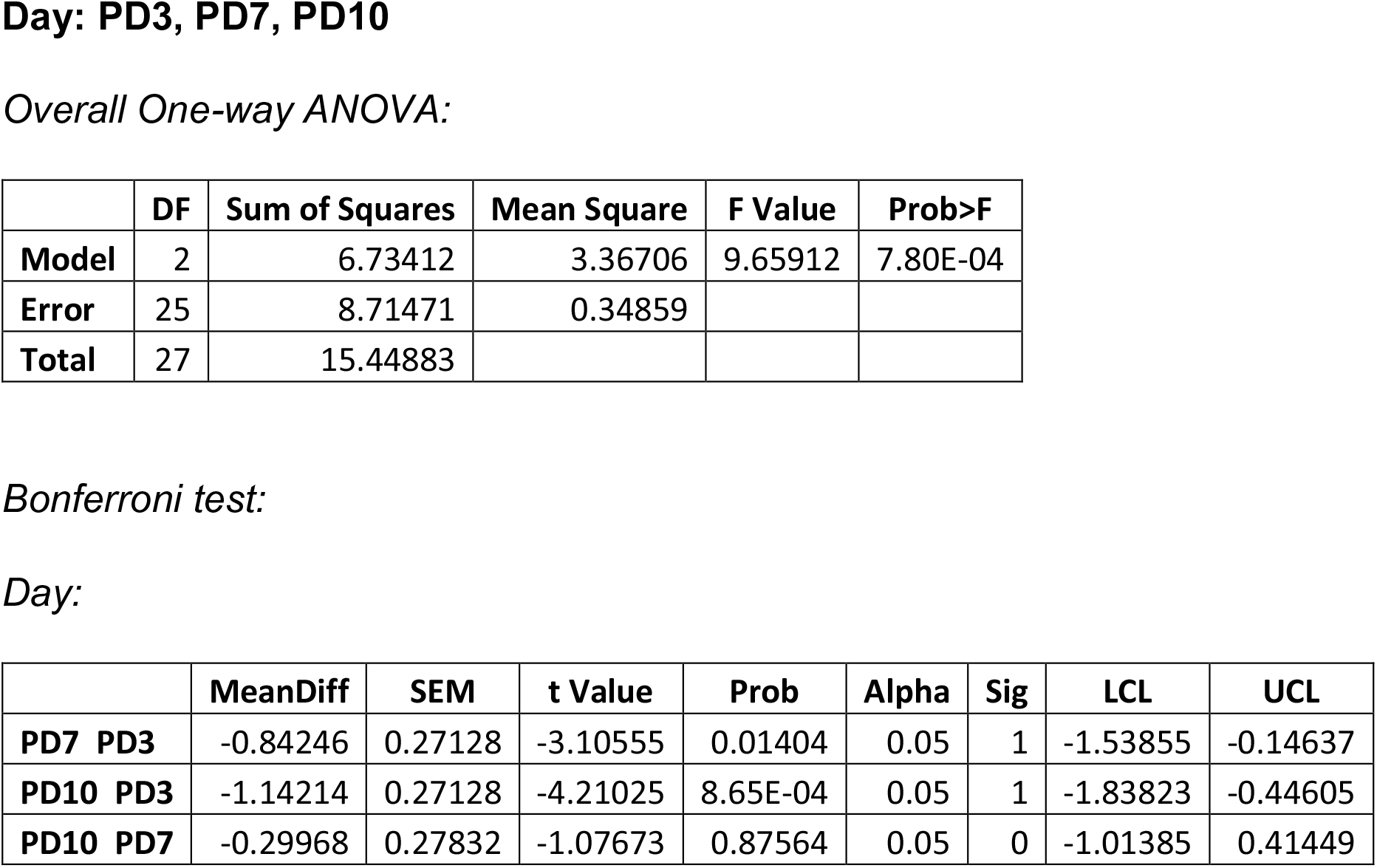
One-way ANOVA of entropy in BALB/cJ and SWR/J mice.

|  | DF | Sum of Squares | Mean Square | F Value | Prob>F |
| --- | --- | --- | --- | --- | --- |
| Model | 2 | 2.78328 | 1.39164 | 7.97756 | 0.0019 |
| Error | 27 | 4.71 | 0.17444 |  |  |
| Total | 29 | 7.49329 |  |  |  |

*Bonferroni test:* *Day:*
|  | MeanDiff | SEM | t Value | Prob | Alpha | Sig | LCL | UCL |
| --- | --- | --- | --- | --- | --- | --- | --- | --- |
| PD7 PD3 | -0.18026 | 0.18679 | -0.96508 | 1 | 0.05 | 0 | -0.65703 | 0.2965 |
| PD10 PD3 | -0.71713 | 0.18679 | -3.8393 | 0.00203 | 0.05 | 1 | -1.19389 | -0.24036 |
| PD10 PD7 | -0.53686 | 0.18679 | -2.87421 | 0.02341 | 0.05 | 1 | -1.01362 | -0.0601 |
| **Model** | 2 | 6.73412 | 3.36706 | 9.65912 | 7.80E-04 |
| **Error** | 25 | 8.71471 | 0.34859 |  |  |
| **Total** | 27 | 15.44883 |  |  |  |

*Bonferroni test:* *Day:*
|  | MeanDiff | SEM | t Value | Prob | Alpha | Sig | LCL | UCL |
| --- | --- | --- | --- | --- | --- | --- | --- | --- |
| **PD7 PD3** | -0.84246 | 0.27128 | -3.10555 | 0.01404 | 0.05 | 1 | -1.53855 | -0.14637 |
| **PD10 PD3** | -1.14214 | 0.27128 | -4.21025 | 8.65E-04 | 0.05 | 1 | -1.83823 | -0.44605 |
| **PD10 PD7** | -0.29968 | 0.27832 | -1.07673 | 0.87564 | 0.05 | 0 | -1.01385 | 0.41449 |

**S. Table 27:** Two-way ANOVA of mean proximity in *Ptpn11* +/+ and *Ptpn11* D61G/+ mice.

|  | DF | Sum of Squares | Mean Square | F Value | P Value |
| --- | --- | --- | --- | --- | --- |
| **Genotype** | 1 | 3977.162 | 3977.162 | 15.965 | 1.30E-04 |
| **Day** | 3 | 10843.43 | 3614.476 | 14.50912 | 8.15E-08 |
| **Interaction** | 3 | 430.2255 | 143.4085 | 0.57567 | 6.32E-01 |
| **Model** | 7 | 16604.09 | 2372.014 | 9.52166 | 7.59E-09 |
| **Error** | 92 | 22918.82 | 249.1176 |  |  |
| **Corrected Total** | 99 | 39522.92 |  |  |  |

Bonferroni test: Genotype:
|  | MeanDiff | SEM | t Value | Prob | Alpha | Sig | LCL | UCL |
| --- | --- | --- | --- | --- | --- | --- | --- | --- |
| *Ptpn11* D61G/+ *Ptpn11* +/+ | 12.87304 | 3.22179 | 3.99562 | 1.30E-04 | 0.05 | 1 | 6.47429 | 19.27178 |

Day:
|  | MeanDiff | SEM | t Value | Prob | Alpha | Sig | LCL | UCL |
| --- | --- | --- | --- | --- | --- | --- | --- | --- |
| PD3 D1 | -18.1161 | 4.46424 | -4.05804 | 6.22E-04 | 0.05 | 1 | -30.1539 | -6.07824 |
| PD5 D1 | -22.9548 | 4.46424 | -5.14194 | 9.15E-06 | 0.05 | 1 | -34.9927 | -10.917 |
| PD5 PD3 | -4.83877 | 4.46424 | -1.0839 | 1.00E+00 | 0.05 | 0 | -16.8766 | 7.19906 |
| PD7 D1 | -29.8159 | 4.46424 | -6.67882 | 1.08E-08 | 0.05 | 1 | -41.8537 | -17.778 |
| PD7 PD3 | -11.6998 | 4.46424 | -2.62078 | 6.16E-02 | 0.05 | 0 | -23.7376 | 0.33805 |
| PD7 PD5 | -6.86101 | 4.46424 | -1.53688 | 0.76652 | 0.05 | 0 | -18.8988 | 5.17682 |

**Genotype: *Ptpn11*+/+ and *Ptpn11* D61G/+**

**S. Table 28:** Two-way ANOVA of entropy in Ptpn11 +/+ and Ptpn11 D61G/+ mice.

|  | DF | Sum of Squares | Mean Square | F Value | P Value |
| --- | --- | --- | --- | --- | --- |
| Genotype | 1 | 1.3485 | 1.3485 | 3.83195 | 5.33E-02 |
| Day | 3 | 17.04024 | 5.68008 | 16.1408 | 1.65E-08 |
| Interaction | 3 | 0.15153 | 0.05051 | 0.14353 | 9.34E-01 |
| Model | 7 | 19.88981 | 2.8414 | 8.07427 | 1.25E-07 |
| Error | 92 | 32.37557 | 0.35191 |  |  |
| Corrected Total | 99 | 52.26538 |  |  |  |

Bonferroni test: Day:
|  | MeanDiff | SEM | t Value | Prob | Alpha | Sig | LCL | UCL |
| --- | --- | --- | --- | --- | --- | --- | --- | --- |
| PD3 D1 | -0.65231 | 0.16779 | -3.88774 | 1.15E-03 | 0.05 | 1 | -1.10475 | -0.19987 |
| PD5 D1 | -0.90658 | 0.16779 | -5.40316 | 3.07E-06 | 0.05 | 1 | -1.35902 | -0.45414 |
| PD5 PD3 | -0.25427 | 0.16779 | -1.51543 | 7.99E-01 | 0.05 | 0 | -0.70671 | 0.19817 |
| PD7 D1 | -1.15023 | 0.16779 | -6.85526 | 4.79E-09 | 0.05 | 1 | -1.60267 | -0.69779 |
| PD7 PD3 | -0.49791 | 0.16779 | -2.96752 | 2.30E-02 | 0.05 | 1 | -0.95035 | -0.04547 |
| PD7 PD5 | -0.24364 | 0.16779 | -1.4521 | 0.89926 | 0.05 | 0 | -0.69608 | 0.2088 |

Overall Two-way ANOVA:

**S. Table 29:** Two-way ANOVA of mean proximity in *Ptpn11* +/+ and *Ptpn11* N308D/+ mice.

|  | DF | Sum of Squares | Mean Square | F Value | P Value |
| --- | --- | --- | --- | --- | --- |
| Genotype | 1 | 84.06739 | 84.06739 | 0.35415 | 5.55E-01 |
| Day | 1 | 1.79816 | 1.79816 | 0.00757 | 9.31E-01 |
| Interaction | 1 | 505.3925 | 505.3925 | 2.12904 | 1.53E-01 |
| Model | 3 | 589.5332 | 196.5111 | 0.82783 | 4.87E-01 |
| Error | 38 | 9020.475 | 237.3809 |  |  |
| Corrected Total | 41 | 9610.008 |  |  |  |

Genotype: *Ptpn11*+/+ and *Ptpn11* N308D/+
Treatment: Veh, SL327
*Overall Two-way ANOVA:*

**S. Table 30:**
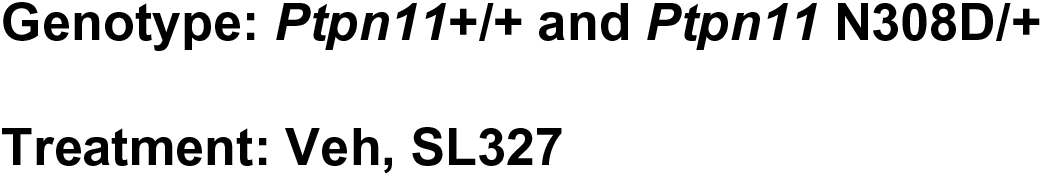
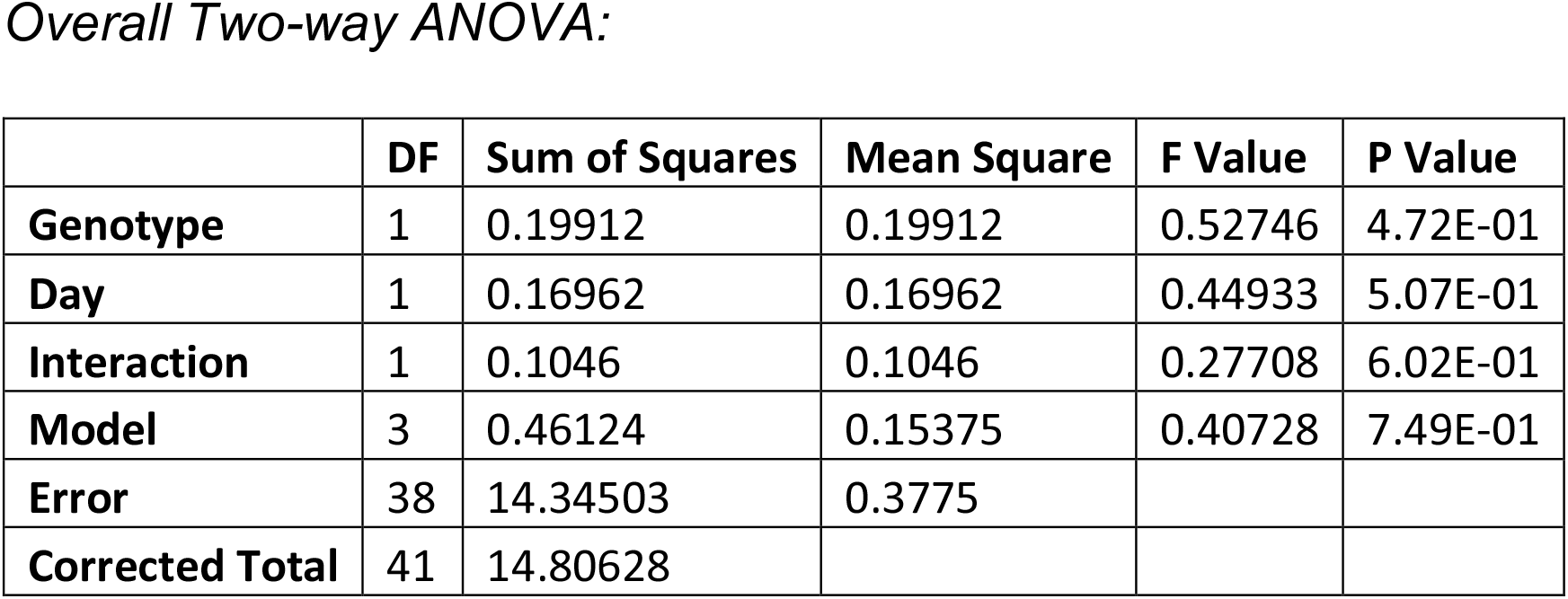
Two-way ANOVA of entropy in *Ptpn11* +/+ and *Ptpn11* N308D/+ mice.

|  | DF | Sum of Squares | Mean Square | F Value | P Value |
| --- | --- | --- | --- | --- | --- |
| **Genotype** | 1 | 0.19912 | 0.19912 | 0.52746 | 4.72E-01 |
| **Day** | 1 | 0.16962 | 0.16962 | 0.44933 | 5.07E-01 |
| **Interaction** | 1 | 0.1046 | 0.1046 | 0.27708 | 6.02E-01 |
| **Model** | 3 | 0.46124 | 0.15375 | 0.40728 | 7.49E-01 |
| **Error** | 38 | 14.34503 | 0.3775 |  |  |
| **Corrected Total** | 41 | 14.80628 |  |  |  |

**S. Table 31:** Two-way ANOVA of mean proximity in C57Bl/6J and DBA/2J mice.

|  | DF | Sum of Squares | Mean Square | F Value | P Value |
| --- | --- | --- | --- | --- | --- |
| **Genotype** | 1 | 1039.771 | 1039.771 | 3.60823 | 6.96E-02 |
| **Day** | 2 | 431.4297 | 215.7148 | 0.74858 | 4.84E-01 |
| **Interaction** | 2 | 1885.235 | 942.6175 | 3.27108 | 5.54E-02 |
| **Model** | 5 | 3356.436 | 671.2872 | 2.32951 | 7.37E-02 |
| **Error** | 24 | 6916.003 | 288.1668 |  |  |
| **Corrected Total** | 29 | 10272.44 |  |  |  |

**S. Table 32:** Two-way ANOVA of entropy in C57Bl/6J and DBA/2J mice.

|  | DF | Sum of Squares | Mean Square | F Value | P Value |
| --- | --- | --- | --- | --- | --- |
| **Genotype** | 1 | 1.39241 | 1.39241 | 4.44148 | 4.57E-02 |
| **Day** | 2 | 6.87801 | 3.43901 | 10.96963 | 4.13E-04 |
| **Interaction** | 2 | 0.49123 | 0.24562 | 0.78346 | 4.68E-01 |
| **Model** | 5 | 8.76166 | 1.75233 | 5.58953 | 1.49E-03 |
| **Error** | 24 | 7.52406 | 0.3135 |  |  |
| **Corrected Total** | 29 | 16.28572 |  |  |  |

*Bonferroni test:* *Genotype:*
|  | MeanDiff | SEM | t Value | Prob | Alpha | Sig | LCL | UCL |
| --- | --- | --- | --- | --- | --- | --- | --- | --- |
| **DBA/2J C57Bl/6J** | 0.43088 | 0.20445 | 2.10748 | 4.57E-02 | 0.05 | 1 | 0.00891 | 0.85284 |

*Peaks:*
|  | MeanDiff | SEM | t Value | Prob | Alpha | Sig | LCL | UCL |
| --- | --- | --- | --- | --- | --- | --- | --- | --- |
| **PD5(NE) PD3** | 1.13995 | 0.2504 | 4.55251 | 3.89E-04 | 0.05 | 1 | 0.49551 | 1.78439 |
| **PD5(SW) PD3** | 0.8089 | 0.2504 | 3.23043 | 0.0107 | 0.05 | 1 | 0.16446 | 1.45334 |
| **PD5(SW) PD5(NE)** | -0.33105 | 0.2504 | -1.32208 | 0.59581 | 0.05 | 0 | -0.97549 | 0.31339 |

